# Polygenic basis of incipient reproductive isolation in hybridizing populations is revealed by pangenomic and epigenetic divergence

**DOI:** 10.1101/2025.10.11.681778

**Authors:** A. A. Ruggieri, F. Cicconardi, N. Bellin, S. H. Montgomery, J. Mallet, S. M. Van Belleghem, W. O. McMillan, B. A. Counterman, R. Papa

## Abstract

Incipient reproductive isolation in the presence of gene flow has traditionally been attributed to a small number of major-effect loci under strong selection, representing only a minor fraction of the genome. Using the *Heliconius erato* adaptive radiation—a butterfly species with populations at varying levels of genetic differentiation—we employed a pan-genome approach to investigate genome-wide mutational divergence, epigenetic changes, and structural variation contributing to divergence. In contrast to earlier studies that identified only a few highly divergent loci, our high-resolution analysis reveals widespread divergence across the genome, suggesting polygenic barriers to gene flow. Epigenetic divergence assessed using ATAC-seq, highlights population-specific differences in chromatin accessibility, which may reflect SNPs distribution or precede genetic differentiation by reshaping regulatory landscapes. We also identified new structural variants, including polymorphic indels in open chromatin, that further associate with genomic divergence. Together, these findings show that the genomic landscape of *H. erato* is shaped by a dynamic interplay of mutational changes, epigenetic modifications, and structural variation. We emphasize the role of developmental, behavioral, and ecological adaptations and provide a broader perspective on the functional genetic basis of genome-wide divergence in *Heliconius*. This emerging polygenic framework offers a more detailed understanding of how reproductive barriers evolve under ongoing gene flow.

## INTRODUCTION

The concept of species has shifted over time—from a focus on adaptive divergence to emphasizing genetic incompatibilities that maintain species boundaries. This change highlights population-level genetic divergence as the foundation of speciation, with the notable exception of hybrid speciation, where new species arise through hybridization between distinct parental lineages (Mallet, 2007; Abbott et al., 2013). Incipient reproductive isolation—the early evolution of barriers that reduce or prevent gene flow—begins with the accumulation of genetic differences at specific barrier loci. These differences can increase over time, reinforcing isolation, often through selection on morphological, reproductive, behavioral, or ecological traits—commonly referred to as “speciation genes” (Wu, 2001; Feder & Nosil, 2012; Sakamoto et al., 2019).

Population genomics has been central to identifying barrier loci and understanding how they persist during early speciation. A promising approach involves studying populations that exhibit local adaptation and phenotypic divergence despite ongoing gene flow, especially in sympatric and parapatric contexts (Via, 2012; Feulner et al., 2015; Mérot et al., 2020; Van Belleghem et al., 2021). In such cases, divergence tends to be concentrated at loci under strong selection, forming “genomic islands of speciation”—regions of elevated differentiation that emerge when divergent selection counteracts gene flow (Nosil & Feder, 2012). Over time, hitchhiking and linkage may allow divergence to spread, reinforcing isolation across broader genomic regions.

While GWAS and fine-scale mapping have improved our understanding of these barriers, major gaps remain, particularly in identifying the number and functional roles of contributing loci. Structural variants (SVs), particularly small-scale variants, may be underappreciated drivers of regulatory divergence and speciation, especially given the difficulty of detecting non-coding regulatory elements in non-model organisms (Monaghan et al., 2020). However, recent advances—such as chromatin accessibility assays (e.g., ATAC-seq; Buenrostro et al., 2015), population-specific genome assemblies, and improved computational tools—are beginning to close these gaps.

Hybrid zones, where diverging populations interbreed and produce intermediate phenotypes, provide powerful natural laboratories for studying early genomic divergence (Barton & Hewitt, 1989; Velo-Antón, 2021). The *Heliconius* genus, with over 40 species and hundreds of divergent mimetic subspecies across diverse hybrid zones and varying levels of reproductive isolation, is a leading model for studying the genetics of incipient speciation. Prior work has shown that introgression, selection, and assortative mating contribute to barriers between populations (Edelman et al., 2019; Martin et al., 2019; Rosser et al., 2024). In *H. erato*, population genetic studies have suggested that divergence is largely limited to a few color pattern loci (Van Belleghem et al., 2017; Kronforst & Papa, 2015; Heliconius Genome Consortium, 2012). While these genes are critical for mimicry, predator avoidance, and mating, it is unlikely that the rest of the genome remains completely homogeneous, especially given known behavioral, metabolic, and ecological adaptations (Montejo-Kovacevich et al., 2022).

In this study, we extend the analysis of genome-wide divergence by integrating de novo genome assemblies, chromatin accessibility data, and improved gene annotations across *H. erato* populations with varying levels of geographic isolation. Our findings reveal a multilocus architecture—spanning both genetic and epigenetic variation—that highlights the complexity of incipient reproductive isolation. This polygenic framework suggests that genome-wide divergence emerges earlier and more broadly than previously assumed, challenging the classical view of speciation driven by a few isolated genomic islands.

## RESULTS

### Genome assemblies, pan-genome alignment, and lineage-specific sequence composition

We generated new genome assemblies for five *Heliconius erato* populations—*H. e. notabilis, H. e. etylus, H. e. hydara, H. e. chestertonii*, and *H. e. favorinus*—and integrated them with the previously published *H. e. demophoon* genome to construct a comprehensive pan-genome (Fig. 1A,C). The genome of *H. e. demophoon* spans 377.2 Mb. The other genomes exhibited limited variation in total size, remaining within 13% of the *H. e. demophoon* genome. Among them, *H. e. chestertonii* had the smallest genome at 326.2 Mb. The genome sizes of *H. e. hydara, H. e. favorinus, H. e. etylus*, and *H. e. notabilis* were 356.5 Mb, 363.7 Mb, 336.7 Mb, and 338.4 Mb, respectively (Fig 1B; Table S1a). Altogether, the assembled pan-genome spans 688 Mb and was used as a reference to align resequencing data as well as ATAC-seq data (Fig 1B).

**Figure 1:**
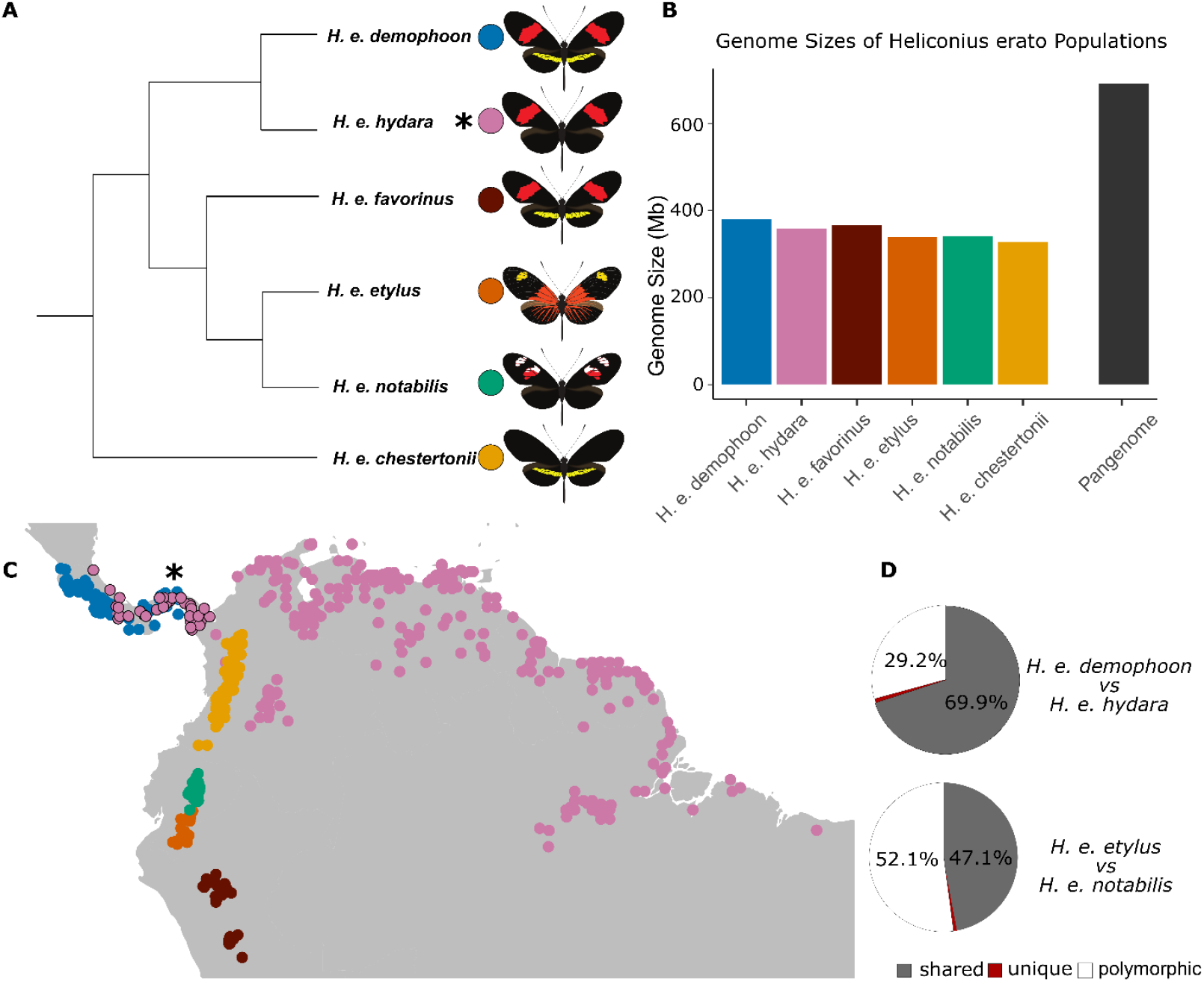
Geographic Distribution, Phylogenetic Relationship of Geographical Populations, and Pan-Genome Overview. **(A)** Dendrogram showing the phylogenetic relationships among the geographical mimetic populations of *Heliconius erato* used in this study. **(B)** Pan-genome overview generated from the six geographical populations. Gray represents the percentage of the total sequence length shared among all six populations, black represents sequences not exclusive to one morph but not shared among all six, and other colors indicate sequences exclusive to each population. **(C)** Geographic distribution of the populations based on sampling data obtained from [Heliconius Maps](https://heliconius-maps.github.io/) (Rosser & Mallet 2024). The asterisk shows where the specific samples of *H. e. hydara* used in this study were sampled from. **(D)** Distribution of ATAC-seq peaks in forewing between adjacent populations. “Shared” indicates peaks consistently present in both populations, “polymorphic” indicates peaks not consistently present in all samples of a population, and “unique” indicates peaks identified only in one population.

### Genes under selective forces between adjacent subspecies

We applied a new approach to distinguish potential *F*_*st*_ signals of divergence from background noise by identifying regions with an elevated density of points above the genome-wide average, in addition to the highest *F*_*st*_ values. This method allows us to detect regions where *F*_*st*_ is consistently high, minimizing the influence of random outliers—even when calculated using small 1 kb windows. Using this approach, we identified 91 regions with elevated *F*_*st*_ in the *demophoon–hydara* comparison (covering 2.943 Mb, or 0.427% of the pan-genome) and 76 regions in *etylus–notabilis* (ranging from 53,000 to 280,000 bp, covering 2.916 Mb, or 0.423% of the pan-genome; Fig. 2A).

**Figure 2:**
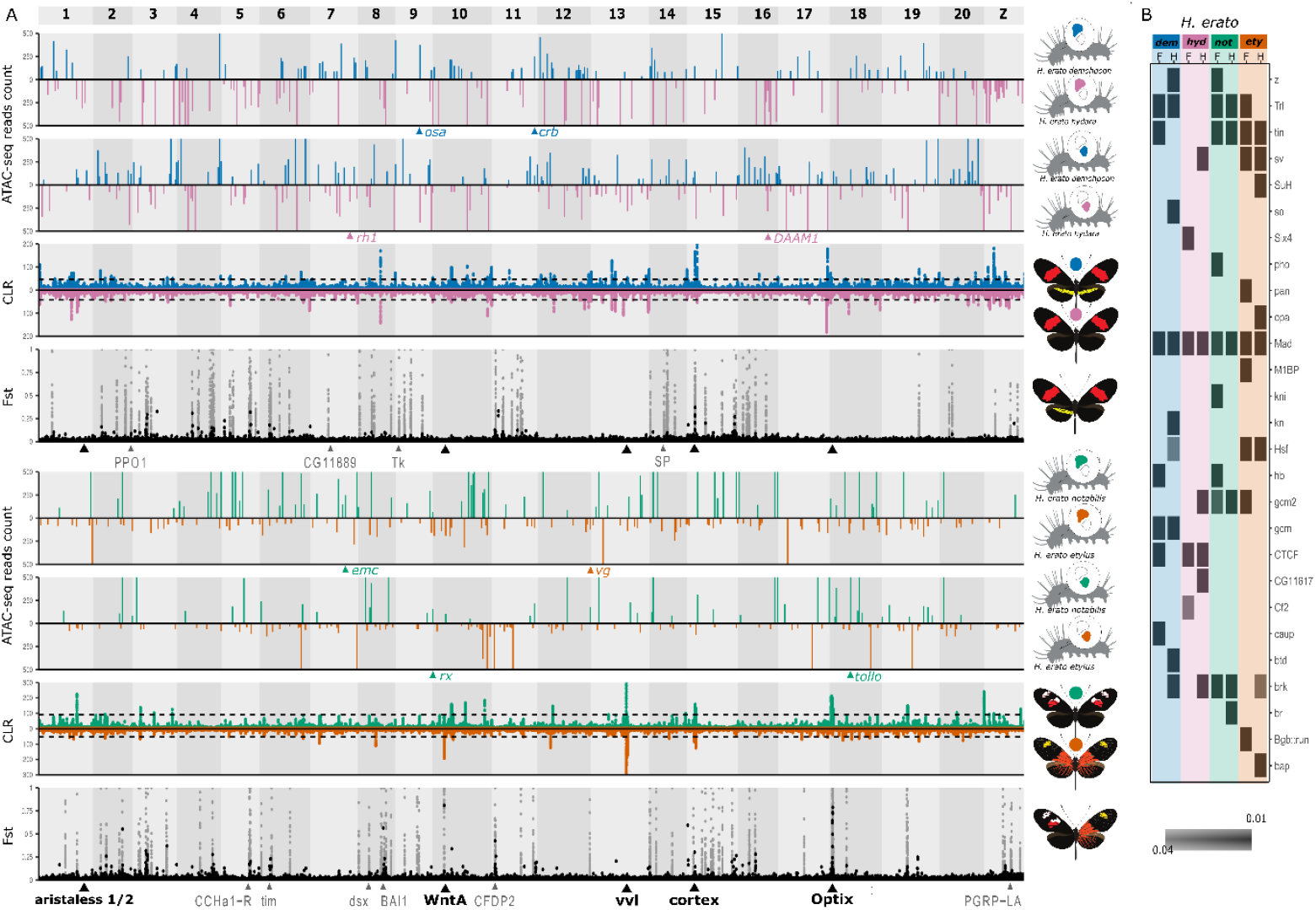
Genome-Wide Analysis of Genetic and Epigenetic Data from Two Adjacent populations. **(A)** The first four tracks show comparisons between *H. e. demophoon* and *H. e. hydara*, while the last four tracks compare *H. e. notabilis* with *H. e. etylus*. The two ATAC-seq tracks (y-axis = ATAC-seq read count) indicate differences in chromatin dynamics between the forewing and hindwing, focusing on ATAC peaks that are unique to one population or the other. The genome was divided into 1 kb non-overlapping windows, and for each window, the average peak read count in *H. e. hydara* was subtracted from that in *H. e. demophoon*. Results are displayed in pink (lower half) if chromatin activity was higher in *H. e. hydara* and in blue (upper half) if activity was higher in *H. e. demophoon*. The third track (y-axis = CLR) represents selective sweep signals between the two populations. The fourth track (y-axis = *F*_*st*_) shows *F*_*st*_ values from population comparisons, with black dots indicating *F*_*st*_ values calculated in 50 kb windows and gray dots indicating significant *F*_*st*_ signals in 1 kb windows. The 1 kb *F*_*st*_ background signal is omitted for clarity. These analyses are repeated for the comparison between *H. e. notabilis* and *H. e. etylus*, with results displayed in green and orange. The black triangle shows the classic color pattern gene’s position, the grey ones represent examples of putative new involved genes from the fst, and the different colored triangles on the ATAC-seq track show putative new genes from the ATAC-seq signal **(B)** Transcription factor binding site enrichment analysis for each wing and each adjacent population.

These regions also exhibit elevated *D*_*xy*_ compared to the genomic background—even relative to all high-*F*_*st*_ points (Fig. S6). The co-occurrence of high *F*_*st*_ and high *D*_*xy*_ suggests resistance to gene flow (Cruickshank & Hahn, 2014), indicating that these regions are likely not introgressing, possibly due to selection maintaining their divergence.

Within these regions, we identified 54 genes in *demophoon–hydara* and 88 in *etylus–notabilis*, including the well-known wing color pattern genes: *WntA* and *optix* in *etylus–notabilis*, and *cortex* in *demophoon–hydara* (Table S2). We also found strong divergence in *doublesex*, a gene associated with sexual dimorphism and color pattern variation in *Papilio* swallowtail and *Coliadinae* butterflies (VanKuren et al., 2022; Rodriguez-Caro et al. 2021). In addition, four genes linked to phenoloxidases (PO)—key components of the insect immune system—were detected (González-Santoyo et al., 2012; Marieshwari et al., 2023; Table S2).

We further identified genes involved in chromatin modification and transcription regulation, including *Chromodomain-helicase-DNA-binding protein 7* and *Integrator complex subunit 7*, suggesting a potential role for chromatin dynamics in divergence. These genes are implicated in chromatin remodeling and defense against environmental stressors and may underlie adaptive differences between populations. Additionally, the *timeless* gene is a cornerstone of circadian rhythm, affecting daily activity and mating timing, especially in *Drosophila* and mosquito species (Rosato et al., 2006). *Chitin synthase chs-2* plays a central role in forming the insect exoskeleton, influencing development and desiccation resistance — traits under strong environmental selection (Moussian, 2010). Moreover, genes involved in chemosensation and reproduction — like *gustatory receptor 64f-like*, and *seminal fluid proteins (HACP027, HACP057, HACP011)* — are fundamental in mate recognition, post-mating physiology, and sexual selection. These traits often diverge rapidly between populations and species, reinforcing reproductive isolation and speciation processes in insects (Chapman et al., 2003; Shukla & Palli, 2012).Altogether, our results point to a multilocus architecture underlying barriers to gene flow, involving both classic speciation genes and additional loci linked to immunity, regulation, and chromatin structure.

### Chromatin dynamics

In addition to a finer scale *F*_*st*_ analysis, we also investigate diversity and possible barrier loci at the epigenetics level, using ATAC-seq. The number of ATAC-seq peaks identified per population ranged from 13,165 in *H. e. notabilis* to 22,804 in *H. e. chestertonii*. Across all populations, a total of 26,291 peaks were detected, of which 57.2% (15,036) were consistently found in all populations (shared ATAC-peaks). Peaks unique to a single population (unique ATAC-peaks) made up only 1.26% (332), ranging from 0.03% (4) in *H. e. notabilis* to 0.68% (156) in *H. e. chestertonii* (Table S1a–b). These low numbers are likely due to our highly conservative criterion, where a peak was considered unique to one population only if it was present in every individual from that population and absent from all others. The remaining 41.54% consisted of polymorphic ATAC-peaks, present in more than one population but not all (Fig. 1D). These results show that the studied populations maintain an invariant core of ATAC-seq peaks while exhibiting a high number of polymorphic regions, with completely unique ATAC-seq peaks being relatively low.

Analysis of chromatin accessibility differences between adjacent subspecies revealed that across the *H. e. demophoon* and *H. e. hydara* hybrid zone, 51.4% to 74.9% (7,696-11,003) of the ATAC-seq peaks are shared (see Table S4). Only a small fraction, less than 1% (ranging from 117 to 161 ATAC-peaks), are completely unique to one geographical subspecies, meaning these peaks are found in all samples of one population but absent in all other populations (Table S4). This indicates that between 29.2% and 47.5% (4,594-7,121) of the ATAC-seq peaks are polymorphic between two populations, appearing in at least one sample but not in all of both populations. Similarly, in the *H. e. notabilis* and *H. e. etylus* adjacent populations, between 38% and 52% (5997-7871) ATAC-seq peaks are shared, less than 1% (50-138) of peaks are unique, and between 46,63% and 60,91% (6937-9535) are variables (Table S4).

Our subsequent analyses focused on the set of unique ATAC-seq peaks identified between adjacent populations, which represent fixed binary differences between populations. These peaks may correspond to particularly relevant regulatory elements as they are the most divergent between adjacent populations. To explore the potential mechanisms underlying this regulatory divergence, we searched for transcription factors (TFs) that might bind to these regions. We performed TF binding site enrichment analysis on these ATAC-seq peaks and identified several potentially important TFs (Fig. 2B). The gene *Mothers against dpp* (*Mad*) is related to wing development in *H. melpomen*e (Baxter et al 2010). *Mad* was also previously identified as a possible gene involved in chromatin divergence between *H. erato, H. charithonia*, and *H. melpomene* (Ruggieri et al 2022). Several more TFs may be relevant for our study, including *sine oculis* (*so*), crucial for the formation of compound eyes in *Drosophila* (Serikak & O’Tousa 1994). Further, *Suppressor of Hairless* (*Su(H)*) operates within the Notch signaling pathway, which is fundamental for regulating cell differentiation, proliferation, and apoptosis during wing development in *Drosophila* (Klein et al. 2000). A notable gene includes *pangolin* (*pan*), involved in segment polarity through the *Wnt* signaling pathway (Brunner et al. 1997). It has been previously found in *H. erato* and *H. melpomene* as a possible gene involved in wing development (Van Belleghem et al. 2023). Because these genes are active at different stages of development, they may contribute to generating a cascade effect that would explain the genome-wide chromatin activity divergence between populations.

We identified the nearest genes to each unique ATAC-seq peak as a proxy for potential target loci, revealing a set of candidate genes that may be under positive selection (Table S5; Table 1). Across the *demophoon-hydara* hybrid zone, 344 genes were identified, while 237 in *etylus-notabilis*. Across both datasets, a total of 581 genes were annotated, of which 40 exhibit functions potentially associated with divergent evolution in these butterflies. These genes were classified into six functional categories: Circadian Rhythm and Behavioral Timing, Cuticle Formation and Pigmentation, Developmental Patterning and Morphogenesis, Hormone Metabolism and Detoxification, Neurotransmission and Sensory Perception, and Reproductive and Post-Mating Response (Table 1).

**Table 1:**
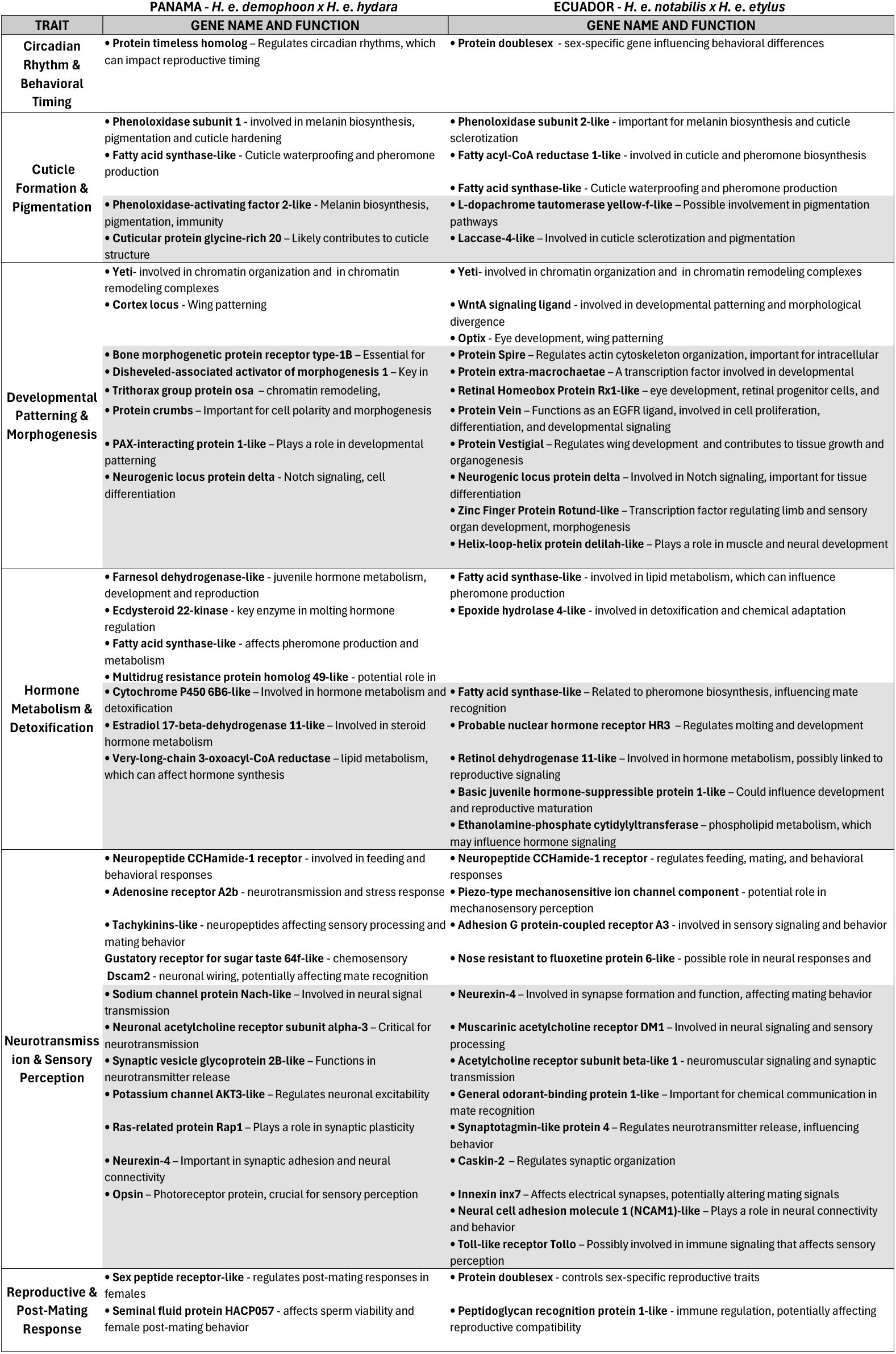
Relevant genes under ATAC-seq or *F*_*st*_ signal. Table reporting identified relevant genes, general traits they participate in, gene name, a short description of their functions, hybrid zone they have been identified with (Panama: *H. e. demophoon x H. e. hydara*; Ecuador: *H. e. notabilis x H. e. etylus*), and the type of data the genes have been identified with, in white are the genes identified with Fst, and in gray the ones identified with ATAC-seq.

In addition to fine-scale *F*_*st*_ analysis, we investigated potential barrier loci at the epigenetic level using ATAC-seq. The number of ATAC-seq peaks per population ranged from 13,165 in *H. e. notabilis* to 22,804 in *H. e. chestertonii*. Across all populations, we identified 26,291 peaks, of which 57.2% (15,036) were shared across all populations. Unique ATAC-seq peaks—present in all individuals of one population and absent from all others—accounted for only 1.26% (332 peaks), ranging from just 4 in

*H. e. notabilis* to 156 in *H. e. chestertonii* (Table S1a–b). These low numbers reflect our highly conservative criteria, where a peak was considered unique to one population only if it was present in every individual from that population and absent from all others. The remaining 41.54% of peaks were polymorphic, present in more than one but not all populations (Fig. 1D). These results suggest that while subspecies share a core set of regulatory elements, substantial variation exists at polymorphic sites, with truly unique peaks being rare.

Across hybrid zones, we observed similar patterns. In the *H. e. demophoon* × *H. e. hydara* zone, 51.4– 74.9% (7,696–11,003) of peaks were shared, less than 1% (117–161) were unique, and 29.2–47.5% (4,594–7,121) were polymorphic (Table S4). In *H. e. notabilis* × *H. e. etylus*, 38–52% (5,997–7,871) were shared, <1% (50–138) were unique, and 46.6–60.9% (6,937–9,535) were polymorphic. We focused on the small set of unique ATAC-seq peaks between each pair of adjacent populations, as these represent fixed, binary regulatory differences between populations and may be especially relevant to divergence.

To identify potential transcription factors (TFs) that may bind to the unique ATAC-seq peaks—and thereby shed light on which regulatory pathways might be diverging—we performed TF binding site enrichment analysis on these regions (Fig. 2B). Several TFs were identified as candidates: *Mothers against dpp* (Mad), involved in wing development in *H. melpomene* (Baxter et al., 2010) and previously linked to chromatin divergence (Ruggieri et al., 2022); *sine oculis* (*so*), required for compound eye development in *Drosophila* (Serikak & O’Tousa, 1994); *Suppressor of Hairless* (*Su(H)*), part of the Notch signaling pathway involved in wing morphogenesis (Klein et al., 2000); and *pangolin* (*pan*), associated with Wnt signaling and segment polarity, also previously implicated in wing development in *H. erato* and *H. melpomene* (Brunner et al., 1997; Van Belleghem et al., 2023). Because these TFs act at different developmental stages, they may contribute to a regulatory cascade that drives genome-wide chromatin divergence between populations.

To gain insight into the potential downstream effects of regulatory divergence, we identified the nearest genes to each unique ATAC-seq peak as putative target loci. This revealed 344 candidate genes in the *demophoon–hydara* comparison and 237 in *etylus–notabilis*. In total, 581 genes were annotated, some of which have putative roles in divergence. These were grouped into six functional categories: Circadian Rhythm and Behavioral Timing, Cuticle Formation and Pigmentation, Developmental Patterning and Morphogenesis, Hormone Metabolism and Detoxification, Neurotransmission and Sensory Perception, and Reproductive and Post-Mating Response (Table 1; Table S5). Genes like *fatty acid synthase* and *acyl-CoA reductase* facilitate cuticle waterproofing and pheromone production, essential for desiccation resistance and intraspecific communication (Blomquist & Vogt, 2003). Pigmentation-related enzymes, including *phenoloxidase subunits* and *yellow-like* genes, are involved in melanin synthesis, which plays dual roles in immune defense and visual signaling (True, 2003). Additionally, regulators like *Disheveled* and *Protein delta*, part of the Wnt and Notch signaling pathways, mediate tissue patterning and morphogenesis (Morey 2024). Detoxification genes such as *cytochrome P450s* support metabolic resilience in diverse habitats by breaking down xenobiotics (Feyereisen, 2012), while *CCHamide receptors* and *guanylate cyclase* control feeding, foraging, and locomotor responses (Vermehren et al 2006).

### Correlation between chromatin differences and *F*_*st*_ across levels of isolation

We hypothesized that the observed differences in chromatin activity may reflect or potentially drive the onset of genetic changes. Therefore, we expected to find an *F*_*st*_ or selective sweep signal in the open chromatin regions represented by ATAC-seq. To explore a potential connection between variation in chromatin dynamics and the *F*_*st*_ signal, we employed two distinct testing methods: an initial Kolmogorov–Smirnov (KS) test and a custom test based on the binomial test. The Kolmogorov– Smirnov test checks whether the overall distribution of *F*_*st*_ in a unique ATAC-seq peak is higher than the background noise. This test was not significant across either hybrid zone (*demophoon-hydara*,adj p-value=0.2 and D=0.06 Fig. 4A; *notabilis-etylus* adj p-value=1 and D=0.07, Table S6). The binomial test for a higher probability of having a unique ATAC-seq peak in *F*_*st*_ intervals compared to background noise, testing each interval independently and, therefore, may be better at identifying small differences. This test generated significant results for the *demophoon-hydara* comparison only in the *F*_*st*_ intervals of 0.2-0.3 and 0.6-0.7 (see Fig. 4B). For the *notabilis-etylus* putative hybrid zone, only a single interval was significant in the Custom binomial test (between 0.4-0.5, see Table S6). These results suggest that there may be a connection between *F*_*st*_ and ATAC-seq across hybrid zones, but the signals appear very weak since only one of the tests identified a few significant results. Similarly, these tests were applied to assess the relationship between unique ATAC-seq peaks and selective sweep signals across the four adjacent populations; none of these comparisons yielded statistical significance (Fig. S7).

**Figure 3:**
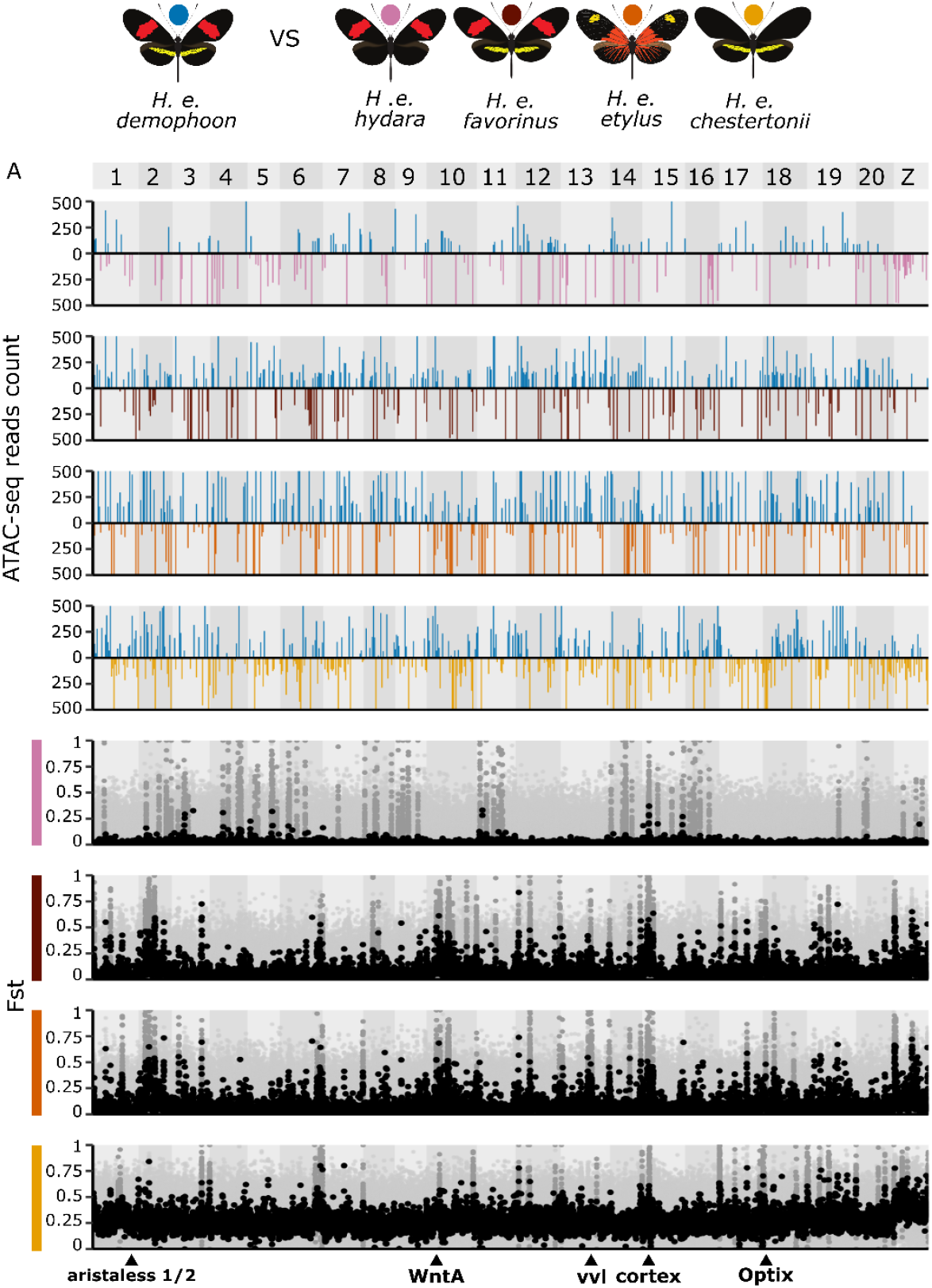
Pairwise Comparisons for H. e. demophoon. Plots representing forewing chromatin dynamic differences and *F*_*st*_ values, as described in Figure 2. From top to bottom, comparisons are shown for H. e. demophoon versus H. e. hydara, H. e. favorinus, H. e. etylus, and H. e. chestertonii. Each *F*_*st*_ plot displays *F*_*st*_ values calculated in 50 kbp windows (black), signals identified from 1 kbp windows using custom analyses to distinguish noise from true signals (dark gray), and background noise from 1 kbp windows (light gray.

**Figure 4:**
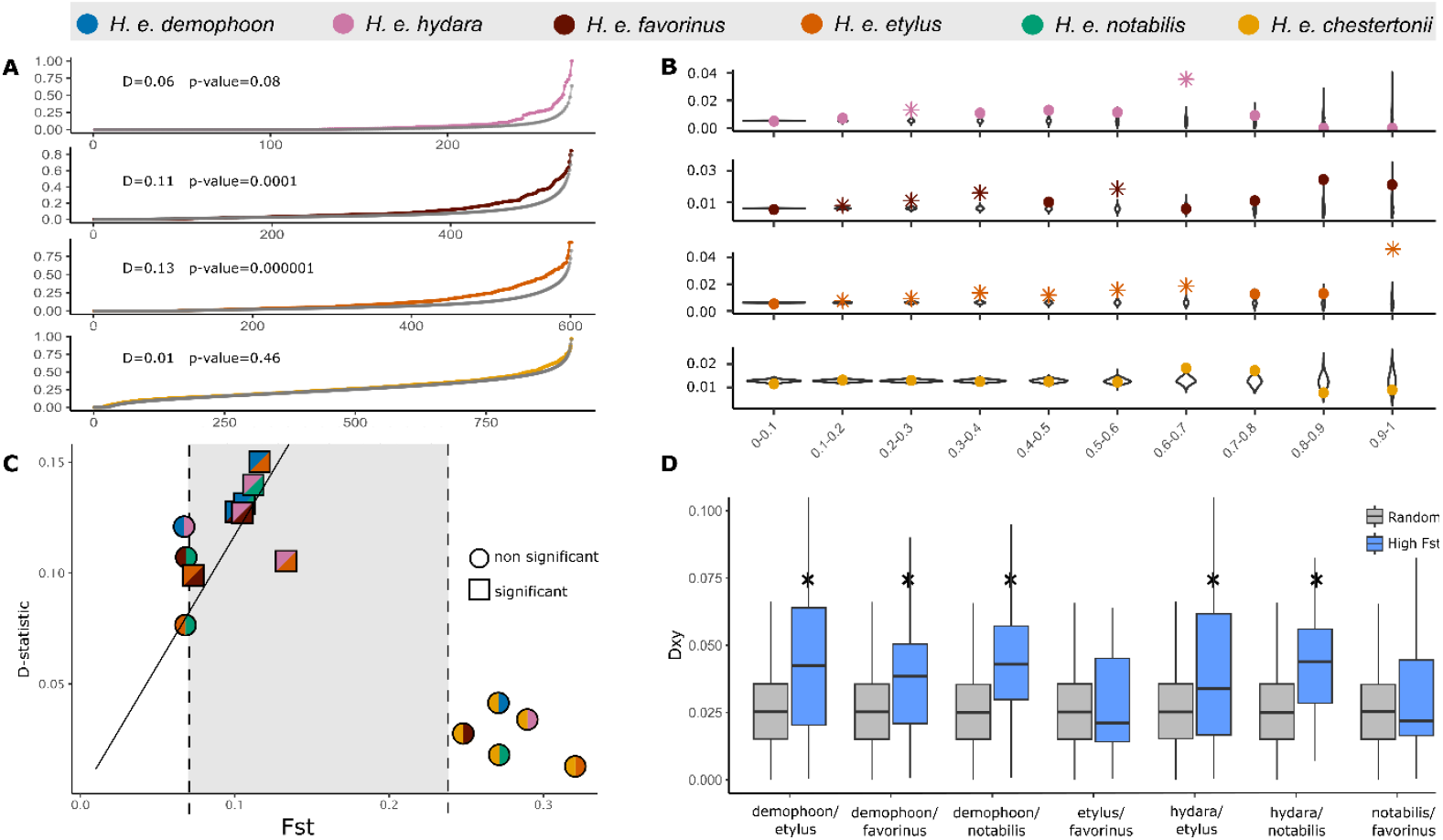
Distance-Related Patterns and Non-Homologous Sequence Dynamics. **(A)** Results from the Ks test for the same comparisons. The D-statistic from the Ks test is reported along with its adjusted p-value. **(B)** Results from the custom binomial test for the same pairwise comparisons. Statistically significant results are indicated by asterisks. **(C)** Scatterplot summarizing the results of Ks tests for all 15 possible pairwise comparisons. The y-axis represents the D-statistic from the Ks test, while the x-axis indicates the average *F*_*st*_ value for each comparison. Statistically significant tests (adjusted p-value < 0.05) are represented by squares, with different colors indicating the populations being compared. **(D)** Boxplot showing Dxy comparisons between unique peaks with high *F*_*st*_ values and random points for each of the significant pairs identified in panel A. Asterisks indicate the significance of the Ks test. Random samples were generated as the ranked average of 1,000 points for each comparison.

Given the hypothesis that a potential *F*_*st*_ signal may emerge in cases of reduced gene flow, we conducted both analyses on non-hybrid zone pairs, given their increasing divergence and geographical distance, with *H. e. demophoon* as the reference (comparing *H. e. demophoon* vs. *H. e. favorinus, H. e. etylus*, and *H. e. chestertonii*). The KS test produced significant results for *H. e. demophoon* vs. *H. e. favorinus* (p-value < 0.01 and D= 0.11) and *H. e. etylus (*p-value < 0.01 and D= 0.13) but no significance when compared to *H. e. chestertonii* (p-value=0.46 and D =0.01; Fig. 4A). Furthermore, the values of D derived from the KS test increased with population divergence until reaching *H. e. chestertonii* (Fig. 4A). The Binomial test also revealed a larger number of significant correlations in *demophoon-favorinus* (a total of 4 intervals) and in *demophoon-etylus* with 4 and 7 intervals respectively, but none only to decline when *H. e. demophoon* was compared to *H. e. chestertonii* (Fig. 4B). The Custom binomial test was also performed using *F*_*st*_ calculated for 50k windows giving results that were comparable with the original test with the exception of *H. e. demophoon vs H. e. chestertonii* where we found 3 significant intervals (0.4-0.5;0.5-0.6;0.7-0.8; Fig S8.). Taken together, these results suggest that there may be a weak increase in *F*_*st*_ in ATAC-seq data between adjacent populations, but it is possibly obscured by gene flow or not small enough genetic interval measured. When gene flow is reduced or absent, the signal becomes more evident and increases with geographic distance.

We conducted the KS test for all possible pairwise comparisons to investigate whether the observed positive correlation pattern persisted in other comparisons (15 total pairs). We extracted the D-value, which represents how much higher the *F*_*st*_ value of the unique ATAC-peaks is compared to the background, and an adjusted p-value for each pair, and then plotted these results against the average *F*_*st*_ value (Fig. 4C). Our analysis shows that the KS test is significant within a certain *F*_*st*_ threshold (between 0.7 and 0.24; Fig. 4C). Within this defined range, we observed a proportional increase in the D-value as the average *F*_*st*_ value increased (F-statistic: 158.1 and p-value: 5.645e-05). Notably, no comparisons involving *H. e. chestertonii* were significant, possibly due to high background divergence. The KS test versus *F*_*st*_ plot was also produced using different datasets to confirm the consistency of the observed patterns. We utilized *F*_*st*_ values calculated for 50 kbp windows instead of 1 kbp windows, and we considered unique ATAC-seq peaks identified in 2 out of 3 samples. This additional testing was performed to ensure that the results for *H. e. chestertonii* were not influenced by the number of ATAC-seq samples (since it is the only one with a total of 2 samples). Notably, all these tests yielded comparable results and exhibited consistent trends (Fig S9).

In population pairs that do not currently hybridize, there is a correlation between genetic differentiation and ATAC-seq signals, which increases with the average *F*_*st*._ This pattern is consistent in all pairwise comparisons except for *H. e. chestertonii*. We further focused on these sets of unique ATAC-seq peaks to understand whether the *F*_*st*_ signal originated from strong selection or reduction of gene flow by studying the relationship between *F*_*st*_ and *D*_*xy*_ (absolute divergence). We extracted high *F*_*st*_ values (above

0.95 percentile) from the unique peaks of the significant pairwise comparison from Figure 4A, and compared them with background noise (average of 1000 ranked random sampling) from each comparison. Five of the seven pairs show a higher *D*_*xy*_ than the background (Figure 4B, significance is based on the KS tests), supporting that these regions belong to genomic regions that oppose gene flow.

### Simulation results

Our results suggest that *F*_*st*_ can act as an indicator of barrier loci; however, its signal may be obscured under certain conditions. In cases of very low divergence—such as across hybrid zones—the signal may be too weak to rise above background noise. Conversely, when divergence is too high (e.g., in all comparisons involving *H. e. chestertonii*), the background differentiation may have caught up with the selective signal, making it difficult to detect barriers. To test these observations and assess the theoretical soundness of our hypothesis, we conducted simulations under varying levels of gene flow between two populations. The simulation (Fig. S11) demonstrated that when gene flow is high (m=0.1), the *F*_*st*_ values for both the regions under selection and the neutral regions are indistinguishable, reaching a dynamic equilibrium at a very low Fst level (maximum 0.000125; top right panel). When gene flow decreases, a separation occurs, with the weakly selected regions having a higher *F*_*st*_ than the neutral regions, and these also reach a dynamic equilibrium (top left panel). When the gene flow is very low or completely absent (bottom panels), there is an initial separation where the regions under weak selection have a higher *F*_*st*_ than the neutral ones on the same chromosome. However, the two distributions tend to converge over time, and in the absence of gene flow, they converge around 100,000 generations. Our theoretical simulations support the empirical findings, as they exhibit patterns consistent with those observed in the real data.

### Evolutionary history of structural variants

Across the constructed pan-genome, 49.1% of the sequence was conserved among all H. erato populations, forming locally collinear blocks. In contrast, 37% of the pan-genome consisted of non-homologous sequences, with population-specific proportions ranging from 10% in *H. e. etylus* to 16%in *H. e. chestertonii*. An additional 13.9% of the genome was partially conserved and shared among some, but not all, populations (Fig. 5A). The conserved regions exhibited near-complete synteny across all populations, interrupted only by 16 transpositions totaling 196,166 bp (0.03% of the pan-genome), with individual events ranging from 2,016 to 66,539 bp (Fig. 5A). To explore structural variation more broadly, we analyzed genome-wide correlations between structural variant (SV) frequency and chromosome length. SV frequency was inversely correlated with chromosome length (Fig. S1), likely reflecting the effects of linked selection. Longer chromosomes typically experience lower recombination rates per base pair, intensifying selective sweeps and purifying selection, which reduces overall diversity (Edelman et al., 2019; Cicconardi et al., 2021). This pattern mirrors the distribution of SNP variation (Ruggieri et al., 2022).

**Figure 5:**
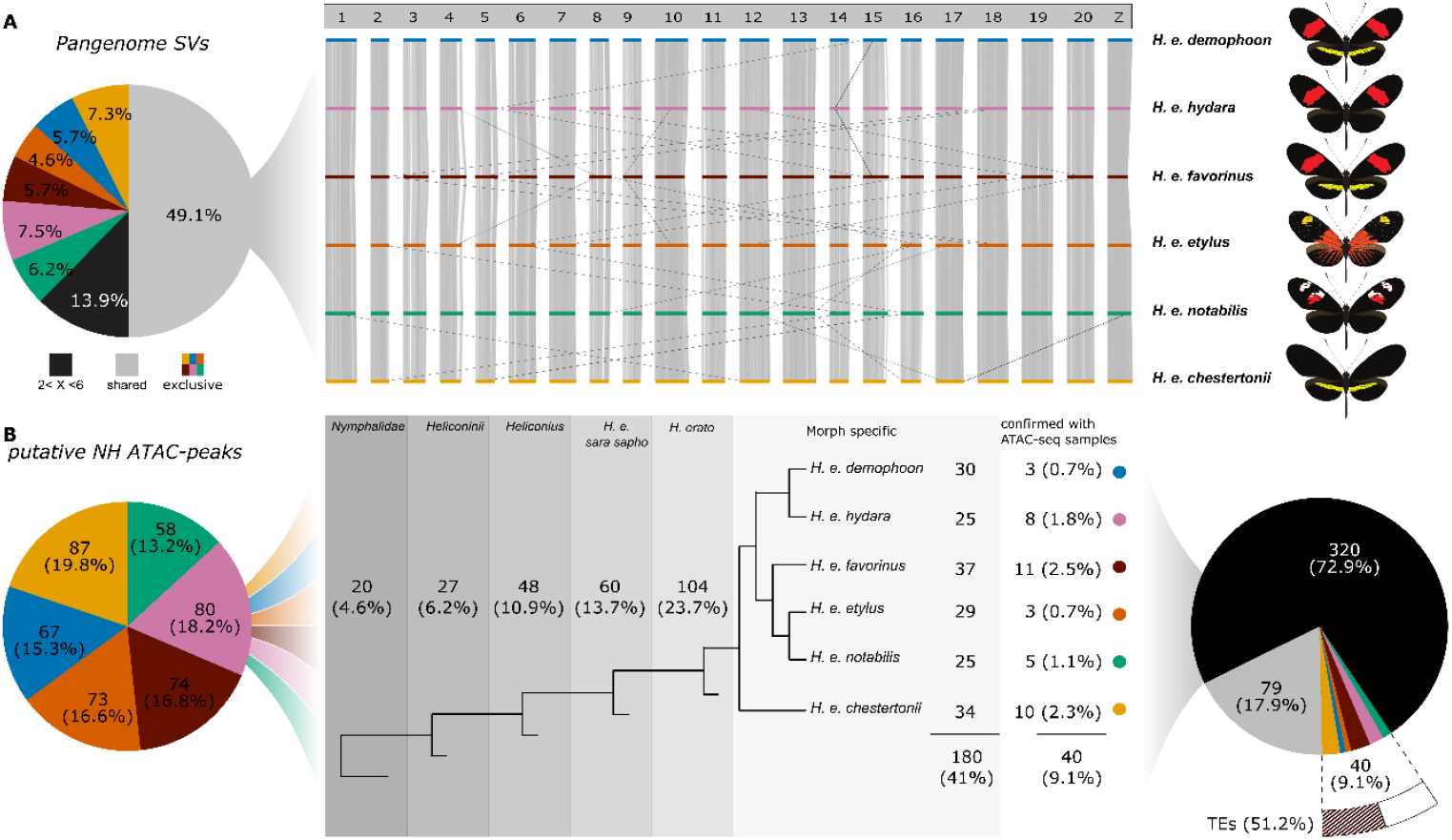
Structural and Regulatory Divergence Among Populations. **(A)** Genome-wide synteny plot of shared sequences, with each line corresponding to a different population. Gray lines represent sequences longer than 5 kbp. Only transpositions confirmed with genome reads, as described in Materials and Methods, are shown. **(B)** Pie chart depicting the number of ATAC-seq peaks that fall within non-homologous sequences (NH peaks) for each population. The second panel shows the distribution of these putative NH peaks at different phylogenetic levels within the Nymphalidae family. The rightmost pie chart summarizes how many putative NH peaks are confirmed as non-homologous based on ATAC-seq data, as detailed in the Materials and Methods section. Additionally, it reports the percentage of exclusive non-homologous peaks covered by transposable elements.

Using a single genome per population, we initially identified a conserved genomic core along with population-specific SVs. However, incorporating multiple individuals per population and applying ATAC-seq in a broader phylogenetic context revealed a more complex structural landscape. Initially, we identified several population-specific putative non-homologous peaks by intersecting ATAC-seq peaks with non-homologous regions from the pan-genome: 66 in *H. e. demophoon*, 88 in *H. e. hydara*, 70 in *H. e. favorinus*, 53 in *H. e. etylus*, 74 in *H. e. notabilis*, and 88 in *H. e. chestertonii*. To assess the evolutionary origins of these peaks, we searched for homologous sequences in other *H. erato* subspecies and across broader Heliconiine and Nymphalid datasets (Cicconardi et al., 2022; Fig. 5B). Of the 439 candidate non-homologous ATAC-seq peaks, only 41% (180) were exclusive to the six studied populations. The remaining peaks were found in other *H. erato* populations (23.7%, 104 peaks), the *H. e. sara/sapho* clade (13.7%, 60), other *Heliconius* species (10.9%, 48), other Heliconiini (6.2%, 27), and other Nymphalidae (4.6%, 20).

To further validate population-specificity, we intersected the putative non-homologous peaks of each population with the population-specific ATAC-seq data from the other populations. Since each morph’s dataset includes three individuals, it offers greater resolution for detecting polymorphism. This comparison filtered out shared elements and confirmed that only 9.1% (40 peaks) were truly non-homologous (Fig. 5B). These results show that relying on a single genome per population substantially overestimates non-homologous content by failing to capture within-population variation.

Next, we examined the 40 confirmed ATAC peaks and their associated genomic context. These regions showed high transposable element (TE) content, with an average of 51.2% of base pairs occupied by TEs, and four peaks were entirely composed of TEs. The TE composition across all populations was dominated by RC/Helitrons (22.8%), followed by SINEs (12.3%) and LTR/Gypsy elements (8.8%). LINE elements, including LINE/Jockey and LINE/L2, were also prominent, while DNA transposons such as DNA/TcMar-Mariner and DNA/PiggyBac were present in lower proportions (Fig. S12). Population-specific patterns emerged: *H. e. demophoon* and *H. e. notabilis* showed high RC/Helitron and LTR/Gypsy content; *H. e. hydara* had a more balanced TE composition, including abundant LINEs and DNA transposons; H. e. favorinus exhibited a diverse TE profile; *H. e. etylus* had a notable abundance of LINE/L2 elements; and *H. e. chestertonii* showed high levels of SINE and RC/Helitrons. These patterns suggest population-specific TE proliferation, possibly shaped by local selection or drift.

We identified the nearest gene to each of the 40 confirmed ATAC peaks to infer potential functional relevance (Table S8). These 39 genes (one peak lacked a nearby gene) include a mix of annotated and hypothetical proteins, some with well-known roles in development and physiological regulation. Notable examples include: Cortex/Ivory, a key pigmentation and wing patterning gene in Heliconius (Livraghi et al., 2021); BLOC-1, involved in melanosome development and pigment variation in both mice and insects (Di Pietro et al., 2006; Zhang et al., 2017); Juvenile hormone acid O-methyltransferase, which regulates growth, size, and reproduction timing (Dominguez et al., 2018; Guo et al., 2021). The combination of high TE content and proximity to genes involved in development, pigmentation, and hormonal regulation suggests that these regions could influence gene expression and contribute to adaptive divergence between populations.

## DISCUSSION

### Multilocus Architecture of Incipient Reproductive Isolation

The genetic basis of early reproductive isolation remains a central question in evolutionary biology: Is isolation driven by a few major-effect loci or by numerous loci with varying effect sizes? We address this by analyzing *Heliconius erato*, a butterfly species with populations at different stages of divergence across Central and South America. Earlier genomic studies, Van Belleghem et al. (2017), identified only a few genomic regions—primarily wing patterning genes that modulate mimetic warning color variation —with elevated *F*_*st*_, supporting the hypothesis that a small number of major-effect loci account for most genome-wide patterns of divergence. These color pattern genes not only directly affect a butterfly’s survival but also reinforce mate choice (Southcott & Kronforst, 2018; Kuo et al., 2024; Rossi et al., 2020, 2024), making them potential targets for early reproductive isolation. However, when *H. erato* populations were compared with the more geographically isolated *H. himera* (McMillan et al. 1997; Merrill et al. 2014), and *H. e. chestertonii* (Muñoz et al. 2010; Van Belleghem et al. 2018), genome-wide divergence increased to the point that *F*_*st*_ peaks near known color pattern loci became undetectable (Van Belleghem 2021). These results suggest that genome-wide patterns of divergence appear abruptly rather than gradually and, importantly, may remain undetectable in the presence of gene flow.

To revisit this hypothesis, we constructed an *H. erato* pan-genome using six reference genomes from populations spanning a range of geographic distances and levels of gene flow (Fig. 1A–B). We reanalyzed resequencing data from Van Belleghem et al. (2017), focusing on hybrid zones in Panama (*H. e. hydara × H. e. demophoon*) and Ecuador (*H. e. notabilis × H. e. etylus*) (Fig. 2). Contrary to approaches of earlier studies, which identified only a few divergence peaks, our analysis revealed an elevated genome-wide *F*_*st*_ divergence and selective sweeps (Fig. 2). These additional divergent regions point to new potential targets of selection beyond wing patterning, which may be functionally tied to behavioral, ecological, or physiological adaptations (Tables 1, S2, S5). Many of these regions also show elevated *D*_*xy*_, consistent with the presence of barrier loci (Cruickshank & Hahn, 2014; Feder et al., 2012).

This improved resolution to characterize patterns of genomic divergence in the early stages of population differentiation stems from enhanced mapping using population-specific pan-genome assemblies and a finer-scale analysis based on 1 kbp *F*_*st*_ windows (Fig. S13). Previous studies, such as Van Belleghem et al 2017, relied on a single *H. e. demophoon* reference genome and used 50 Kbp sliding windows, which likely masked smaller signals of selection. Recent work has highlighted that selection, gene flow, and recombination heterogeneity demand higher-resolution approaches (Diopere et al., 2018; Yuan et al., 2021; Lewis et al., 2019). Our use of 1 Kbp windows proved effective across allopatric populations, except for comparisons involving *H. e. chestertonii* from Colombia (Fig. 3). This results likely depend from *H. chestertonii* being the only population of *H. erato* to show not only wing color based interspecfic mate choice, but also evidence of hybrid unviability, which can explain its higher overall genomic divergence, which can obscure selection signals (Van Belleghem et al., 2021). For example, *F*_*st*_ distributions between *H. e. demophoon* and *H. e. chestertonii* display broader variance around the mean, reducing signal sharpness compared to other comparisons, where density peaks near the mean and only a few values deviate widely (Fig. S10).

### Epigenetic Divergence and Regulatory Variation Under Gene Flow

A multilocus structure is also evident in epigenetic divergence (Fig. 3). ATAC-seq of wing tissue reveals widespread population-specific chromatin accessibility, implying regulatory divergence beyond the known patterning genes (Fig. 2, 3A). These differences may influence diverse wing traits, including structure, shape, pheromone production, and sensory adaptations (Table S5). Enrichment analyses show divergence in transcription factor binding motifs, suggesting possible altered regulatory networks (Fig. 2B). Across hybrid zones, differences in chromatin accessibility show only a weak correlation with elevated *F*_*st*_ (Fig. 4A, 4B). This may reflect a mismatch in resolution: *F*_*st*_ is calculated over 1,000 bp windows, while ATAC-seq peaks typically span ∼400 bp. Regulatory divergence may result from just a few causal mutations, which may not be sufficient to generate strong *F*_*st*_ signals at this scale. In addition, divergence in chromatin accessibility may be driven by upstream regulators of chromatin remodeling (Adrian-Kalchhauser et al., 2020), further decoupling it from local sequence divergence. However, in population pairs with reduced gene flow, unique ATAC-seq peaks overlap with regions of high *F*_*st*_ and *D*_*xy*_, suggesting resistance to introgression (Cruickshank & Hahn, 2014; Charlesworth, 2006) that grows with overall genetic divergence. These results suggest that chromatin divergence— i.e., differences in gene regulation—can act as barrier loci limiting gene flow, highlighting a more prominent role for epigenetic variation in the speciation process. Taken together, these results suggest that there may be a weak increase in *F*_*st*_ in the population specific ATAC-seq data between adjacent populations, but it is possibly obscured by gene flow or because of lack of resolution due to masking and filtering of indel variation. When gene flow is reduced or absent, the signal becomes more evident and increases with geographic distance.

These results align with recent work suggesting a role for chromatin accessibility in speciation. For instance, Smith et al. (2015) found that epigenetic divergence exceeded genetic divergence in *Etheostoma olmstedi*. Similarly, Lewis et al. (2020) showed that selection on *Optix* target sequences helps maintain phenotypic boundaries in *Heliconius* hybrid zones.

Genomic divergence in *Heliconius* has already been linked to a highly polygenic architecture, particularly in low-recombination regions that resist introgression (Martin et al., 2019). Likewise, hybrid incompatibilities in butterflies follow a polygenic model, particularly on the Z chromosome, where small-effect loci accumulate (Xiong et al., 2023). This supports a polygenic threshold model, where reproductive isolation arises incrementally. Our findings suggest this multilocus architecture emerges earlier in divergence than previously thought.

### Targets of Selection for Incipient Reproductive Isolation in *Heliconius erato*

Genomic research on *Heliconius* butterflies has long focused on wing color pattern genes, yet complete reproductive isolation in this genus has been shown to extend beyond coloration (Meier et al. 2021). A broader suite of traits, including neural pathways, chemical communication, and sensory perception, plays a crucial role in ecological adaptation and speciation (Mérot et al., 2017, Montgomery & Merrill, 2017; Rivas-Sánchez et al., 2024, Wright et al., 2024). Our genomic investigation reveals a complex, genome-wide signature of divergence that likely reflects incipient reproductive isolation across multiple functional axes. Unlike earlier studies that relied on a single reference genome (*H. e. demophoon*) and broad-scale divergence metrics (e.g., 50 kb FST windows) (Van Belleghem et al, 2017), our approach uses denovo reference genomes for each population, finer-resolution methods, and chromatin accessibility, uncovering a markedly heterogeneous and polygenic landscape. This emerging polygenic architecture in the presence of gene flow highlights divergence far beyond the canonical wing color pattern genes (*WntA, optix, cortex locus*), identifying strong candidate genes in a suite of relevant traits across pathways affecting behavior, development, physiology, and sensory adaptation.

Divergence in circadian regulation and behavioral timing likely plays a key role in structuring reproductive schedules, with genes linked to circadian clocks and sex-specific expression influencing the timing of mating activity and courtship displays (Rivas-Sánchez et al., 2023; Dell’Aglio et al., 2022). These shifts can reduce hybridization by reinforcing temporal or behavioral isolation mechanisms. Alongside behavioral rhythms, differences in cuticle formation and pigment biosynthesis highlight another axis of divergence. Genes involved in melanin production, chitin remodeling, and cuticle architecture suggest modifications in pigmentation, waterproofing, and even pheromone dispersion (Montejo-Kovacevich et al. 2021). Such traits influence both environmental adaptation through desiccation resistance or camouflage and mate recognition, where subtle changes in wing coloration or texture may serve as visual mating cues and ecological performance.

Morphological divergence emerges as another important trait, driven by changes in genes that regulate developmental patterning and tissue morphogenesis. Variation in components of Wnt, Notch, and EGFR signaling pathways, along with chromatin modifiers and cytoskeletal regulators, points to differences in body plan specification and organ formation. These developmental changes may underlie subtle yet crucial differences in wing shape, sensory organ development, traits that can influence mate choice, and habitat adaptation. This morphological variation is paralleled by divergence in neural signaling and sensory perception (Wright et al., 2024; Montgomery & Merrill, 2017; Rivas-Sánchez et al., 2024). Our results show differentiation in genes tied to synaptic transmission, chemosensory reception, and photoreception, thus suggesting population-specific tuning of sensory systems. Such modifications can alter how individuals perceive visual and chemical stimuli in their environment (Montgomery et al., 2021), influencing both mate recognition and host plant selection. These sensory shifts are likely to contribute strongly to pre-mating isolation and ecological specialization.

At the physiological level, we also observe divergence in hormone metabolism and detoxification pathways. Genes involved in juvenile hormone regulation, steroid biosynthesis, and xenobiotic detoxification reflect changes in developmental timing, pheromone production, and adaptation to local environmental toxins (e.g., host plant chemistry). These physiological shifts may synchronize development with local ecological conditions, reinforce reproductive timing, or influence female receptivity, all critical components in maintaining population boundaries (Yao et al., 2025; Cao et al., 2013).

Finally, divergence in reproductive and post-mating processes further strengthens isolation. Genes associated with seminal fluid composition, female post-mating responses, and reproductive immunity suggest that barriers to gene flow can persist beyond courtship and copulation. These molecular incompatibilities may reduce fertilization success, bias sperm competition, or modulate female re-mating behavior, contributing to reproductive isolation even when mating occurs.

### Structural Variation in Speciation

Structural variation is a major driver of genome evolution and speciation (Thawornwattana et al., 2023), yet single-genome-per-population approaches fail to capture much of this diversity. Our chromatin accessibility analysis showed that many population-specific ATAC-seq peaks are polymorphic, highlighting extensive structural variation within populations. Of the non-homologous regulatory elements identified from the pan-genome, only ∼9% were confirmed as truly non-homologous after incorporating multiple individuals. Some of these peaks are also found in other Nymphalidae species, suggesting the retention of ancestral variation. Notably, some of the confirmed non-homologous peaks reflect distinct, population-specific transposable element content (Fig. 4; Fig. S12), supporting the role of mobile elements in regulatory evolution and divergence (Oliver & Greene, 2009; Ruggieri et al., 2022; Ray et al., 2019).

## CONCLUSION

Our findings challenge the long-standing assumption that genomic divergence in populations experiencing high gene flow, such as *Heliconius erato* hybrid zones, is primarily driven by a few major-effect loci. Instead, we provide compelling evidence for a multilocus architecture, where genome-wide divergence, epigenetic modifications, and structural variation collectively influence evolutionary trajectories. The observed genome-wide divergence and extensive population-specific chromatin accessibility provide compelling evidence for incipient reproductive isolation driven by a diverse and polygenic set of traits, many of which were previously undetected under traditional approaches (Wright et al., 2024; Montgomery & Merrill, 2017; Rivas-Sánchez et al., 2024). By integrating fine-scale divergence data across neural, morphological, metabolic, sensory, and ecological dimensions, our study redefines the genomic basis of divergence in *Heliconius erato*. The presence of retained ancestral structural variants further illustrates the dynamic and heterogeneous nature of the *H. erato* genome, suggesting that both historical and contemporary forces shape current patterns of divergence. Collectively, these findings reinforce the role of polygenic adaptive divergence, ultimately driving the earliest stages of reproductive isolation (Xiong et al. 2023). This represents the foundational step in the formation of the extraordinary species diversity seen within the *Heliconius* radiation. Our study underscores the complexity of speciation, showing how genetic variation, epigenetic modification, and structural genome architecture act in concert over evolutionary time (Cicconardi et al., 2023) to initiate and maintain reproductive barriers, setting the stage for ecological diversification and lineage divergence.

## Acknowledgments

We thank S. Planas and Y. Ortiz for Illumina library preparation. We thank B. S. Collazo for assistance in all purchasing and bureaucratic paperwork. For the support of sequencing and computational resources, we thank the University of Puerto Rico Sequencing and Genomics Facility (SGF) INBRE Grant P20 GM103475 from the National Institute for General Medical Sciences (NIGMS), a component of the National Institutes of Health (NIH).

## Author contributions

R.P. conceptualized and designed the study; A.R., and R.P., wrote the original draft manuscript and assembled the figures; A.R., S.V.B, F.C., N.B., designed the methodology; A.R., S.V.B, F.C., N.B., and RP conducted the analyses; J.M., S.M., F.C., wrote, edit and comment the final manuscript; B.A.C., W.O.M., and R.P., collected samples for sequencing.

## Data and materials availability

Custom codes used are available on Gihhub:https://github.com/DNAcastigator/incipient-speciation-pangenome

ATAC-seq data are available on NCBI: [PRJNA795145](https://www.ncbi.nlm.nih.gov/bioproject/795145) 10Xdata are available at :PRJNA1240918

The necessary butterfly collection permits were obtained from the governments of each country and importation permits were obtained from the Ministry of the Environment of Panama, according to the Panamanian Government and STRI regulations. Rearing and experimentation complied with local government regulations on containment and handling.

Samples collected in Peru were obtained under permits 0289-2014-MINAGRI-DGFFS/DGEFFS, 020- 014/GRSM/ PEHCBM/DMA/ACR-CE, 040–2015/GRSM/PEHCBM/DMA/ACR-CE, granted to Dr

Neil Rosser, and samples from Panama were collected under permits SEX/A-3-12, SE/A-7-13 and SE/AP-14-18.

## Supplementary Materials

Materials and Methods

Figs. S1 to S14

Tables S1 to S8

