## Supplementary and M&M for "Polygenic basis of incipient reproductive isolation in hybridizing populations is revealed by pangenomic and epigenetic divergence"

### MATERIAL AND METHODS

#### *Genome assemblies*

New genomes were generated for five *H. erato* populations (*H. e. notabilis, H. e. etylus, H. e. hydara, H. e. chestertonii, and H. e. favorinus)*. Although some sources classify *Heliconius erato chestertonii* as a distinct species (Jiggins, 2017; Rivas-Sánchez et al., 2024), the majority, particularly those focused on speciation, support its classification as an instance of incipient speciation (Arias et al 2008). Consequently, in this study, *H. e. chestertonii* is considered a geographic subspecies of *H. erato*. For each genome, we extracted high-molecular-weight DNA from flash-frozen pupas. Library preparation using 10X Chromium technology for linked reads (10x Genomics, San Francisco, USA) and Illumina sequencing was carried out by Novogene Co., Ltd, with a target coverage of 100x. We assembled the linked-read sequencing data using the Supernova 2.1.1 assembler (Weisenfeld et al. 2014) using the default recommended settings and a maximum number of reads of 200 million. Raw assembly outputs were transformed to fasta format using the pseudohap2 option to generate two parallel pseudo-haplotypes from the diploid genome. Quality control of the genomes was performed using genome-wide statistics calculated on the phase blocks. Synteny with the *H. e. demophoon* V1 genome was performed using Tigmint v1.2.3 (Jackman et al. 2018). Finally, BUSCO was used to assess genome assembly and annotation completeness (Simão et al. 2015).

#### *Pan-genome*

Using seq-seq-pan (Jandrasits et al., 2018), we generated an *H. erato* population pan-genome aligning a total of six *H. erato* populations: *H. e. demophoon, H. e. hydara, H. e. notabilis, H. e. etylus, H. e. favorinus, H. e. chestertonii*. The pan-genome is a collection of core and variable genomic regions derived from multi-genome alignments, and it can be used as a reference for mapping NGS data. The *H. e. demophoon* v1 reference genome was used as the first genome in the genome list to ensure the resulting pan-genome alignment was ordered correctly. The *H. e. demophoon* chromosome 21 final coordinates on the pan-genome is 688,640,245; sequences after that position were not used to avoid spurious feature mappings. To obtain lineage-specific sequences, we first recorded the sequence coordinates of each genome relative to the pan-genome as in Ruggieri et al. 2022. We then subtracted a merged library of sequence coordinates of all other genomes from these coordinates using BEDTools v2.27.1, obtaining only the sequences exclusive to each genome (Quinlan and Hall, 2010).

#### *ATAC-seq library preparation*

ATAC-seq libraries were prepared following the protocol described by Lewis and Reed (2019), with some modifications. Caterpillars of each species were raised on their respective host plants until the wandering stage at the 5th instar. Live larvae were briefly placed on ice, then pinned and dissected in 1X ice-cold PBS. Developing wings were dissected from 5th instar caterpillars, and the left and right forewings and left and right hindwings were pooled, respectively.

Next, the tissues were homogenized, and the nuclei were extracted by submerging them in 350 µl of sucrose solution inside 2 ml dounce homogenizers. After homogenization on ice, the solution was centrifuged to collect the nuclei. The pellet was resuspended in a cold lysis buffer to burst the cell membranes and release the nuclei. The nuclei were examined under a microscope with a counting chamber to confirm dissociation, assess their concentration, and calculate the number of nuclei (approximately 400,000) to be used for further processing. This number was determined based on the requirement to obtain the same amount of DNA as present in 50,000 human nuclei, which is the optimized amount for ATAC-seq (Buenrostro et al., 2013).

For quality control, a 15 µl aliquot of the nuclear suspension was stained with trypan blue and imaged at 64x magnification using a hemocytometer. After confirming nuclear quality and concentration, a subsample of the nuclear volume corresponding to 400,000 nuclei was pelleted, resuspended in a transposition mix containing Tn5 enzyme (Illumina DNA Prep) in a transposition buffer, and incubated at 37 °C for exactly 30 minutes. The resulting tagged fragments were purified using a PCR MinElute Purification Kit and amplified with custom-made Nextera primers and a NEBNext High-fidelity 2x PCR Master Mix (New England Labs). The amplified libraries were sequenced as 37 to 76 bp paired-end fragments using NextSeq 500 Illumina technology at the Sequencing and Genomics Facility of the University of Puerto Rico.

***ATAC-seq analysis***

ATAC-seq reads were trimmed with TRIMMOMATIC (Bolger et al 2014) using 2:30:10:2:keepBothReads LEADING:3 TRAILING:3 MINLEN:32. All the reads were mapped to the pan-genome with bowtie2 (Langmead & Salzberg 2012). Reads with a phred score lower than 20 and non-uniquely mapped were discarded for further analysis (SAMTools,-q 20, -f 0 × 02). Duplicates were removed with Picard MarkDuplicates (http://broadinstitute.github.io/picard/). ATAC-seq peaks were called using MACS2 (Zhang et al 2008) with the genome size set to the pan-genome size (-g 688640245) and the q-value stricter than the default (-q 0.001). We measured ATAC-seq quality using the Fraction of Reads in Peaks (Frip) score, which is the ratio of reads mapped inside the identified ATAC-seq peak to the total number of mapped reads. We retained only those samples with a score above 0.2, which is generally acceptable (https://www.encodeproject.org/atac-seq/; Table S3).

The ATAC-seq peaks for each sample in each group were intersected with bedtools and retained only if all three samples had the same peak with a reciprocal minimal 50% overlap (3/3 peak). The resulting peak lists were used for all analyses in the project's main figures. We also identified a list of variable peaks present in at least two of the three samples (2/3 peaks) that were used in supplementary analyses to support the main finding. Once we obtained the list of conserved peaks, the ATAC-seq reads mapping within the peaks' intervals were counted with" bedtools coverage -count.", generating the reads count matrix used for the plot in Figure 2A and Figure 3A.

Peaks were considered "shared" between populations when they aligned at least 50% (using bedmap, from the bebops toolkit, Neph, et al 2012; using the option --fraction-either 0.5). In the case that a larger peak in one population aligned with multiple smaller peaks in the other population, the smaller peaks were merged, and the reads were. Unique peaks were selected by subtracting every peak identified in one population from the list of 3/3 peaks of the other population (bedtools subtract -A). In this way, we identified peaks that were consistently present in the population of reference and completely absent in the other population. These peaks will be hereafter referred to as "Unique ATAC-peaks". For the putative non-homologous ATAC-seq peaks, the lineage-specific sequences obtained from the pan-genome were intersected with the list of 3/3 peaks for each population. Subgroups of these ATAC-seq peaks were annotated to the closest gene using Homer's "annotatePeaks.pl" (Heinz et al 2010). The closest gene to an ATAC-seq peak was used as a proxy for the potential target of the ATAC-seq peak.

***Resequencing***

Resequencing data was obtained from Van Belleghem et al. 2017. The dataset includes individuals with divergent *H. erato* mimetic subspecies populations that are found closely adjacent and sometimes connected by hybrid zones. To analyze the genomic data, the existing 100 bp paired-end Illumina resequencing data were mapped to the consensus pan-genome reference genome using BWA v0.7.13 (Li & Durbin 2009) with default parameters. We removed PCR duplicates using Picard v1.1 (http://picard.sourceforge.net) and sorted the data using SAMtools (Danecek 2021). Genotypes were called using the Genome Analysis Toolkit (GATK) Haplotypecaller (Van der Auwera et al 2013) with default parameters. For joint genotyping, we used GATK's genotypeGVCFs with default parameters. We considered a minimum expected heterozygosity of 0.025 to match the populations' high heterozygosity and grouped individuals based on subpopulation and sampling location. In the downstream analysis, genotype calls were evaluated based on the following criteria: Quality (QUAL) $\geq$ 30, minimum depth $\geq$ 10, maximum depth $\leq$ 100 (to avoid false SNPs in repetitive regions), overall depth $\leq$ 100 times the number of samples, strand bias (FS) < 200, Quality by depth $\geq$ 5, and genotype quality (GQ) $\geq$ 30 for variant calls.

*Population genomics analysis*

Genome-wide selective sweep signatures were identified using SweepFinder2 (Degiorgio et al., 2016). A custom Python script was used to generate allele counts for biallelic SNPs, which were polarized with *H. hermathena*. The default settings were used, with 1000 bp window size. $F_{st}$ and $D_{xy}$ were calculated using Simon Martin's (popgenWindows.py; https://github.com/simonhmartin/genomics_general) using the non-overlapping sliding windows of 1000, 5000, 10000, and 50000 bp. To capture the potentially small effects of regulatory elements identified by ATAC-seq (which average ~400 bp in length), we calculated $F_{st}$ using 1,000 bp windows. Regulatory regions can evolve rapidly and play important adaptive roles, but because they are typically small, using larger windows could miss these signals leading to an underestimation of how strongly selection has shaped genomic differences. This approach increases background noise since the small window size tends to have higher noise due to technical issues (da Silvia Ribeiro et al 2022) and biological variability (Weir et al 2005), which may not reflect meaningful population-wise biological differences (Figure S13;S14). Therefore, we could not use the $F_{st}$ values as they were generated; a strategy was needed to identify a possible biological signal above the background noise. Initially, the $F_{st}$ data was grouped every 10 points (each point is the value of $F_{st}$ in one 1000 bp window), wherein we calculated both the number of points exceeding the average $F_{st}$ value (above average distribution) and the maximum $F_{st}$ value within each group. For a singal to be considered valid (strongly divergent form the background and not due to chances), we established specific criteria: a) the signal must demonstrate a higher point density above the average (above average distribution value for that group exceeding the 0.99 quantiles of the overall genome above average distribution), and b) it must include at least one data point surpassing the 0.99 percentile of the entire $F_{st}$ distribution.

***Integration of ATAC-seq and $F_{st}$ patterns***

To explore the potential relationships between $F_{st,}$ selective sweep, and chromatin dynamics, we primarily used two approaches that aimed at identifying signals above background noise: the Kolmogorov-Smirnov test (two-sample KS-test) and a custom test based on the binomial test (Custom binomial test). All the analyses were performed in R using the $F_{st}$ calculated in 1 kbp windows size unless otherwise noted.

*ks-test:* The ks-test is a non-parametric statistical test used to understand whether two distributions come from the same underlying one. In this study, the first distribution was composed of the combined unique ATAC-seq peaks from the 3/3 list of the two populations under study; the second distribution was an average of 1000 random sampling of the same number of ranked points from the genome. The frequencies for the random sampling were modeled on the distribution of all the ATAC-seq peaks identified in the two populations. The test was run with the null hypothesis as “less” because a lower CDF (cumulative distribution function) represents higher values from the first distribution. Hence, this test verified that the unique ATAC-seq peak distribution had overall higher values of $F_{st}$ than the background in the populations studied. The test outputs p-values as well as a D-statistic, which can be interpreted as the maximum absolute difference between the CDF distribution that we used as a proxy for the intensity of $F_{st}$ signal in unique ATAC-seq regions compared to the genome background for further analysis.

*Custom binomial test:* The Custom binomial test splits the genome into 10 $F_{st}$ intervals (increasing by 0.1), then calculates an expected probability to find a unique ATAC-seq peak in each interval (using the average of 1000 random sampling). Next, a binomial test is performed for each interval using the empirical number of unique ATAC-seq peaks as the “number of successes”, the total number of ATAC-seq peaks in that $F_{st}$ interval as “number of trials” and the average of random sampling as the “hypothesized probability of success”. The test outputs p-values for each test (null hypothesis “greater”) that are then adjusted using the Holm-Bonferroni Method correction, a more powerful and less conservative version of the classic Bonferroni correction. In summary, this approach measures whether there is a higher probability of finding a unique ATAC-seq peak in each $F_{st}$ interval than expected, highlighting a higher frequency of peaks in specific intervals compared to randomness.

*Transcription binding site motif enrichment:* Differential motif enrichment was performed on unique ATAC-seq peaks from the different adjacent pairs of populations using the XSTREME tool from the MEME suite (Grant & Bailey 20211;Bailey 2021). As a background model, we used the totality of ATAC-peaks identified for each population for their respective analyses; only motifs with an E-value < 0.05 were considered significant.

***Evolution under progressively reduced gene-flow simulations***

To support our empirical finding with a theoretical model, we conducted simulations of the evolutionary dynamics of two populations using SLiM 3 (Haller & Messer, 2019). The simulations shared the same structure and parameters, differing only in the migration rate after 2,000 generations. We modeled two populations, each with two chromosomes. The first chromosome contained 2% of nucleotides under weak selection (s = 0.01), while the remaining sites were neutral (s = 0). The second chromosome was entirely neutral and served as a control, as neutral sites on the first chromosome could experience hitchhiking effects, whereas those on the second chromosome should not.

To improve computational efficiency, we scaled down the simulations by a factor of 100, following the guidelines in the SLiM manual (page 145). Specifically, we reduced both the population size and the number of generations by 100×, while increasing the mutation and recombination rates by the same factor. The effective population size used was 10,000, assuming the original was 1,000,000.

The mutation rate was scaled from $2.9 \times 10^{-9}$ (Keightley et al., 2015) to $2.9 \times 10^{-7}$, and the recombination rate from $2 \times 10^{-8}$ (Martin et al., 2019) to $2 \times 10^{-6}$.

All simulations began with a migration rate of 0.5, which was then reduced after 2,000 generations to one of the following: 0.1, 0.00001, 0.000000001, or 0. For each parameter set, we ran 100 replicate simulations.

***Phylogenetic comparisons***

We analyzed the dynamics of non-homologous ATAC-seq peaks outside of the *Heliconius erato* species. This analysis was based on the HAL file generated by Cicconardi et al. (2023), which was updated to include *H. e. hydara*, *H. e. favorinus*, *H. e. etylus*, *H. e. notabilis*, and *H. e. chestertonii*. A genomic region was considered present at a given tree tip if it contained at least 80% of the reference sequence.

***Genome architecture variability analysis***

We studied the synteny among sequences present in all 6 populations and identified possible transpositions. The positions of the transpositions between conserved regions in different genomes were calculated with a custom script from the XMFA file of the original pan-genome. A transposition was considered real only if a) it happened between chromosomes or moved at least 100 kbp from the reference; b) the sequence moved was longer than 5k; and c) we identified at least 20 reads from the genome sequencing mapping across the beginning or the end of the transposition (that mapped partially on the transposed sequence and part of the surrounding region), indicating that the transposition was not due to a technical misassembly of the scaffold. For this last step, we used minimap2 (Li 2018) to map the sequences and samtools to map from a mapping quality of at least 10.

***TE identification***

To identify transposable elements (TEs), we used RepeatMasker (Tarailo-Graovac and Chen, 2009) on the consensus pan-genome, utilizing the TE library generated by Ruggieri et al. (2022) with default parameters.

SUPPLEMENTARY FIGURES

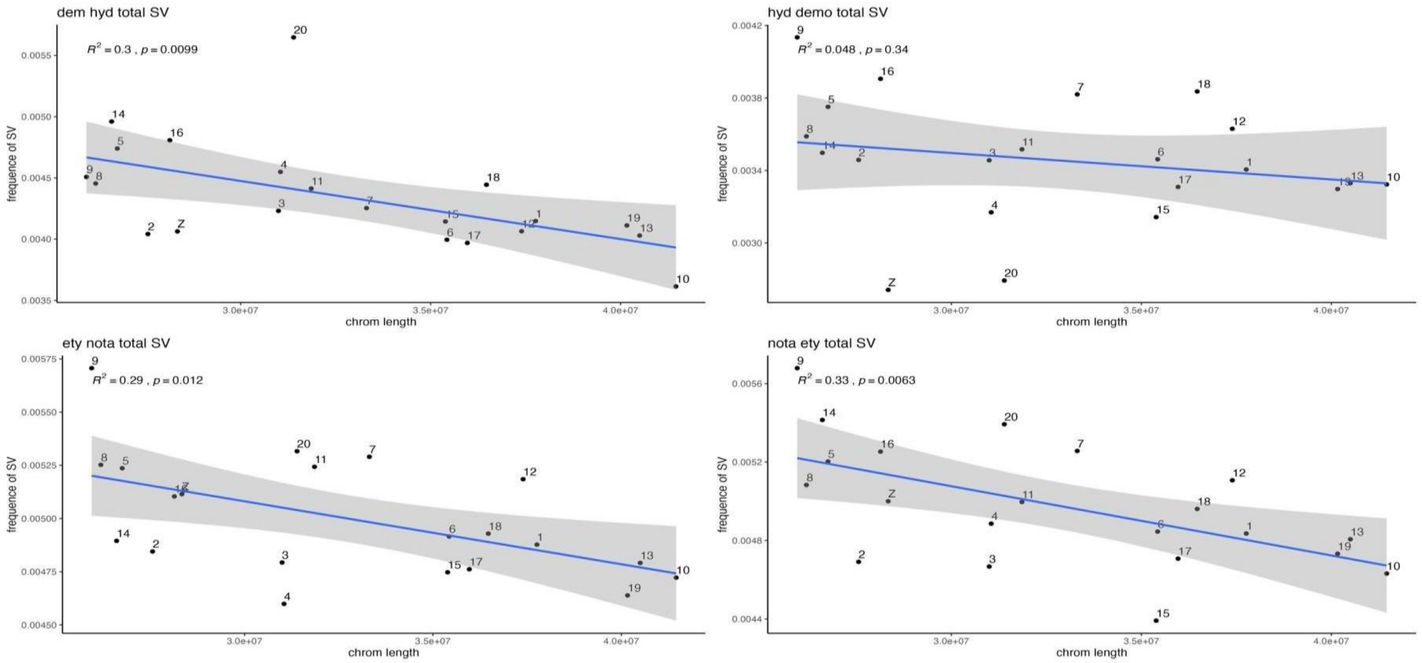

**Figure S1:** Correlation between the frequency of structural variants (SV) and chromosome length. Each panel includes the reported R-squared values and the significance of linear regression analyses. The top-left panel represents the comparison between *H. e. demophoon* and *H. e. hydara*, while the top-right panel shows the reverse comparison (*H. e. hydara* vs *H. e. demophoon*). The bottom-left panel illustrates the comparison between *H. e. etylus* and *H. e. notabilis*, with the bottom-right panel showing the reverse comparison (*H. e. notabilis* vs *H. e. etylus*).

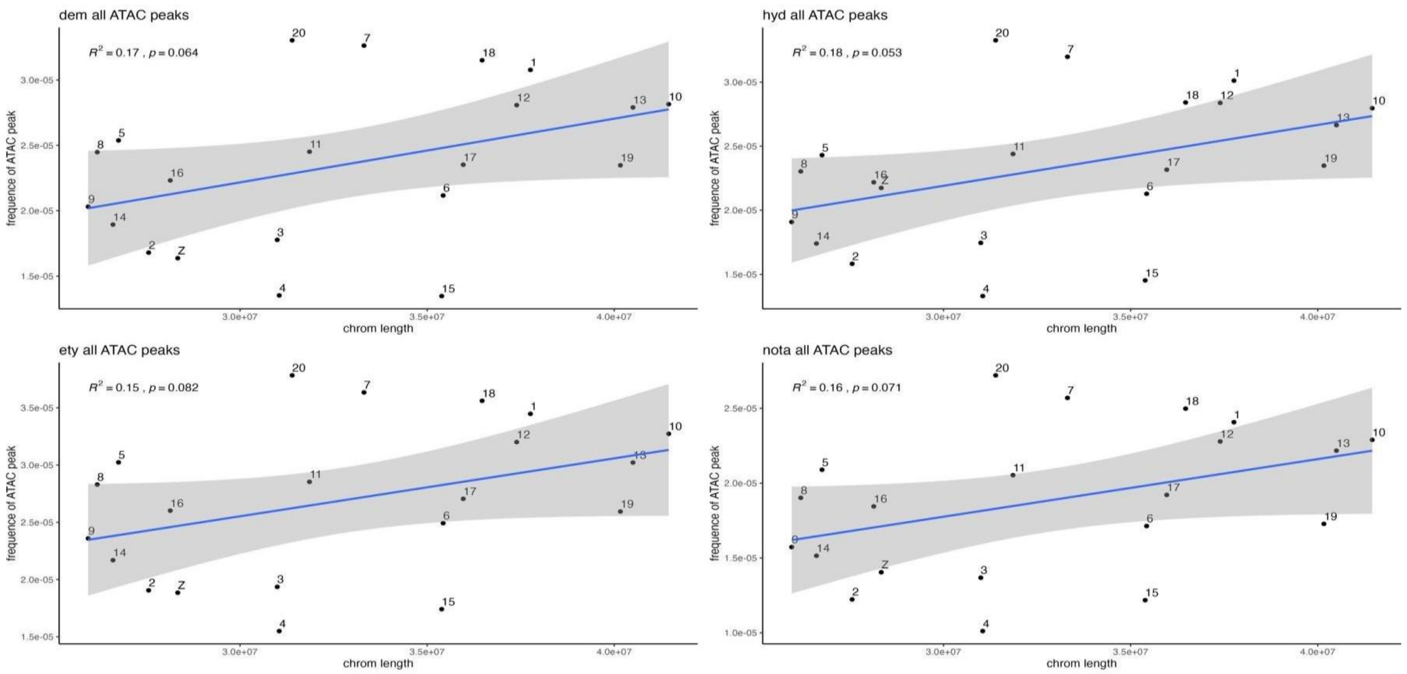

**Figure S2:** Correlation between the frequency of total ATAC-seq peaks identified in each morph and chromosome length. Each panel includes the reported R-squared values and the significance of linear regression analyses. The top-left panel represents *H. e. demophoon*, the top-right panel shows *H. e. hydara*, the bottom-left panel illustrates *H. e. etylus*, and the bottom-right panel corresponds to *H. e. notabilis*.

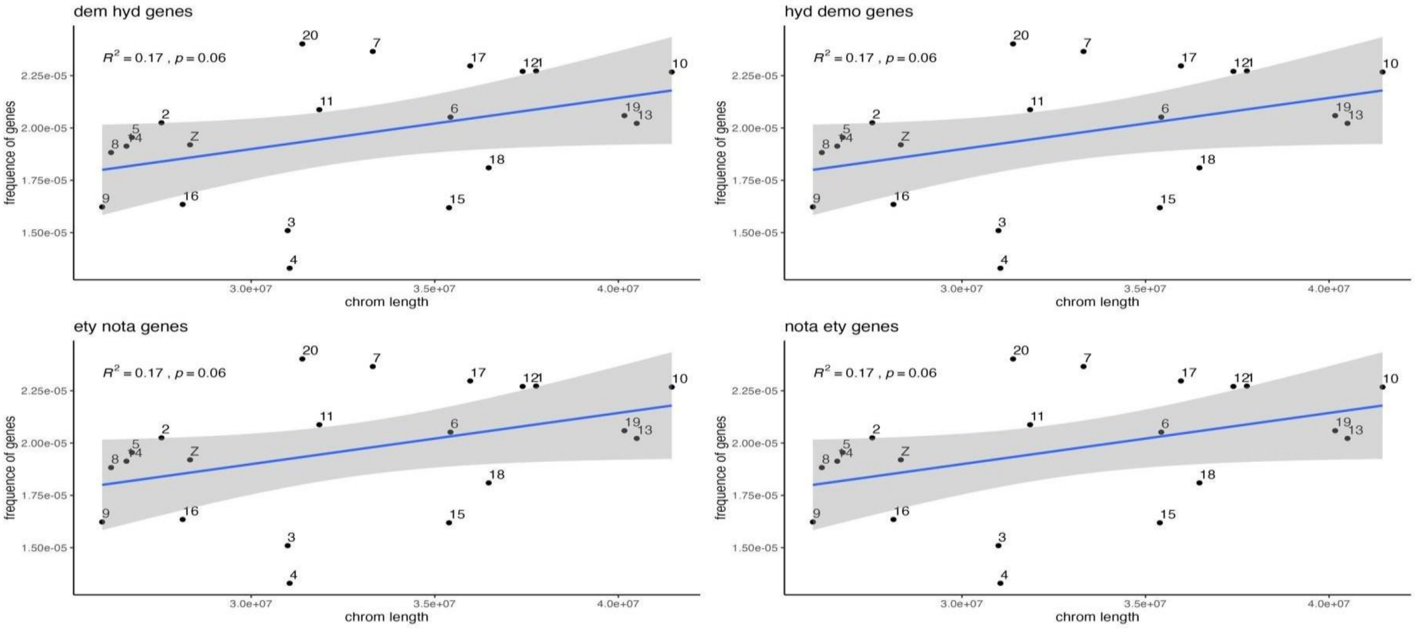

**Figure S3:** Correlation between the frequency of genes and chromosome length. Each panel includes the reported R-squared values and the significance of linear regression analyses. The top-left panel represents *H. e. demophoon*, the top-right panel shows *H. e. hydara*, the bottom-left panel illustrates *H. e. etylus*, and the bottom-right panel corresponds to *H. e. notabilis*.

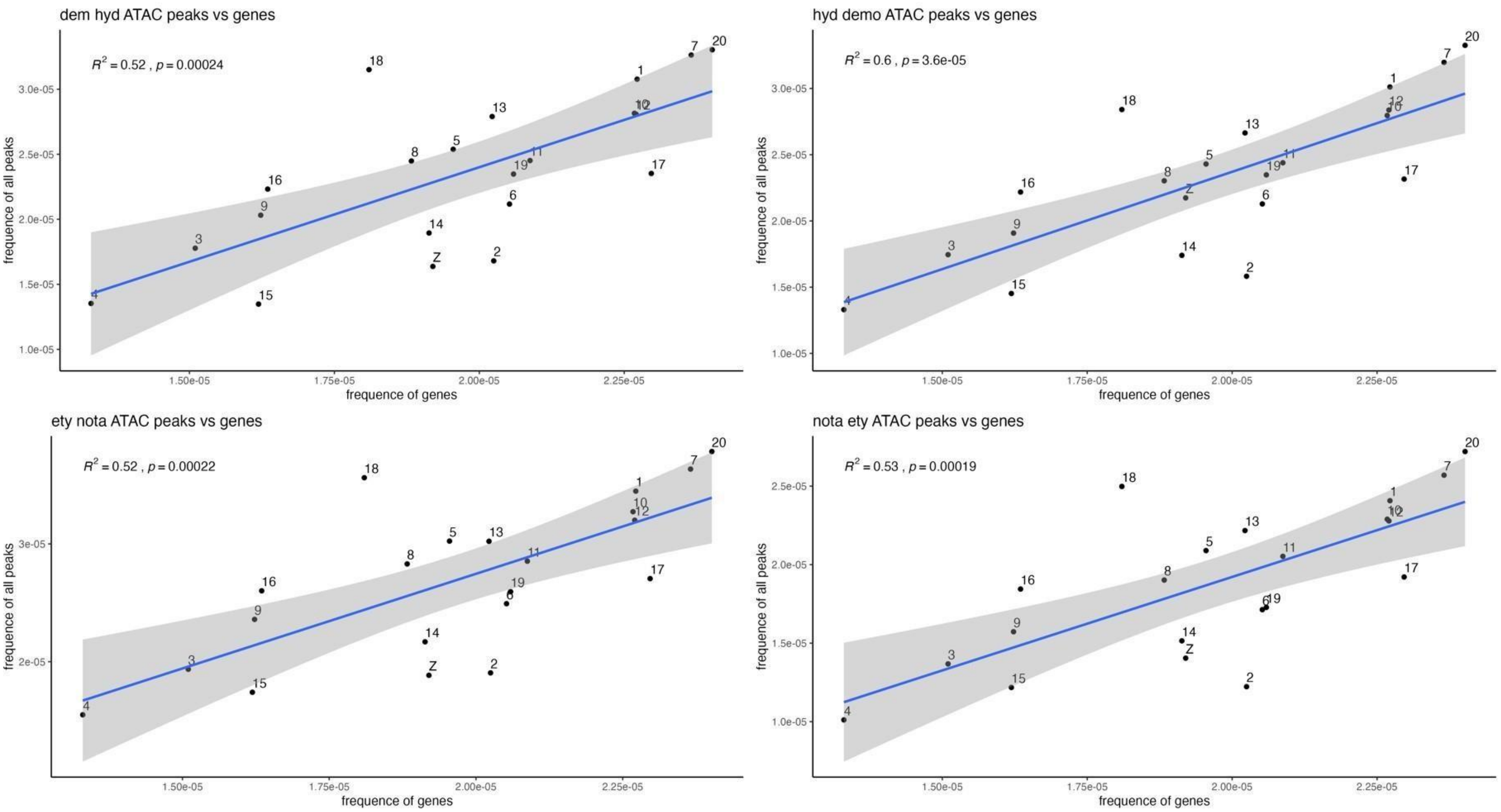

**Figure S4:** Correlation between the frequency of total ATAC-seq peaks identified in each morph and the frequency of genes. Each panel includes the reported R-squared values and the significance of linear regression analyses. The top-left panel represents *H. e. demophoon*, the top-right panel shows *H. e. hydara*, the bottom-left panel illustrates *H. e. etylus*, and the bottom-right panel corresponds to *H. e. notabilis*.

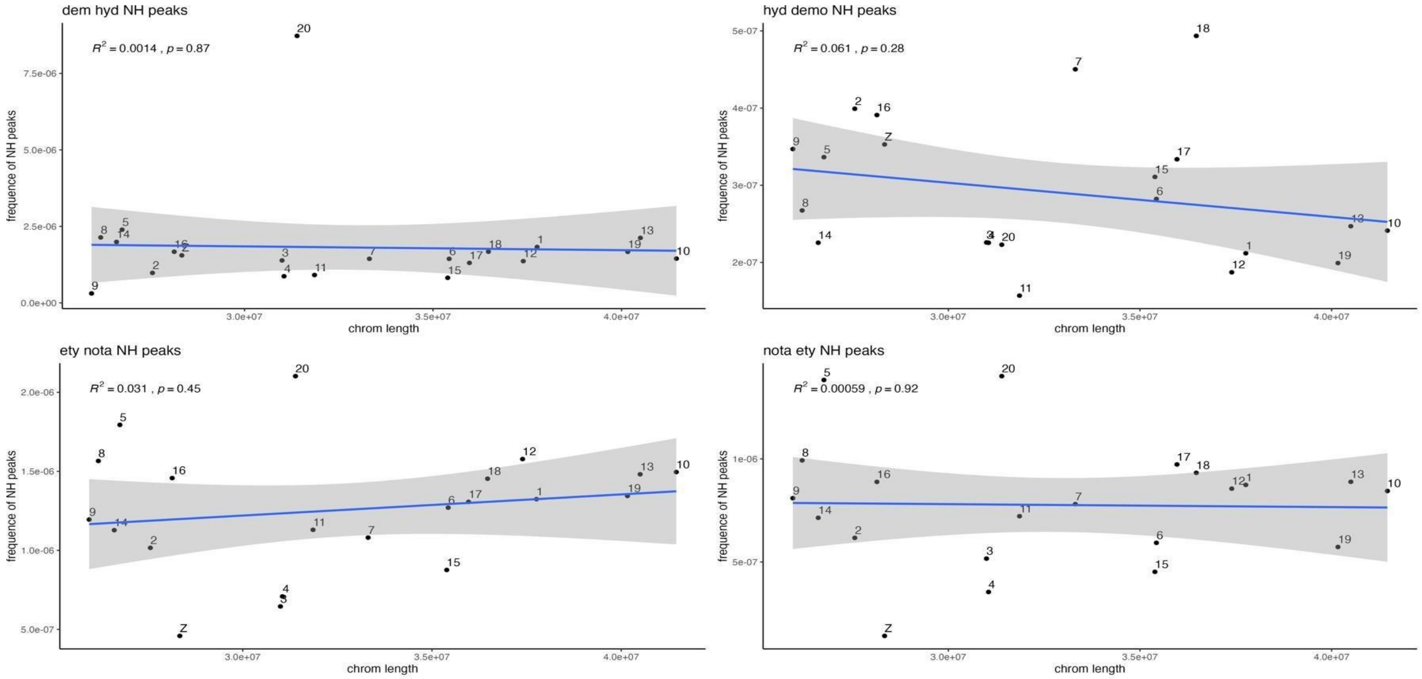

**Figure S5:** Correlation between the frequency of non-homologous ATAC-seq peaks identified in each morph and chromosome length. Each panel includes the reported R-squared values and the significance of linear regression analyses. The top-left panel represents *H. e. demophoon*, the top-right panel shows *H. e. hydara*, the bottom-left panel illustrates *H. e. etylus*, and the bottom-right panel corresponds to *H. e. notabilis*.

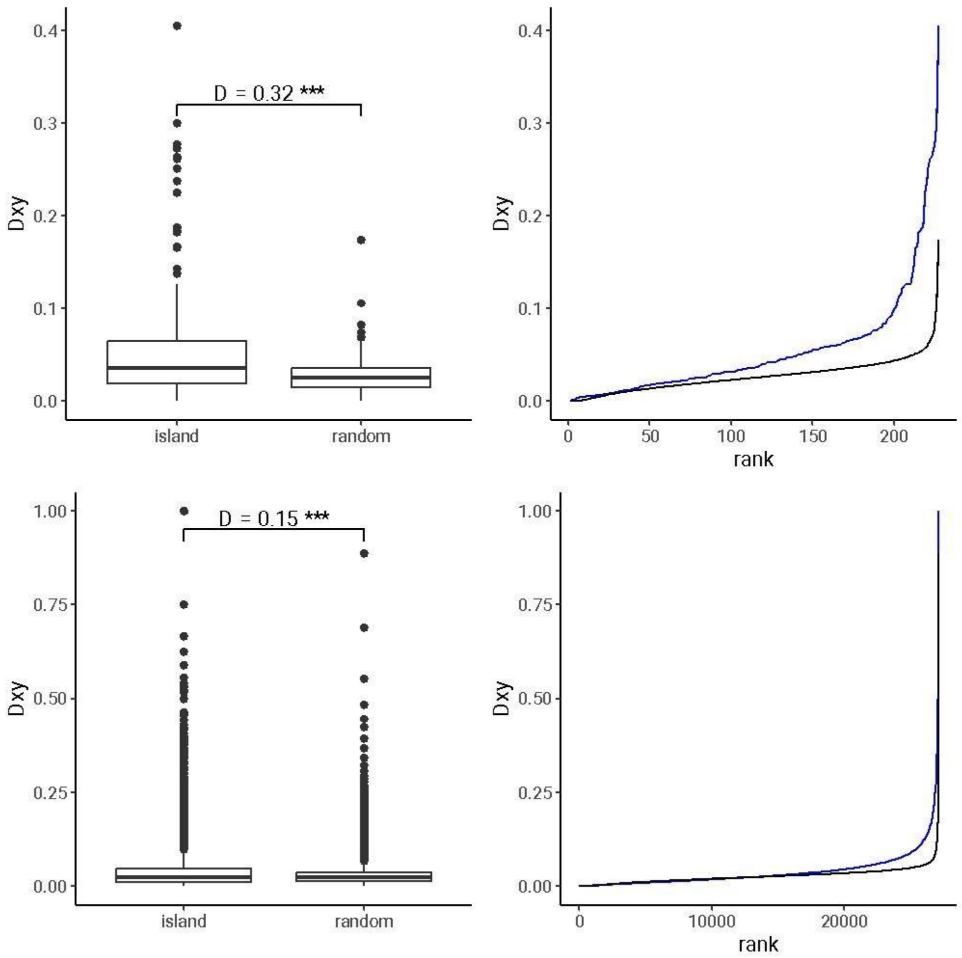

**Figure S6:** Distribution of *Dxy* (genetic diversity) in high *Fst* regions. The top-left panel shows a boxplot illustrating *Dxy* values in high *Fst* regions identified as significant in 1k-window *Fst* analysis, compared to an equal number of randomly sampled points. The top-right panel displays the distribution of their ranked values. The bottom-left panel presents a boxplot of *Dxy* values in high *Fst* regions from the overall 1k-window *Fst* data, compared to randomly sampled points. The bottom-right panel shows the distribution of their ranked values.

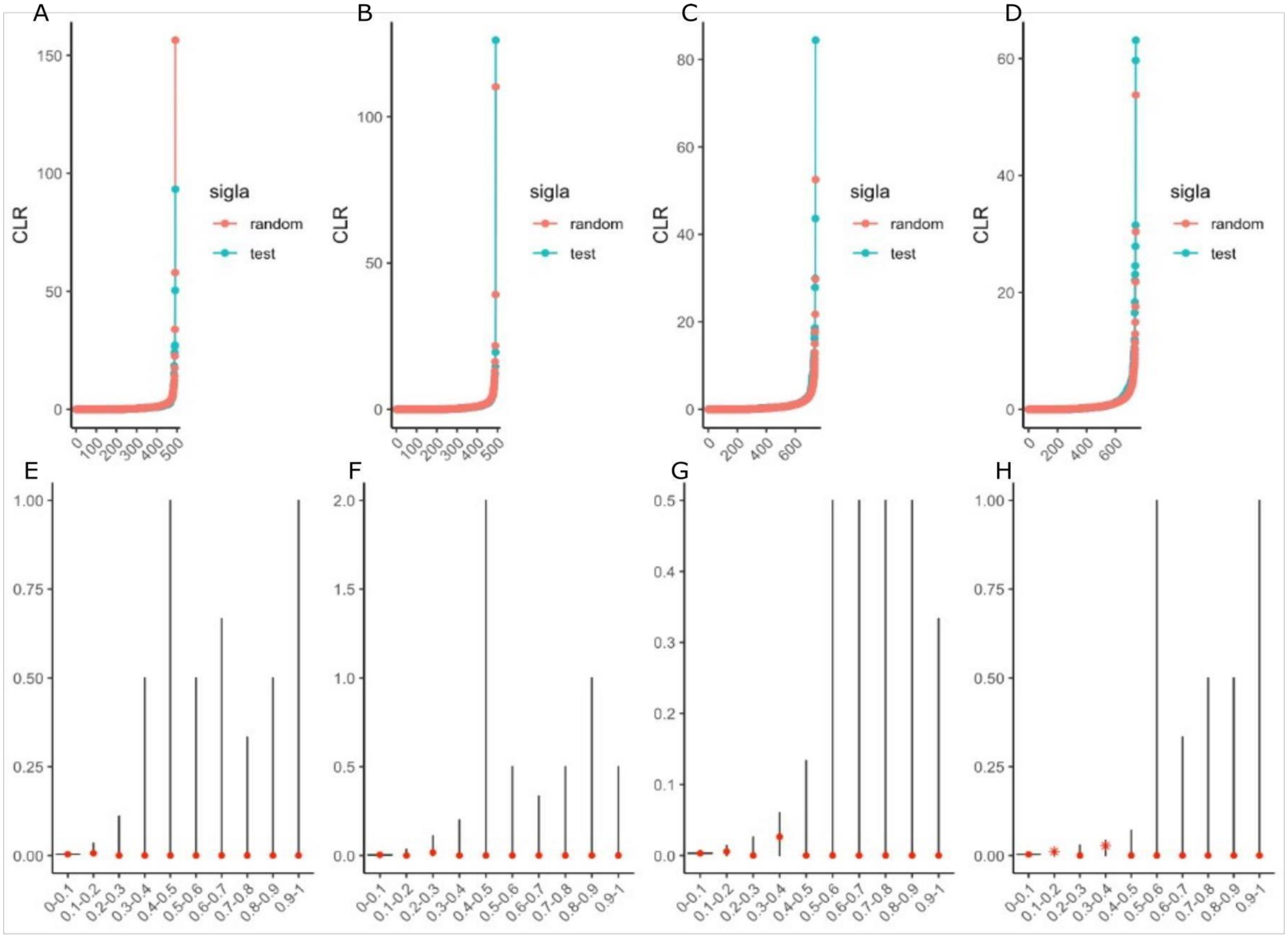

**Figure S7:** Statistical tests for selective sweep analysis. The first set of tests includes the Kolmogorov-Smirnov (KS) test performed on CLR values for different geographical morphs: (A) *H. e. demophoon*, (B) *H. e. hydara*, (C) *H. e. notabilis*, and (D) *H. e. etylus*. The second set of tests involves a binomial test applied to the same geographical morphs, denoted as (E), (F), (G), and (H), respectively.

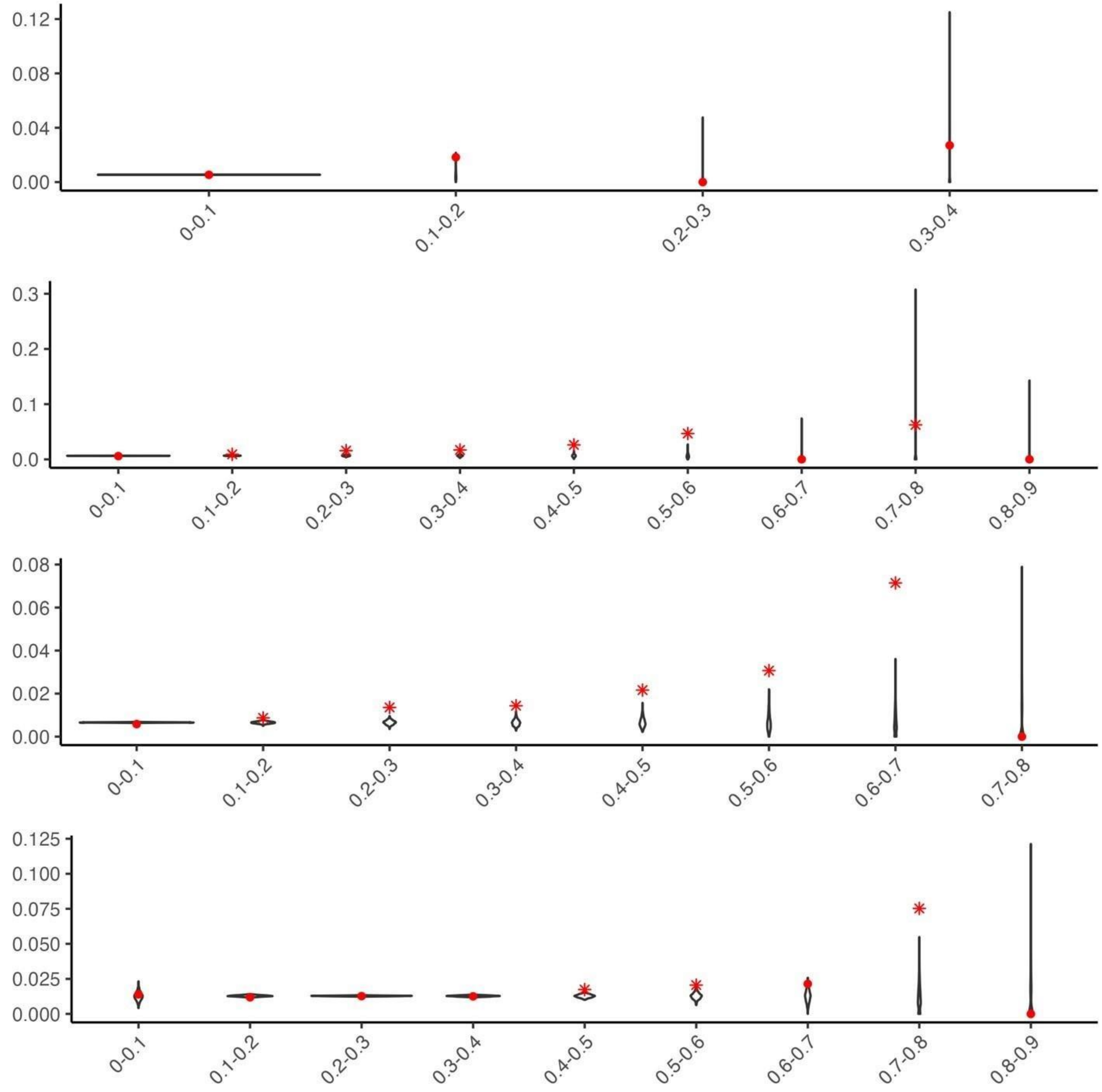

**Figure S8:** Special application of the binomial test conducted on 50k-window *Fst* values for *H. e. demophoon* compared to *H. e. hydara*, *H. e. favorinus*, *H. e. etylus*, and *H. e. chestertonii*.

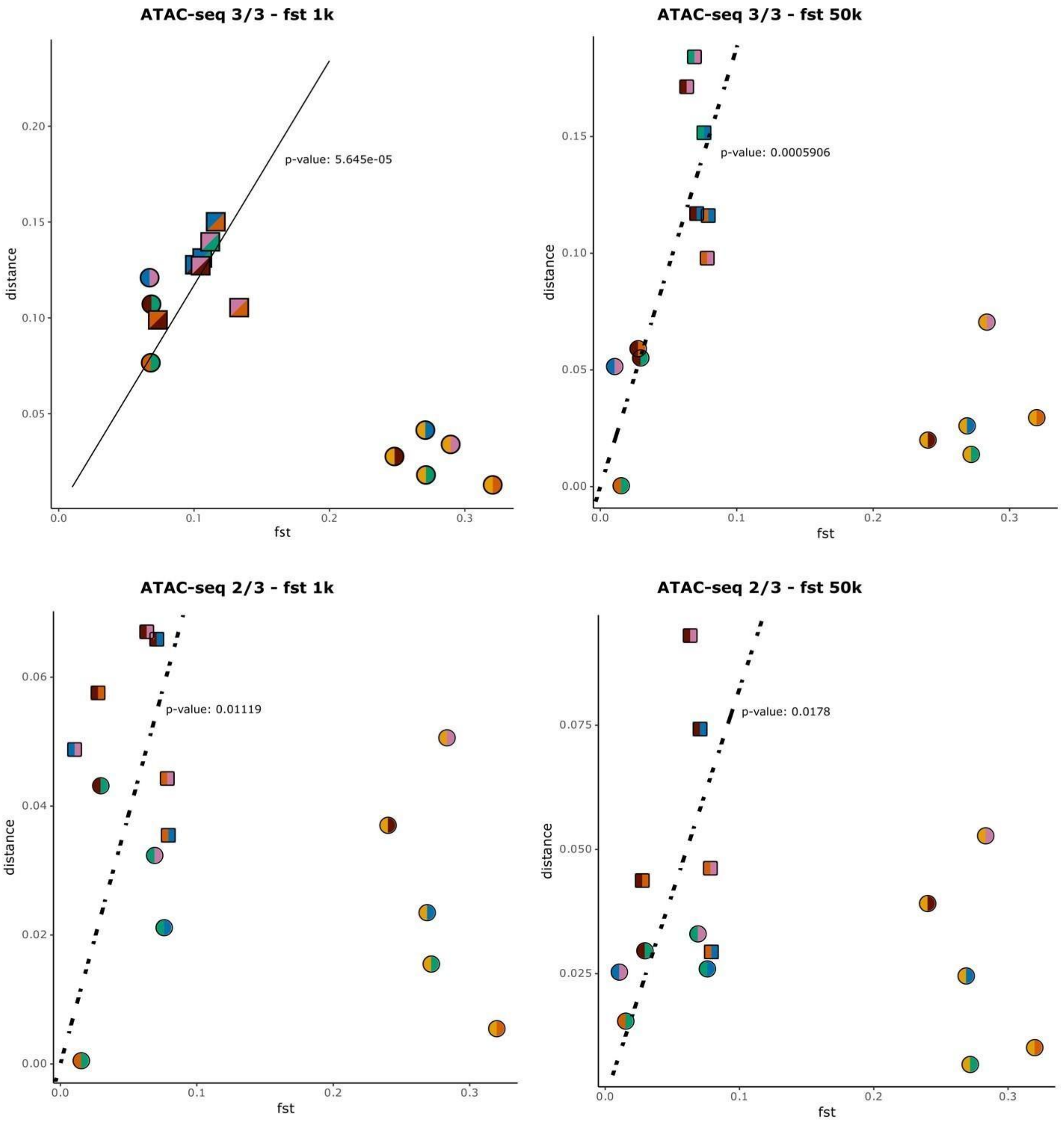

**Figure S9:** Summary of 15 pairwise Kolmogorov-Smirnov (KS) tests performed on different datasets. The top-left panel shows the comparison between ATAC-seq peaks identified in all 3 out of 3 samples and *Fst* values measured in 1k windows. The top-right panel displays the comparison between ATACseq peaks identified in all 3 out of 3 samples and *Fst* values measured in 50k windows. The bottom-left panel presents comparisons involving ATAC-seq peaks identified in at least 2 out of 3 samples with *Fst* measured in 1k windows, while the bottom-right panel shows comparisons with *Fst* measured in 50k windows. For each plot, a linear regression performed on significant comparisons, along with its p-value, is reported.

**Figure S10:** The density plot illustrates the distribution of distances from the average for the 1k-window

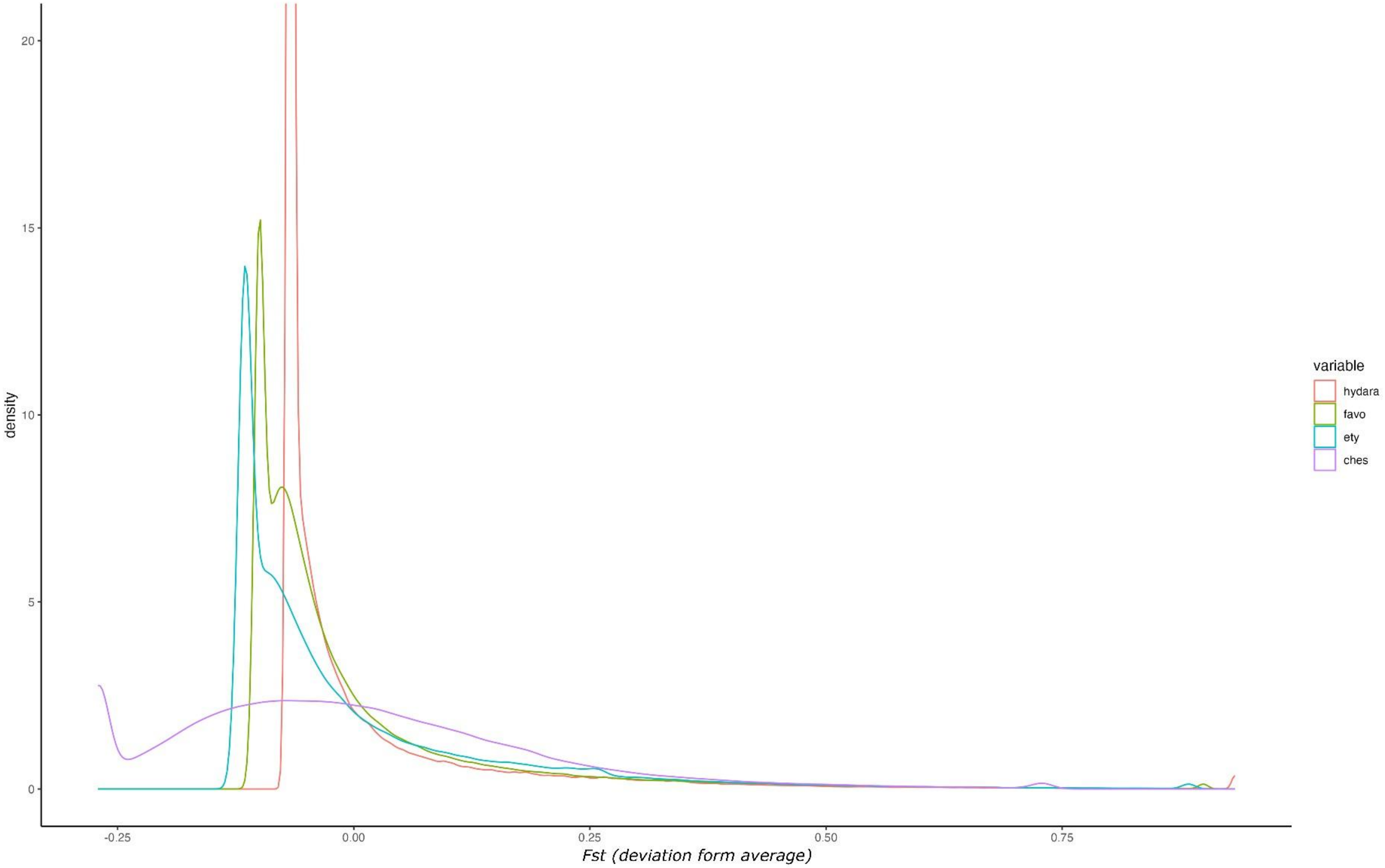

*Fst* values across *H. e. demophoon*, *H. e. hydara*, *H. e. etylus*, and *H. e. notabilis*. The x-axis represents the deviation from the mean *Fst* value for each data point within the respective populations

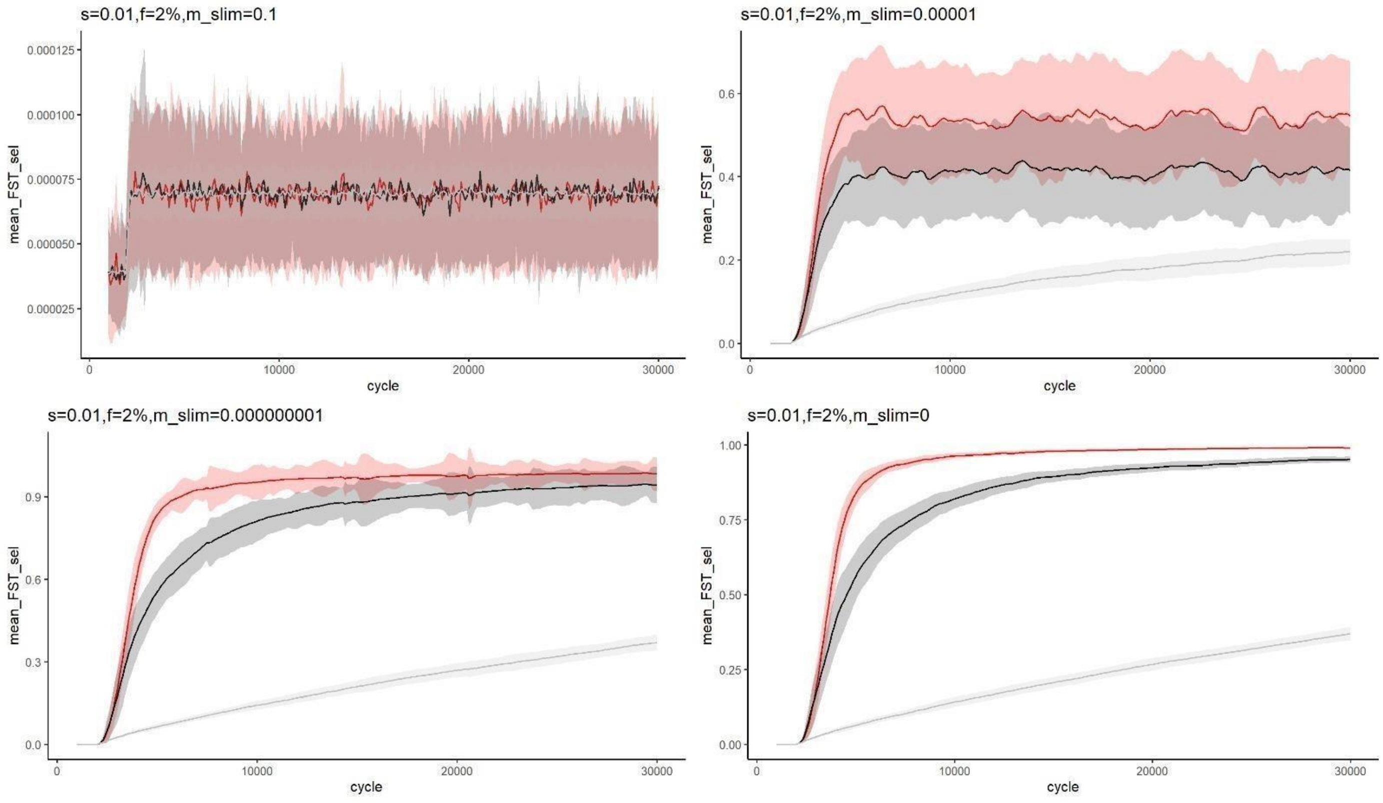

**Figure S11:** Evolution simulations with different levels of gene flow. A total of 100 simulations were performed for each panel. The red line represents *Fst* for the region under weak selection, the black line represents *Fst* for a neutral region on the same chromosome (subject to potential hitchhiking), and the gray line represents *Fst* for a completely neutral chromosome. The shaded area indicates the variance across simulations. All simulations used the same parameters, including generations and population sizes, with only the gene flow rate changing after 2,000 generations. In the top-right panel, the migration rate is 0.1; in the top-left panel, it is 0.00001; in the bottom-right panel, it is 0.000000001; and in the bottom-left panel, it is 0.

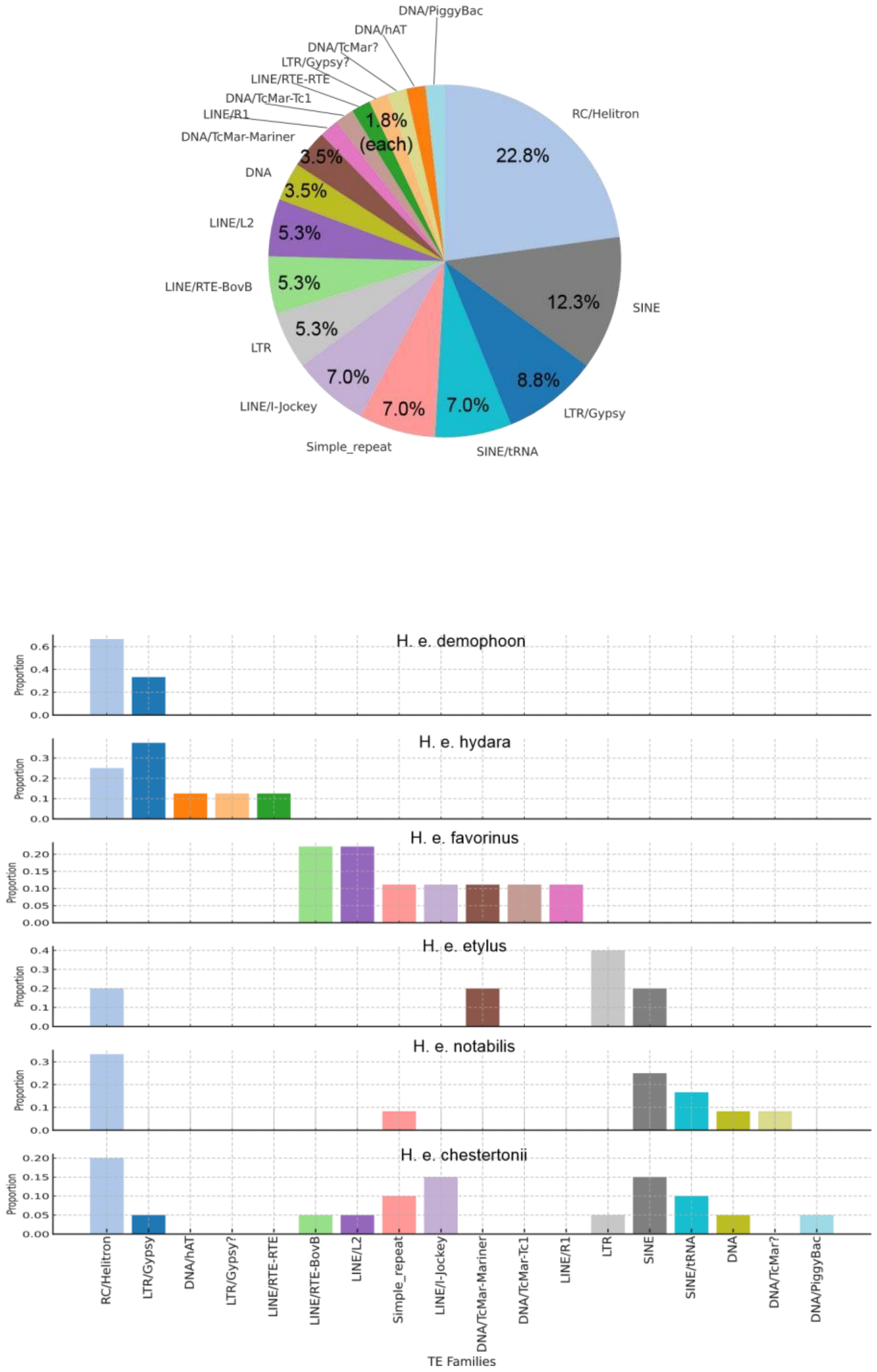

**Figure S12:** Transposable element (TE) composition of non-homologous ATAC-seq peaks. The pie chart at the top shows the overall TE composition across all identified non-homologous ATAC-seq peaks. The bottom panel displays the TE composition separated by morph.

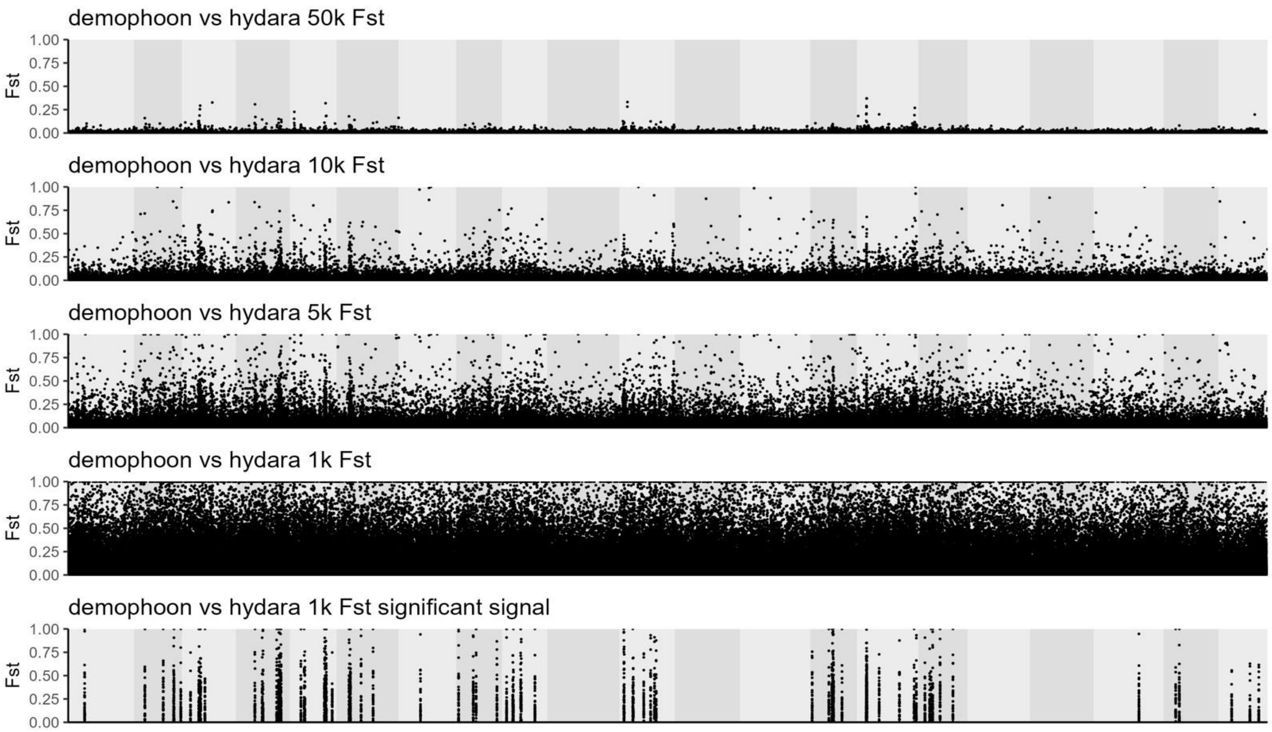

**Figure S13:** Genome-wide view of *Fst* calculated in different non-overlapping window sizes. The first panel shows genome-wide *Fst* values calculated in 50k windows, the second panel in 10k windows, the third panel in 5k windows, and the fourth panel in 1k windows. The final panel highlights only the significant regions identified in the 1k-window *Fst* analysis.

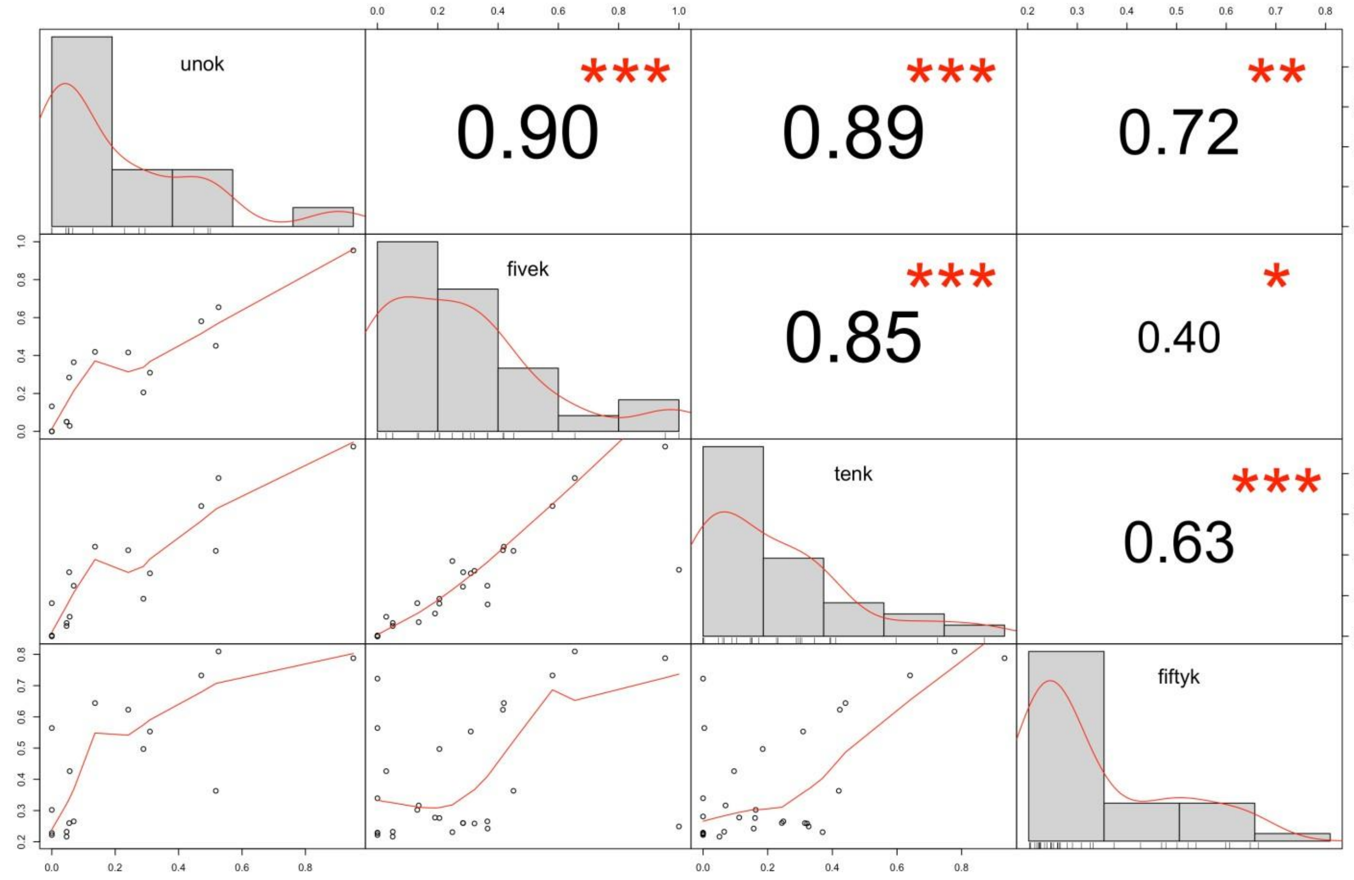

**Figure S14:** Conservation of signal among different *Fst* window sizes. This plot shows the correlation between regions with strong *Fst* values in 50k windows and the corresponding regions in *Fst* analyses using other window sizes (10k, 5k, and 1k).

### SUPPLEMENTARY TABLES

A

| pop | genome size | bp NH | NH perc % | npeaks | unique | unique % pop | confirmed NH peak |
|---|---|---|---|---|---|---|---|
| *demophoon* | 377207547 | 39762511 | 11 | 16455 | 58 | 0.35 | 3 |
| *hydara* | 356541278 | 51685917 | 14 | 16242 | 33 | 0.20 | 8 |
| *favorinus* | 363693562 | 39369963 | 11 | 15280 | 29 | 0.19 | 11 |
| *etylus* | 336746453 | 32075924 | 10 | 18811 | 52 | 0.28 | 3 |
| *notabilis* | 338400759 | 43070876 | 13 | 13165 | 4 | 0.03 | 5 |
| *chestertonii* | 326196349 | 50560864 | 16 | 22804 | 156 | 0.68 | 10 |

B

| total | shared | shared % | unique | unique % |
|---|---|---|---|---|
| 26291 | 15036 | 57.19 | 332 | 1.26 |

TableS1: population general stats. A) summary of genomic and ATAC-seq information per population. B) general summary on ATAC-seq pooling all the pop.

**demophoon vs hydara genes in Fst signals**

| | |
|---|---|
| evm.TU.Herato0206.31 | lysosomal acid glucosylceramidase-like |
| evm.TU.Herato0206.32 | farnesol dehydrogenase-like |
| evm.TU.Herato0211.98 | uncharacterized protein LOC112048264 |
| evm.TU.Herato0211.99 | RNA-directed DNA polymerase from mobile element jockey-like isoform X1 |
| evm.TU.Herato0215.128_evm.TU.Herato0215.129 | phenoloxidase subunit 1 |
| evm.TU.Herato0215.130 | phenoloxidase subunit 1 |
| evm.TU.Herato0215.43 | uncharacterized protein LOC117985158 |
| evm.TU.Herato0215.44 | fatty acid synthase-like |
| evm.TU.Herato0215.45 | uncharacterized protein LOC117985158 |
| evm.TU.Herato0301.47 | rootletin |
| evm.TU.Herato0309.1 | Nucleic-acid-binding protein from transposon X-element |
| evm.TU.Herato0412.1 | uncharacterized protein LOC113402078 isoform X2 |
| evm.TU.Herato0412.2 | ---NA--- |
| evm.TU.Herato0413.1 | Retrovirus-related Pol polyprotein from transposon opus-like Protein |
| evm.TU.Herato0413.2 | RNA-directed DNA polymerase from mobile element jockey |
| evm.TU.Herato0503.111 | carboxylesterase 1C |
| evm.TU.Herato0503.73 | beta-1,4-N-acetylgalactosaminyltransferase bre-4-like |
| evm.TU.Herato0507.1 | neuropeptide CCHamide-1 receptor |
| evm.TU.Herato0508.67 | uncharacterized protein LOC113394814 isoform X1 |
| evm.TU.Herato0601.102 | protein unc-13 homolog C-like |
| evm.TU.Herato0601.103 | RNA-directed DNA polymerase from mobile element jockey-like |
| evm.TU.Herato0601.104 | protein timeless homolog |
| evm.TU.Herato0601.105 | protein timeless homolog isoform X1 |
| evm.TU.Herato0701.327 | ecdysteroid 22-kinase |
| evm.TU.Herato0701.328 | piggyBac transposable element-derived protein 4-like |
| evm.TU.Herato0801.165 | Down syndrome cell adhesion molecule-like protein Dscam2 |
| evm.TU.Herato0801.17 | nephrin-like |
| evm.TU.Herato0821.65 | polyribonucleotide nucleotidyltransferase 1, mitochondrial |
| evm.TU.Herato0821.66 | 3'-5' ssDNA/RNA exonuclease TatD |

| | |
|---|---|
| evm.TU.Herato0901.158 | proteasome assembly chaperone 2 |
| evm.TU.Herato0901.159 | peroxisomal biogenesis factor 3 |
| evm.TU.Herato0901.303 | unnamed protein product |
| evm.TU.Herato0901.57 | tachykinins-like |
| evm.TU.Herato1106.1 | TRAS3 protein |
| evm.TU.Herato1106.2 | craniofacial development protein 2-like |
| evm.TU.Herato1108.401 | probable serine/threonine-protein kinase DDB_G0286465 |
| evm.TU.Herato1108.68 | probable uridine nucleosidase 2 |
| evm.TU.Herato1108.71 | gustatory receptor for sugar taste 64f-like |
| evm.TU.Herato1408.178,evm.TU.Herato1408.179 | sex peptide receptor-like isoform X2 |
| evm.TU.Herato1411.103 | dynein heavy chain 10, axonemal |
| evm.TU.Herato1505.164,evm.TU.Herato1505.165,evm.TU.Herato1505.86 | PARN |
| evm.TU.Herato1505.85 | Cortex |
| evm.TU.Herato1505.87 | dynein assembly factor 5, axonemal |
| evm.TU.Herato1512.5 | seminal fluid protein HACP057 |
| evm.TU.Herato1601.77,evm.TU.Herato1601.78 | RNA-directed DNA polymerase from mobile element jockey |
| evm.TU.Herato1603.2 | zinc finger DNA binding protein |
| evm.TU.Herato1908.17 | uncharacterized protein LOC113400081 |
| evm.TU.Herato2001.136 | L-ascorbate oxidase-like |
| evm.TU.Herato2001.137,evm.TU.Herato2001.138 | adenosine receptor A2b |
| evm.TU.Herato2101.158 | multidrug resistance protein homolog 49-like |
| evm.TU.Herato2101.159 | dendritic arbor reduction protein 1-like isoform X1 |
| evm.TU.Herato2101.388,evm.TU.Herato2101.389,evm.TU.Herato2101.390 | major facilitator superfamily domain-containing protein 6 |

**notabilis vs etylus genes in Fst signal**

| | |
|---|---|
| evm.TU.Herato0101.589 | chromodomain-helicase-DNA-binding protein 7 isoform X1 |
| evm.TU.Herato0101.590 | probable NADH dehydrogenase [ubiquinone] iron-sulfur protein 6, mitochondrial |
| evm.TU.Herato0101.591 | UPF0184 protein C9orf16 homolog |
| evm.TU.Herato0101.592 | ras-related protein Rab-14 |
| evm.TU.Herato0101.593 | carnitine O-acetyltransferase-like isoform X1 |
| evm.TU.Herato0101.594 | kinesin-associated protein 3 |
| evm.TU.Herato0206.21 | fatty acid synthase-like |
| evm.TU.Herato0209.38_evm.TU.Herato0209.39 | phenoloxidase subunit 2-like |
| evm.TU.Herato0209.40 | RNA-directed DNA polymerase from mobile element jockey-like |
| evm.TU.Herato0209.41 | RNA-directed DNA polymerase from mobile element jockey-like |
| evm.TU.Herato0209.42 | phenoloxidase subunit 2-like |
| evm.TU.Herato0214.23,novel_gene_22 | rabphilin-3A isoform X1 |
| evm.TU.Herato0215.146,evm.TU.Herato0215.147,evm.TU.Herato0215.9 | esterase FE4-like |
| evm.TU.Herato0215.7 | esterase FE4-like |
| evm.TU.Herato0215.8 | piggyBac transposable element-derived protein 2-like |
| evm.TU.Herato0301.61,evm.TU.Herato0301.62,evm.TU.Herato0301.63,novel_gene_23 | valacyclovir hydrolase-like |
| evm.TU.Herato0301.64,evm.TU.Herato0301.65 | valacyclovir hydrolase |
| evm.TU.Herato0309.2 | ---NA--- |
| evm.TU.Herato0310.1 | hypothetical protein EVAR_58729_1 |
| evm.TU.Herato0413.1 | Retrovirus-related Pol polyprotein from transposon opus-like Protein |
| evm.TU.Herato0413.2 | RNA-directed DNA polymerase from mobile element jockey |
| evm.TU.Herato0507.1 | neuropeptide CCHamide-1 receptor |
| evm.TU.Herato0605.1 | nucleic-acid-binding protein from mobile element jockey-like |
| evm.TU.Herato0606.237 | uncharacterized protein LOC113398345 isoform X2 |
| evm.TU.Herato0801.11 | structural maintenance of chromosomes protein 2 |
| evm.TU.Herato0801.12 | phytanoyl-CoA dioxygenase, peroxisomal-like |

| | |
|---|---|
| evm.TU.Herato0801.13 | uncharacterized protein LOC113397033 |
| evm.TU.Herato0801.302 | isopentenyl-diphosphate Delta-isomerase 1 |
| evm.TU.Herato0801.303 | kinesin heavy chain-like |
| evm.TU.Herato0801.87,evm.TU.Herato0801.88 | protein doublesex isoform X1 |
| evm.TU.Herato0807.1,evm.TU.Herato0807.2 | Brain-specific angiogenesis inhibitor 1 |
| evm.TU.Herato0821.63 | uncharacterized protein LOC116773344 |
| evm.TU.Herato0821.64 | ribonuclease P protein subunit rpr2 |
| evm.TU.Herato0821.65 | polyribonucleotide nucleotidyltransferase 1, mitochondrial |
| evm.TU.Herato0821.66 | 3'-5' ssDNA/RNA exonuclease TatD |
| evm.TU.Herato0901.255 | hypothetical protein OBRU01_15968 |
| evm.TU.Herato0901.256 | Retrovirus-related Pol polyprotein from transposon 412 |
| evm.TU.Herato1001.161 | WntA signaling ligand |
| evm.TU.Herato1103.1 | zinc finger DNA binding protein |
| evm.TU.Herato1104.1 | Zinc finger DNA binding protein |
| evm.TU.Herato1105.1 | ---NA--- |
| evm.TU.Herato1108.384 | nose resistant to fluoxetine protein 6-like |
| evm.TU.Herato1202.820 | protein deltex |
| evm.TU.Herato1411.45 | angiogenic factor with G patch and FHA domains 1 isoform X2 |
| evm.TU.Herato1505.54 | guanylate cyclase 32E |
| evm.TU.Herato1505.55 | galactokinase-like |
| evm.TU.Herato1507.220 | putative acetyltransferase |
| evm.TU.Herato1507.259,evm.TU.Herato1507.260,evm.TU.Herato1507.261,evm.TU.Herato1507.262 | uncharacterized protein LOC117987370 |
| evm.TU.Herato1507.347 | esterase FE4-like |
| evm.TU.Herato1601.83 | ---NA--- |
| evm.TU.Herato1603.1 | Gag-like protein |
| evm.TU.Herato1701.353_evm.TU.Herato1701.354 | lipopolysaccharide-induced tumor necrosis factor-alpha factor homolog |
| evm.TU.Herato1701.355 | lipopolysaccharide-induced tumor necrosis factor-alpha factor homolog |
| evm.TU.Herato1701.356 | lipopolysaccharide-induced tumor necrosis factor-alpha factor homolog |
| evm.TU.Herato1801.58,evm.TU.Herato1801.59,evm.TU.Herato1801.60 | SCY1-like protein 2 |
| evm.TU.Herato1801.61 | leucine repeat-rich protein |
| evm.TU.Herato1801.62 | leucine repeat-rich protein |

| | |
|---|---|
| evm.TU.Herato1801.63 | integrator complex subunit 7 |
| evm.TU.Herato1801.64 | homeobox protein SIX6-like |
| evm.TU.Herato1801.65 | epoxide hydrolase 4-like |
| evm.TU.Herato1801.66 | adhesion G protein-coupled receptor A3 |
| evm.TU.Herato1801.67 | kinesin-like protein KIF20B |
| evm.TU.Herato1801.68 | putative rabkinesin 6 |
| evm.TU.Herato1801.69_evm.TU.Herato1801.70 | pre-mRNA-splicing factor Slu7 |
| evm.TU.Herato1801.71 | sorting nexin-12 |
| evm.TU.Herato1801.74 | blood vessel epicardial substance-like |
| evm.TU.Herato1801.75 | dnaJ homolog subfamily C member 25 homolog |
| evm.TU.Herato1801.76 | DALR anticodon-binding domain-containing protein 3 |
| evm.TU.Herato1801.77 | max dimerization-like protein |
| evm.TU.Herato1801.78 | craniofacial development protein 2-like |
| evm.TU.Herato1805.60,evm.TU.Herato1805.61,evm.TU.Herato1805.62 | uncharacterized aarF domain-containing protein kinase 1 |
| evm.TU.Herato1805.63 | ubiquinone biosynthesis O-methyltransferase, mitochondrial-like |
| evm.TU.Herato1904.217 | ---NA--- |
| evm.TU.Herato1904.218 | fatty acyl-CoA reductase 1-like |
| evm.TU.Herato1904.222_evm.TU.Herato1904.223 | putative fatty acyl-CoA reductase CG5065 |
| evm.TU.Herato1904.224 | Piezo-type mechanosensitive ion channel component |
| evm.TU.Herato1904.225 | hypothetical protein EVAR_12794_1 |
| evm.TU.Herato1904.226 | hypothetical protein |
| evm.TU.Herato1904.230 | glucose dehydrogenase [FAD, quinone]-like |
| evm.TU.Herato2101.292 | Y+L amino acid transporter 2 |
| evm.TU.Herato2101.293 | uncharacterized protein LOC117995498 |
| evm.TU.Herato2101.294 | ---NA--- |
| evm.TU.Herato2101.295 | ---NA--- |
| evm.TU.Herato2101.296 | uncharacterized protein LOC117989496 |
| evm.TU.Herato2101.392 | uncharacterized protein LOC116777930 isoform X2 |
| evm.TU.Herato2101.399 | peptidoglycan recognition protein 1-like isoform X1 |

TableS2: genes withing Fst signal identified

| sample name | organism | stage | wing | filtered reads | # peaks | FRiP score |
|---|---|---|---|---|---|---|
| E3_FW | H. e. demophoon | 5th instar | forewing | 1769739 | 14771 | 0.27 |
| Not1_HW | H. e. notabilis | 5th instar | hindwing | 425629 | 5791 | 0.30 |
| LB_28 | H. e. notabilis | 5th instar | hindwing | 3647599 | 17706 | 0.46 |
| LB_16 | H. e. hydara | 5th instar | forewing | 5307641 | 22277 | 0.54 |
| LB_29 | H. e. favorinus | 5th instar | forewing | 4807567 | 20963 | 0.52 |
| LB_27 | H. e. hydara | 5th instar | forewing | 6788764 | 24672 | 0.54 |
| LB_17 | H. e. hydara | 5th instar | hindwing | 4960205 | 21625 | 0.57 |
| BR8_Favorinus_Brain_redoBR7 | H. e. favorinus | 5th instar | forewing | 1696138 | 18796 | 0.29 |
| Not2_FW | H. e. notabilis | 5th instar | forewing | 4763907 | 21362 | 0.42 |
| Not1_FW | H. e. notabilis | 5th instar | forewing | 6229449 | 24050 | 0.50 |
| Not2_FW | H. e. notabilis | 5th instar | forewing | 4763907 | 21362 | 0.42 |
| LI7_demophoon_FW | H. e. demophoon | 5th instar | hindwing | 4075717 | 28539 | 0.37 |
| LB_19 | H. e. favorinus | 5th instar | hindwing | 6359448 | 21068 | 0.56 |
| LB_21 | H. e. hydara | 5th instar | forewing | 4383080 | 27916 | 0.29 |
| LB_18 | H. e. favorinus | 5th instar | hindwing | 5422264 | 24091 | 0.38 |
| LB_30 | H. e. hydara | 5th instar | hindwing | 10360642 | 27559 | 0.56 |
| LI7_demophoon_HW | H. e. demophoon | 5th instar | forewing | 2438333 | 21901 | 0.24 |
| LB_6 | H. e. favorinus | 5th instar | forewing | 7763380 | 45087 | 0.41 |
| LB_41 | H. e. demophoon | 5th instar | forewing | 6924461 | 25476 | 0.45 |
| E_ety2_FW | H. e. etylus | 5th instar | hindwing | 13046479 | 26273 | 0.69 |
| Et4_HW | H. e. etylus | 5th instar | hindwing | 4631453 | 27294 | 0.28 |
| LB_7 | H. e. favorinus | 5th instar | forewing | 4460626 | 31410 | 0.31 |
| Et4_FW | H. e. etylus | 5th instar | hindwing | 10151822 | 31012 | 0.52 |
| LB_42 | H. e. demophoon | 5th instar | hindwing | 8606606 | 30192 | 0.49 |
| Not2_HW | H. e. notabilis | 5th instar | hindwing | 3611979 | 30479 | 0.24 |
| E_ety2_HW | H. e. etylus | 5th instar | hindwing | 13949150 | 21508 | 0.76 |
| Not2_HW | H. e. notabilis | 5th instar | hindwing | 3611979 | 30479 | 0.24 |
| E3_HW | H. e. demophoon | 5th instar | hindwing | 2923863 | 25027 | 0.21 |
| E_Not3_HW | H. e. notabilis | 5th instar | hindwing | 10549824 | 11992 | 0.64 |
| Chest2_HW | H. e. chest | 5th instar | forewing | 5340579 | 23635 | 0.39 |
| Chest2_FW | H. e. chest | 5th instar | forewing | 12035516 | 34886 | 0.49 |
| E_Ches1_FW | H. e. chest | 5th instar | hindwing | 12676179 | 30999 | 0.68 |
| Et5_HW | H. e. etylus | 5th instar | forewing | 10904324 | 49179 | 0.40 |
| LB_20 | H. e. hydara | 5th instar | forewing | 15676858 | 81833 | 0.39 |
| Et5_FW | H. e. etylus | 5th instar | forewing | 8833798 | 71982 | 0.28 |
| FW-pboy | H. e. demophoon | 5th instar | forewing | 14092769 | 64209 | 0.42 |
| E_Ches1_Hw | H. e. chest | 5th instar | hindwing | 18248292 | 52120 | 0.60 |

TableS3: Quality measumernet for ATAC-seq data used in the study

| comparison | morph | Wing | total | shared peak | shared peak % | unique peak | unique peak % | Variable peaks | Variabl |
|---|---|---|---|---|---|---|---|---|---|
| demophoon v | demophoon | FW | 15721 | 11003 | 69.99 | 124 | 0.79 | 4594 | 29.22 |
| | hydara | | 14691 | 11003 | 74.90 | 117 | 0.80 | 3571 | 24.31 |
| | demophoon | HW | 14978 | 7696 | 51.38 | 161 | 1.07 | 7121 | 47.54 |
| | hydara | | 14877 | 7696 | 51.73 | 144 | 0.97 | 7037 | 47.30 |
| notabilis vs et | notabilis | FW | 14877 | 7871 | 52.91 | 69 | 0.46 | 6937 | 46.63 |
| | etylus | | 16722 | 7871 | 47.07 | 138 | 0.83 | 8713 | 52.11 |
| | notabilis | HW | 12216 | 5997 | 49.09 | 50 | 0.41 | 6169 | 50.50 |
| | etylus | | 15655 | 5997 | 38.31 | 123 | 0.79 | 9535 | 60.91 |

Table S4: ATAC-seq numbers in hybrid zones

demophoon-hydara hybridzone

| Chr | Start | End | istance from TSS | gene ID | gene name |
|---|---|---|---|---|---|
| pan | 1241506 | 1241884 | 36048 | evm.model.Herato0101.16 | putative ATP-dependent RNA helicase DHX35 |
| pan | 1241506 | 1241884 | 36048 | evm.model.Herato0101.16 | putative ATP-dependent RNA helicase DHX35 |
| pan | 2511606 | 2512631 | 1823 | evm.model.Herato0101.32 | WD repeat-containing protein 43 |
| pan | 2608280 | 2608738 | 1749 | evm.model.Herato0101.34 | uncharacterized protein LOC113404900 isoform X1 |
| pan | 2608280 | 2608738 | 1749 | evm.model.Herato0101.34 | uncharacterized protein LOC113404900 isoform X1 |
| pan | 2748279 | 2748528 | 17639 | evm.model.Herato0101.39 | protein enabled isoform X1 |
| pan | 2748279 | 2748528 | 17639 | evm.model.Herato0101.39 | protein enabled isoform X1 |
| pan | 6366710 | 6366962 | 28181 | evm.model.Herato0101.116 | hypothetical protein EVAR_74554_1 |
| pan | 7110891 | 7111366 | -50860 | evm.model.Herato0101.120 | adhesion transmembrane protein |
| pan | 10555853 | 10556334 | -6031 | evm.model.Herato0101.165 | spherulin-2A-like |
| pan | 10555853 | 10556334 | -6031 | evm.model.Herato0101.165 | spherulin-2A-like |
| pan | 11326660 | 11327160 | 5987 | evm.model.Herato0101.188_evm.T | uncharacterized protein LOC113392023 |
| pan | 11636324 | 11636866 | 41364 | evm.model.Herato0101.199 | ---NA--- |
| pan | 11636324 | 11636866 | 41364 | evm.model.Herato0101.199 | ---NA--- |
| pan | 12304831 | 12305318 | -13754 | evm.model.Herato0101.214 | uncharacterized protein LOC115254287 |
| pan | 12304831 | 12305318 | -13754 | evm.model.Herato0101.214 | uncharacterized protein LOC115254287 |
| pan | 13633886 | 13634100 | -13361 | evm.model.Herato0101.233 | lipase 1-like |
| pan | 13641039 | 13641747 | -5961 | evm.model.Herato0101.233 | lipase 1-like |
| pan | 13641039 | 13641747 | -5961 | evm.model.Herato0101.233 | lipase 1-like |
| pan | 19246250 | 19246712 | 6357 | evm.model.Herato0101.315 | carbohydrate sulfotransferase 11 |
| pan | 19246250 | 19246712 | 6357 | evm.model.Herato0101.315 | carbohydrate sulfotransferase 11 |
| pan | 19803749 | 19804139 | -24067 | evm.model.Herato0101.328 | myosin-VIIa isoform X2 |
| pan | 19811956 | 19812548 | -32375 | evm.model.Herato0101.328 | myosin-VIIa isoform X2 |
| pan | 19812271 | 19812736 | -32626 | evm.model.Herato0101.328 | myosin-VIIa isoform X2 |
| pan | 19812271 | 19812736 | -32626 | evm.model.Herato0101.328 | myosin-VIIa isoform X2 |
| pan | 22562146 | 22562365 | 11637 | evm.model.Herato0101.454 | putative Activating molecule in BECN1-regulated autophagy protein 1 |
| pan | 23671630 | 23672039 | -1046 | evm.model.Herato0101.494 | chitin synthase chs-2-like |
| pan | 23671630 | 23672039 | -1046 | evm.model.Herato0101.494 | chitin synthase chs-2-like |
| pan | 26527099 | 26527430 | 50594 | evm.model.Herato0101.573 | programmed cell death protein 10 |
| pan | 30814989 | 30815268 | -2654 | evm.model.Herato0101.726 | integrin beta-6-like |
| pan | 32861049 | 32861503 | -2415 | evm.model.Herato0101.762 | lysosomal thioesterase PPT2 homolog |
| pan | 32947001 | 32947704 | 53776 | evm.model.Herato0101.764 | defense protein 2 |
| pan | 35205254 | 35205514 | 6074 | evm.model.Herato0101.801 | chromobox protein homolog 5-like |
| pan | 47526218 | 47526699 | 10029 | evm.model.Herato0209.49 | uncharacterized protein LOC113395223 isoform X1 |
| pan | 49470162 | 49470492 | 254 | evm.model.Herato0211.19 | gephyrin |
| pan | 51808408 | 51808761 | 8318 | evm.model.Herato0211.62 | serine-rich adhesin for platelets-like isoform X1 |
| pan | 54414211 | 54414554 | 18211 | evm.model.Herato0211.97 | very-long-chain 3-oxoacyl-CoA reductase |
| pan | 56168200 | 56168407 | 80081 | evm.model.Herato0214.21 | RILP protein |
| pan | 56725004 | 56725550 | -14583 | evm.model.Herato0214.40 | PAX-interacting protein 1-like |
| pan | 57750752 | 57751494 | -28201 | evm.model.Herato0215.1 | alpha-tocopherol transfer protein-like |
| pan | 62335562 | 62335924 | -15576 | evm.model.Herato0215.81 | hsp70-binding protein 1 |
| pan | 62335562 | 62335924 | -15576 | evm.model.Herato0215.81 | hsp70-binding protein 1 |
| pan | 62846636 | 62847280 | 13033 | evm.model.Herato0215.90 | UPF0061 protein PFL_0486-like |
| pan | 62846636 | 62847280 | 13033 | evm.model.Herato0215.90 | UPF0061 protein PFL_0486-like |
| pan | 63593567 | 63593960 | -8361 | evm.model.Herato0215.117 | required for meiotic nuclear division protein 1 homolog |
| pan | 63653005 | 63653552 | 51154 | evm.model.Herato0215.117 | required for meiotic nuclear division protein 1 homolog |
| pan | 66877779 | 66878193 | -28283 | evm.model.Herato0301.18 | phenylalanine--tRNA ligase alpha subunit |
| pan | 69956244 | 69957008 | -156764 | evm.model.Herato0301.46 | ras-related protein Rap1 |
| pan | 71048936 | 71049273 | 4620 | evm.model.Herato0301.70 | uncharacterized protein LOC112050341 |
| pan | 71048936 | 71049273 | 4620 | evm.model.Herato0301.70 | uncharacterized protein LOC112050341 |
| pan | 72024285 | 72024687 | -38253 | evm.model.Herato0301.93 | sodium channel protein Nach-like |
| pan | 73175222 | 73175617 | 45031 | evm.model.Herato0301.104 | vacuolar protein sorting-associated protein 72 homolog |

| | | | | | |
|---|---|---|---|---|---|
| pan | 77123835 | 77124491 | -35324 | evm.model.Herato0310.11 | DNA ligase 3 |
| pan | 78670499 | 78670934 | 100885 | evm.model.Herato0310.24 | ---NA--- |
| pan | 81054520 | 81055002 | -79536 | evm.model.Herato0310.51 | hypothetical protein |
| pan | 81054548 | 81055003 | -79550 | evm.model.Herato0310.51 | hypothetical protein |
| pan | 87628124 | 87628485 | 7077 | evm.model.Herato0310.157 | transmembrane protease serine 9-like |
| pan | 87636170 | 87636571 | -2377 | evm.model.Herato0310.158 | transmembrane protease serine 9-like |
| pan | 87636170 | 87636571 | -2377 | evm.model.Herato0310.158 | transmembrane protease serine 9-like |
| pan | 94181948 | 94182261 | -21239 | evm.model.Herato0310.290 | synaptic vesicle glycoprotein 2B-like |
| pan | 95540245 | 95540730 | -154 | evm.model.Herato0310.316 | synaptic vesicle 2-related protein-like |

| | | | | | |
|---|---|---|---|---|---|
| pan | 95540245 | 95540730 | -154 | evm.model.Herato0310.316 | synaptic vesicle 2-related protein-like |
| pan | 96380421 | 96380679 | 12137 | evm.model.Herato0401.2 | cytochrome P450 6B6-like |
| pan | 96380421 | 96380679 | 12137 | evm.model.Herato0401.2 | cytochrome P450 6B6-like |
| pan | 98093809 | 98094620 | 39669 | evm.model.Herato0401.15 | chondroitin sulfate synthase 1 |
| pan | 98093809 | 98094620 | 39669 | evm.model.Herato0401.15 | chondroitin sulfate synthase 1 |
| pan | 98154855 | 98155231 | 4033 | evm.model.Herato0401.16 | cuticular protein glycine-rich 20 |
| pan | 98154855 | 98155231 | 4033 | evm.model.Herato0401.16 | cuticular protein glycine-rich 20 |
| pan | 98743985 | 98744373 | 15289 | evm.model.Herato0401.23 | hypothetical protein RR46_05687 |
| pan | 100003667 | 100003988 | 78888 | evm.model.Herato0401.50 | heterogeneous nuclear ribonucleoprotein 27C isoform X4 |
| pan | 102785869 | 102786176 | 38071 | evm.model.Herato0403.10 | hypothetical protein EVAR_75896_1 |
| pan | 102899348 | 102899743 | -126208 | evm.model.Herato0403.11 | secretory carrier-associated membrane protein 5B isoform X1 |
| pan | 103661608 | 103661941 | 94308 | evm.model.Herato0404.1 | furin-like protease 2 |
| pan | 103661608 | 103661941 | 94308 | evm.model.Herato0404.1 | furin-like protease 2 |
| pan | 104068815 | 104069369 | 398 | evm.model.Herato0405.13 | bumetanide-sensitive sodium-(potassium)-chloride cotransporter |
| pan | 104068815 | 104069369 | 398 | evm.model.Herato0405.13 | bumetanide-sensitive sodium-(potassium)-chloride cotransporter |
| pan | 104247086 | 104247409 | 3584 | evm.model.Herato0405.14 | 40S ribosomal protein S15 |
| pan | 104247088 | 104247414 | 3580 | evm.model.Herato0405.14 | 40S ribosomal protein S15 |
| pan | 106281832 | 106282464 | -19881 | evm.model.Herato0405.41 | uncharacterized protein LOC117990158 |
| pan | 107823674 | 107824048 | 15332 | evm.model.Herato0411.2 | ---NA--- |
| pan | 107823712 | 107824023 | 15326 | evm.model.Herato0411.2 | ---NA--- |
| pan | 109340975 | 109341509 | 86143 | evm.model.Herato0411.15 | Gag protein |
| pan | 129056102 | 129056574 | -67680 | evm.model.Herato0501.37 | bone morphogenetic protein receptor type-1B isoform X1 |
| pan | 129081765 | 129082226 | -58369 | evm.model.Herato0501.38 | 60S ribosomal protein L34-like |
| pan | 129081765 | 129082226 | -58369 | evm.model.Herato0501.38 | 60S ribosomal protein L34-like |
| pan | 130978044 | 130978326 | 28300 | evm.model.Herato0503.21 | zinc finger-containing ubiquitin peptidase 1-like |
| pan | 130978044 | 130978326 | 28300 | evm.model.Herato0503.21 | zinc finger-containing ubiquitin peptidase 1-like |
| pan | 134305451 | 134306072 | -2380 | evm.model.Herato0503.85 | uncharacterized protein LOC112048264 |
| pan | 134583003 | 134583866 | 3683 | evm.model.Herato0503.96 | SET and MYND domain-containing protein 4 |
| pan | 136938426 | 136938819 | 2115 | evm.model.Herato0503.120 | NFU1 iron-sulfur cluster scaffold homolog, mitochondrial-like |
| pan | 137054074 | 137054321 | -91249 | evm.model.Herato0503.121 | protein spaetzle 3 |
| pan | 137107594 | 137107927 | -41708 | evm.model.Herato0503.122 | seminal fluid protein HACP011 |
| pan | 138265958 | 138266498 | 36585 | evm.model.Herato0503.131 | cGMP-dependent protein kinase, isozyme 2 forms cD5/T2-like isoform X1 |
| pan | 138265958 | 138266498 | 36585 | evm.model.Herato0503.131 | cGMP-dependent protein kinase, isozyme 2 forms cD5/T2-like isoform X1 |
| pan | 138829669 | 138829977 | 67480 | evm.model.Herato0503.132 | rap1 GTPase-activating protein 1 isoform X1 |
| pan | 142072489 | 142073336 | -1030 | evm.model.Herato0503.221 | ADP-ribosylation factor 2 |
| pan | 142151959 | 142152432 | -50081 | evm.model.Herato0503.225 | uncharacterized protein LOC113400287 |
| pan | 143035638 | 143036162 | -20815 | evm.model.Herato0503.241 | estradiol 17-beta-dehydrogenase 11-like |
| pan | 143055548 | 143055830 | -1026 | evm.model.Herato0503.241 | estradiol 17-beta-dehydrogenase 11-like |
| pan | 143346959 | 143347536 | -1631 | evm.model.Herato0503.250 | potassium channel AKT3-like |

| | | | | | |
|---|---|---|---|---|---|
| pan | 144529078 | 144529428 | 742 | evm.model.Herato0503.286 | uncharacterized protein LOC111354683 |
| pan | 144529090 | 144529429 | 748 | evm.model.Herato0503.286 | uncharacterized protein LOC111354683 |
| pan | 144533595 | 144534242 | 967 | evm.model.Herato0503.287 | uncharacterized protein LOC111354683 |
| pan | 144541714 | 144542076 | 8944 | evm.model.Herato0503.287 | uncharacterized protein LOC111354683 |
| pan | 144541720 | 144542048 | 8933 | evm.model.Herato0503.287 | uncharacterized protein LOC111354683 |
| pan | 144572306 | 144572978 | -10491 | evm.model.Herato0503.288 | early boundary activity protein 2-like isoform X2 |
| pan | 148425966 | 148426333 | -33852 | evm.model.Herato0508.4 | glyoxylate reductase/hydroxypyruvate reductase-like |
| pan | 153971000 | 153971493 | 15281 | evm.model.Herato0511.7 | hypothetical protein B5V51_8296 |
| pan | 153985020 | 153985697 | -17371 | evm.model.Herato0511.8 | potassium/sodium hyperpolarization-activated cyclic nucleotide-gated channel 4-like isoform X15 |
| pan | 154071791 | 154072116 | -25023 | evm.model.Herato0511.9 | unnamed protein product |
| pan | 165659490 | 165659690 | -141200 | evm.model.Herato0606.49 | pyruvate kinase-like |
| pan | 166141236 | 166141659 | 1321 | evm.model.Herato0606.54 | uncharacterized protein LOC117987079 isoform X3 |
| pan | 166568008 | 166568376 | 120 | evm.model.Herato0606.58 | dnaJ homolog subfamily C member 8 |
| pan | 168523255 | 168523563 | -2682 | evm.model.Herato0606.96 | protein spindle-F |
| pan | 169922458 | 169923234 | -9094 | evm.model.Herato0606.111 | ---NA--- |
| pan | 169922458 | 169923234 | -9094 | evm.model.Herato0606.111 | ---NA--- |
| pan | 170387157 | 170387467 | -20961 | evm.model.Herato0606.122 | trypsin-1-like |
| pan | 170387157 | 170387467 | -20961 | evm.model.Herato0606.122 | trypsin-1-like |
| pan | 170402569 | 170402908 | -36387 | evm.model.Herato0606.122 | trypsin-1-like |
| pan | 171001958 | 171002164 | -19160 | evm.model.Herato0606.132 | barH-like 2 homeobox protein |
| pan | 173821098 | 173821656 | -486 | evm.model.Herato0606.204 | uncharacterized protein C15orf61 isoform X1 |
| pan | 175729386 | 175729609 | 37645 | evm.model.Herato0606.240 | Neuronal acetylcholine receptor subunit alpha-3 |
| pan | 179446428 | 179446683 | 26090 | evm.model.Herato0606.284 | uncharacterized protein LOC116766716 isoform X2 |
| pan | 179677813 | 179678327 | -10412 | evm.model.Herato0606.291 | phospholipase B1, membrane-associated-like |
| pan | 179677813 | 179678327 | -10412 | evm.model.Herato0606.291 | phospholipase B1, membrane-associated-like |
| pan | 179928512 | 179928894 | 350 | evm.model.Herato0606.302 | neurexin-4 isoform X1 |
| pan | 180612690 | 180613172 | -158677 | evm.model.Herato0606.306 | thioredoxin-2 |
| pan | 180612692 | 180613182 | -158671 | evm.model.Herato0606.306 | thioredoxin-2 |
| pan | 181967803 | 181968190 | 12679 | evm.model.Herato0606.331 | Copper chaperone for superoxide dismutase |
| pan | 181967803 | 181968190 | 12679 | evm.model.Herato0606.331 | Copper chaperone for superoxide dismutase |
| pan | 183132152 | 183132966 | 3878 | evm.model.Herato0606.374 | Replicase large subunit |
| pan | 183132152 | 183132966 | 3878 | evm.model.Herato0606.374 | Replicase large subunit |
| pan | 185237320 | 185237710 | -24212 | evm.model.Herato0606.451 | cyclin G |
| pan | 185237320 | 185237710 | -24212 | evm.model.Herato0606.451 | cyclin G |
| pan | 186717420 | 186717773 | 2827 | evm.model.Herato0608.5 | zinc finger BED domain-containing protein 5-like |
| pan | 186717420 | 186717773 | 2827 | evm.model.Herato0608.5 | zinc finger BED domain-containing protein 5-like |
| pan | 189326880 | 189327616 | 38617 | evm.model.Herato0609.84 | unnamed protein product |
| pan | 189326880 | 189327616 | 38617 | evm.model.Herato0609.84 | unnamed protein product |
| pan | 189515688 | 189516421 | 9902 | evm.model.Herato0609.90 | unnamed protein product |
| pan | 189515702 | 189516367 | 9922 | evm.model.Herato0609.90 | unnamed protein product |
| pan | 191678496 | 191679598 | 5958 | evm.model.Herato0701.46 | pyruvate dehydrogenase (acetyl-transferring) kinase, mitochondrial |
| pan | 196135262 | 196135912 | -6567 | evm.model.Herato0701.183 | Disks large-associated protein 5 |
| pan | 196135262 | 196135912 | -6567 | evm.model.Herato0701.183 | Disks large-associated protein 5 |
| pan | 198618200 | 198618573 | -4135 | evm.model.Herato0701.244 | uncharacterized protein LOC112043869 |
| pan | 198618200 | 198618573 | -4135 | evm.model.Herato0701.244 | uncharacterized protein LOC112043869 |
| pan | 200591068 | 200591521 | -62103 | evm.model.Herato0701.283 | E3 ubiquitin-protein ligase ZNRF1 |
| pan | 200591068 | 200591521 | -62103 | evm.model.Herato0701.283 | E3 ubiquitin-protein ligase ZNRF1 |
| pan | 200648053 | 200648308 | 82199 | evm.model.Herato0701.284 | E3 ubiquitin-protein ligase ZNRF1 |
| pan | 200648053 | 200648308 | 82199 | evm.model.Herato0701.284 | E3 ubiquitin-protein ligase ZNRF1 |
| pan | 200739506 | 200739871 | -7417 | evm.model.Herato0701.286 | TATA element modulatory factor |
| pan | 203549522 | 203549807 | -82347 | novel_exon_70 | transposase |
| pan | 206324110 | 206324639 | 36767 | evm.model.Herato0701.400 | supervillin-like isoform X4 |
| pan | 206324110 | 206324639 | 36767 | evm.model.Herato0701.400 | supervillin-like isoform X4 |

| | | | | | |
|---|---|---|---|---|---|
| pan | 208062540 | 208062992 | 5726 | evm.model.Herato0701.447 | ataxin-1-like isoform X1 |
| pan | 208062540 | 208062992 | 5726 | evm.model.Herato0701.447 | ataxin-1-like isoform X1 |
| pan | 208772495 | 208772960 | -8832 | evm.model.Herato0701.470 | zinc finger CCCH domain-containing protein 18 isoform X1 |
| pan | 209401366 | 209401692 | -16 | evm.model.Herato0701.495 | low-density lipoprotein receptor class A domaincontaining protein 4 isoform X2 |
| pan | 209401366 | 209401692 | -16 | evm.model.Herato0701.495 | low-density lipoprotein receptor class A domaincontaining protein 4 isoform X2 |
| pan | 209980449 | 209980822 | 11445 | evm.model.Herato0701.521 | xylosyltransferase oxt |
| pan | 210215322 | 210215934 | 16430 | evm.model.Herato0701.539 | putative Phosphatidylinositol 3-and 4-kinase |
| pan | 211098790 | 211099517 | -14886 | evm.model.Herato0701.583 | hypothetical protein EVAR_58729_1 |
| pan | 211098790 | 211099517 | -14886 | evm.model.Herato0701.583 | hypothetical protein EVAR_58729_1 |
| pan | 217623337 | 217623821 | -66657 | evm.model.Herato0701.693 | opsin |
| pan | 220381348 | 220381943 | -48747 | evm.model.Herato0701.740_evm.The | pre-mRNA-splicing factor ATP-dependent RNA licase PRP16 |
| pan | 220381348 | 220381943 | -48747 | evm.model.Herato0701.740_evm.The | pre-mRNA-splicing factor ATP-dependent RNA licase PRP16 |
| pan | 221848880 | 221849398 | -98 | evm.model.Herato0701.757 | longitudinals lacking protein-like isoform X1 |
| pan | 221848880 | 221849398 | -98 | evm.model.Herato0701.757 | longitudinals lacking protein-like isoform X1 |
| pan | 222463898 | 222464857 | -31030 | evm.model.Herato0701.762 | uncharacterized protein LOC113514709 |
| pan | 228329720 | 228330939 | -500 | evm.model.Herato0801.46 | zinc transporter 7 |
| pan | 228329720 | 228330939 | -500 | evm.model.Herato0801.46 | zinc transporter 7 |
| pan | 229423451 | 229424024 | 13396 | evm.model.Herato0801.66 | spastin isoform X3 |
| pan | 229423451 | 229424024 | 13396 | evm.model.Herato0801.66 | spastin isoform X3 |
| pan | 229601477 | 229601797 | 9270 | evm.model.Herato0801.72 | IQ domain-containing protein G-like |
| pan | 231637346 | 231637664 | 23680 | evm.model.Herato0801.101 | chromosome-associated kinesin KIF4 |
| pan | 234890766 | 234891182 | 9263 | evm.model.Herato0801.181 | uncharacterized protein LOC117990688 |
| pan | 238784046 | 238785966 | -706 | evm.model.Herato0801.280 | caspase-8 |
| pan | 241972392 | 241972956 | -140427 | evm.model.Herato0813.1 | uncharacterized protein LOC115455229, partial |
| pan | 242078846 | 242079350 | -34003 | evm.model.Herato0813.1 | uncharacterized protein LOC115455229, partial |
| pan | 247207232 | 247207550 | -7274 | evm.model.Herato0821.85 | decaprenyl-diphosphate synthase subunit 2-like |
| pan | 247207232 | 247207550 | -7274 | evm.model.Herato0821.85 | decaprenyl-diphosphate synthase subunit 2-like |
| pan | 248979627 | 248980391 | -20158 | evm.model.Herato0821.132 | protein serrate |
| pan | 249036381 | 249036811 | -3903 | novel_exon_96 | RNA-directed DNA polymerase from mobile element jockey-like |
| pan | 249036381 | 249036811 | -3903 | novel_exon_96 | RNA-directed DNA polymerase from mobile element jockey-like |
| pan | 257647352 | 257647926 | 51260 | evm.model.Herato0901.104_evm.T | 39S ribosomal protein L19, mitochondrial |
| pan | 266181301 | 266182399 | -15141 | evm.model.Herato0901.275 | trithorax group protein osa isoform X5 |
| pan | 266181301 | 266182399 | -15141 | evm.model.Herato0901.275 | trithorax group protein osa isoform X5 |
| pan | 266517511 | 266518328 | -19596 | evm.model.Herato0901.280 | another transcription unit protein |
| pan | 270207223 | 270207443 | 44475 | evm.model.Herato0901.324 | dystrophin, isoforms A/C/F/G/H-like |
| pan | 273159097 | 273159999 | -104127 | evm.model.Herato0901.372 | neural cell adhesion molecule 1-like isoform X2 |
| pan | 274572235 | 274572556 | -1658 | evm.model.Herato0904.20 | endonuclease-reverse transcriptase |
| pan | 274572235 | 274572556 | -1658 | evm.model.Herato0904.20 | endonuclease-reverse transcriptase |
| pan | 274871750 | 274872172 | -58398 | evm.model.Herato0904.25 | UPF0160 protein MYG1, mitochondrial isoform X2 |
| pan | 275702685 | 275702924 | -15188 | evm.model.Herato1001.23 | ---NA--- |
| pan | 275763724 | 275763973 | -8977 | evm.model.Herato1001.26 | tektin-4-like |
| pan | 275810761 | 275811045 | 30687 | evm.model.Herato1001.27 | ribosomal protein S6 kinase delta-1-like isoform X2 |
| pan | 276230379 | 276231367 | 33212 | evm.model.Herato1001.39 | facilitated trehalose transporter Tret1-like |
| pan | 276230379 | 276231367 | 33212 | evm.model.Herato1001.39 | facilitated trehalose transporter Tret1-like |
| pan | 279901560 | 279901838 | -26098 | evm.model.Herato1001.106 | rho guanine nucleotide exchange factor 18 isoform X4 |
| pan | 287085602 | 287085877 | -1130 | evm.model.Herato1003.9 | serine protease inhibitor 2.1-like |
| pan | 287085602 | 287085877 | -1130 | evm.model.Herato1003.9 | serine protease inhibitor 2.1-like |
| pan | 288123519 | 288123920 | -18403 | evm.model.Herato1003.23 | zinc transporter 2-like |

| | | | | | |
|---|---|---|---|---|---|
| pan | 288344581 | 288346175 | -703 | evm.model.Herato1003.31 | chymotrypsin-like elastase family member 2A |
| pan | 288344581 | 288346175 | -703 | evm.model.Herato1003.31 | chymotrypsin-like elastase family member 2A |
| pan | 290176710 | 290177364 | -565 | evm.model.Herato1003.65 | chymotrypsin-like elastase family member 2A |
| pan | 290176710 | 290177364 | -565 | evm.model.Herato1003.65 | chymotrypsin-like elastase family member 2A |
| pan | 290198459 | 290198631 | 17234 | evm.model.Herato1003.66 | chymotrypsin-like elastase family member 2A |
| pan | 293748245 | 293749176 | 53589 | evm.model.Herato1003.114 | reverse transcriptase |
| pan | 293748245 | 293749176 | 53589 | evm.model.Herato1003.114 | reverse transcriptase |
| pan | 295892250 | 295892495 | 16996 | evm.model.Herato1003.171 | ras-related protein Rab6 isoform X1 |
| pan | 295892250 | 295892495 | 16996 | evm.model.Herato1003.171 | ras-related protein Rab6 isoform X1 |
| pan | 299148843 | 299149861 | 8432 | evm.model.Herato1003.265 | UPF0489 protein C5orf22 homolog |
| pan | 299148847 | 299149826 | 8416 | evm.model.Herato1003.265 | UPF0489 protein C5orf22 homolog |
| pan | 301297997 | 301298483 | -68 | novel_exon_116 | |
| pan | 302020652 | 302020865 | -5959 | evm.model.Herato1005.84 | agrin-like |
| pan | 302020652 | 302020865 | -5959 | evm.model.Herato1005.84 | agrin-like |
| pan | 306304914 | 306305277 | 5124 | evm.model.Herato1005.202 | Kinetochore-associated protein 1 |
| pan | 306304990 | 306305275 | 5087 | evm.model.Herato1005.202 | Kinetochore-associated protein 1 |
| pan | 309928335 | 309928599 | 64787 | evm.model.Herato1007.84 | chitin deacetylase 5 isoform X6 |
| pan | 309928356 | 309928589 | 64792 | evm.model.Herato1007.84 | chitin deacetylase 5 isoform X6 |
| pan | 313292844 | 313293267 | -29584 | evm.model.Herato1007.146 | uncharacterized LOC106124082 |
| pan | 313675458 | 313675739 | 22072 | evm.model.Herato1007.171 | immunoglobulin-binding protein 1 |
| pan | 313675458 | 313675739 | 22072 | evm.model.Herato1007.171 | immunoglobulin-binding protein 1 |
| pan | 314142975 | 314143586 | 24561 | evm.model.Herato1007.182 | uncharacterized MFS-type transporter C09D4.1 isoform X2 |
| pan | 314264747 | 314265101 | -136 | evm.model.Herato1007.183 | obscurin isoform X3 |
| pan | 327093197 | 327093653 | 2915 | evm.model.Herato1108.138 | steroid receptor RNA activator 1-like |
| pan | 327642753 | 327643340 | 67770 | evm.model.Herato1108.153 | YTH domain-containing family protein 3 isoform X1 |
| pan | 330459216 | 330459453 | -115249 | evm.model.Herato1108.241 | filamin-A isoform X1 |
| pan | 330459216 | 330459453 | -115249 | evm.model.Herato1108.241 | filamin-A isoform X1 |
| pan | 331868953 | 331869523 | -66 | evm.model.Herato1108.271 | transcription termination factor 5, mitochondrial-like |
| pan | 331868953 | 331869523 | -66 | evm.model.Herato1108.271 | transcription termination factor 5, mitochondrial-like |
| pan | 336972106 | 336972311 | 781 | evm.model.Herato1108.386 | ---NA--- |
| pan | 337364581 | 337364838 | -9668 | evm.model.Herato1108.396 | hypothetical protein RR48_11784 |
| pan | 337364581 | 337364838 | -9668 | evm.model.Herato1108.396 | hypothetical protein RR48_11784 |
| pan | 337761254 | 337761507 | -2064 | evm.model.Herato1108.406 | UPF0676 protein C1494.01-like |
| pan | 337761254 | 337761507 | -2064 | evm.model.Herato1108.406 | UPF0676 protein C1494.01-like |
| pan | 337849517 | 337849917 | -349 | evm.model.Herato1108.414 | general transcription factor 3C polypeptide 6-like isoform X2 |
| pan | 338815437 | 338815880 | 26212 | evm.model.Herato1108.449 | probable Ras GTPase-activating protein isoform X6 |
| pan | 346391414 | 346391847 | 47235 | evm.model.Herato1108.603 | protein crumbs |
| pan | 346391414 | 346391847 | 47235 | evm.model.Herato1108.603 | protein crumbs |
| pan | 349036462 | 349037217 | 56287 | evm.model.Herato1201.11 | transmembrane protease serine 11G-like |
| pan | 350177622 | 350179604 | 11370 | evm.model.Herato1202.25 | serine protease 55-like |
| pan | 350177622 | 350179604 | 11370 | evm.model.Herato1202.25 | serine protease 55-like |
| pan | 352425307 | 352425696 | -2476 | evm.model.Herato1202.59 | ethanolaminephosphotransferase 1-like |
| pan | 352470182 | 352470648 | 10765 | evm.model.Herato1202.60 | partitioning defective protein 6 |
| pan | 352470182 | 352470648 | 10765 | evm.model.Herato1202.60 | partitioning defective protein 6 |
| pan | 352507230 | 352507799 | -679 | evm.model.Herato1202.63 | putative kakapo |
| pan | 352681969 | 352682148 | 167189 | novel_exon_140 | Transposon Ty3-G Gag-Pol polyprotein |
| pan | 353972632 | 353972965 | -75263 | evm.model.Herato1202.86 | trypsin-5-like isoform X1 |
| pan | 355647347 | 355648019 | -2281 | evm.model.Herato1202.123 | focal adhesion kinase 1 isoform X1 |
| pan | 355647347 | 355648019 | -2281 | evm.model.Herato1202.123 | focal adhesion kinase 1 isoform X1 |
| pan | 355928038 | 355928225 | 6 | evm.model.Herato1202.130 | probable sodium-coupled neutral amino acid transporter 6 |
| pan | 356755118 | 356755367 | 5810 | evm.model.Herato1202.151 | phosphoenolpyruvate carboxykinase [GTP]-like isoform X1 |

| | | | | | |
|---|---|---|---|---|---|
| pan | 358550536 | 358551004 | -46118 | evm.model.Herato1202.216 | sodium- and chloride-dependent transporter XTRP3 isoform X1 |
| pan | 358550536 | 358551004 | -46118 | evm.model.Herato1202.216 | sodium- and chloride-dependent transporter XTRP3 isoform X1 |
| pan | 358755460 | 358755741 | -14594 | evm.model.Herato1202.225 | GMP reductase 1-like |
| pan | 358755460 | 358755741 | -14594 | evm.model.Herato1202.225 | GMP reductase 1-like |
| pan | 358807971 | 358808246 | -12427 | evm.model.Herato1202.226 | pacifastin light chain |
| pan | 358845437 | 358845926 | -19960 | evm.model.Herato1202.227 | protein draper-like isoform X1 |
| pan | 358845437 | 358845926 | -19960 | evm.model.Herato1202.227 | protein draper-like isoform X1 |
| pan | 359284374 | 359284786 | -3870 | evm.model.Herato1202.247 | KH domain-containing, RNA-binding, signal transduction-associated protein 2-like isoform X1 |
| pan | 359284374 | 359284786 | -3870 | evm.model.Herato1202.247 | KH domain-containing, RNA-binding, signal transduction-associated protein 2-like isoform X1 |
| pan | 367184881 | 367185848 | 54898 | evm.model.Herato1202.407 | calmodulin-binding transcription activator 2 |
| pan | 367184905 | 367185853 | 54913 | evm.model.Herato1202.407 | calmodulin-binding transcription activator 2 |
| pan | 368967223 | 368967942 | 10236 | evm.model.Herato1202.439 | COP9 signalosome complex subunit 7b |
| pan | 369554357 | 369554715 | -31585 | evm.model.Herato1202.447 | BMP and activin membrane-bound inhibitor homolog |
| pan | 370838910 | 370839181 | -7284 | evm.model.Herato1202.464 | dynactin subunit 5 |
| pan | 370838910 | 370839181 | -7284 | evm.model.Herato1202.464 | dynactin subunit 5 |
| pan | 375245994 | 375246291 | -14373 | evm.model.Herato1202.553 | LDLR chaperone boca |
| pan | 375742104 | 375742627 | 18221 | evm.model.Herato1202.573 | exonuclease GOR-like isoform X1 |
| pan | 375742104 | 375742627 | 18221 | evm.model.Herato1202.573 | exonuclease GOR-like isoform X1 |
| pan | 376778549 | 376778911 | 525 | evm.model.Herato1202.592 | derlin-2 |
| pan | 376812446 | 376812946 | -2223 | evm.model.Herato1202.596 | clustered mitochondria protein homolog isoform X1 |
| pan | 376978358 | 376979015 | -2724 | evm.model.Herato1202.600 | DNA-directed RNA polymerase II 16 kDa polypeptide |
| pan | 376978358 | 376979015 | -2724 | evm.model.Herato1202.600 | DNA-directed RNA polymerase II 16 kDa polypeptide |
| pan | 377926461 | 377927041 | 9602 | evm.model.Herato1202.621 | EF-hand domain-containing family member C2-like |
| pan | 377926461 | 377927041 | 9602 | evm.model.Herato1202.621 | EF-hand domain-containing family member C2-like |
| pan | 378528461 | 378528799 | -46737 | evm.model.Herato1202.659 | ---NA--- |
| pan | 378528461 | 378528799 | -46737 | evm.model.Herato1202.659 | ---NA--- |
| pan | 379264227 | 379264875 | 4704 | evm.model.Herato1202.684 | uncharacterized protein LOC117988862 |
| pan | 380074366 | 380074635 | -128090 | evm.model.Herato1202.712 | ---NA--- |
| pan | 380074366 | 380074635 | -128090 | evm.model.Herato1202.712 | ---NA--- |
| pan | 380660179 | 380660621 | 15692 | evm.model.Herato1202.729 | hemicentin-1 isoform X1 |
| pan | 381236152 | 381236647 | 6528 | evm.model.Herato1202.738 | centrosomal protein of 131 kDa |
| pan | 381236152 | 381236647 | 6528 | evm.model.Herato1202.738 | centrosomal protein of 131 kDa |
| pan | 383640261 | 383640727 | -11738 | evm.model.Herato1202.790 | amyloid protein-binding protein 2 |
| pan | 383640261 | 383640727 | -11738 | evm.model.Herato1202.790 | amyloid protein-binding protein 2 |
| pan | 385199154 | 385199416 | 121127 | evm.model.Herato1202.824 | glucose dehydrogenase [FAD, quinone]-like |
| pan | 387450620 | 387451458 | -139649 | evm.model.Herato1301.28 | neuropeptide FF receptor 1-like isoform X1 |
| pan | 389212757 | 389213079 | -219 | evm.model.Herato1301.52 | uncharacterized protein LOC113396369 |
| pan | 389212757 | 389213079 | -219 | evm.model.Herato1301.52 | uncharacterized protein LOC113396369 |
| pan | 389214617 | 389214935 | -2077 | evm.model.Herato1301.52 | uncharacterized protein LOC113396369 |
| pan | 389214617 | 389214935 | -2077 | evm.model.Herato1301.52 | uncharacterized protein LOC113396369 |
| pan | 389776242 | 389776690 | -7570 | evm.model.Herato1301.57 | elongation of very long chain fatty acids protein 6 |
| pan | 390301207 | 390301797 | -100259 | evm.model.Herato1301.64 | serine/threonine-protein kinase DDB_G0283821 |
| pan | 390733110 | 390733478 | -36338 | evm.model.Herato1301.72 | uncharacterized protein LOC116765631 |
| pan | 390733115 | 390733305 | -36422 | evm.model.Herato1301.72 | uncharacterized protein LOC116765631 |
| pan | 391823064 | 391823313 | 19983 | evm.model.Herato1301.97 | hypothetical protein KGM_210035 |
| pan | 393704447 | 393705372 | -476 | evm.model.Herato1301.126 | ---NA--- |
| pan | 396654243 | 396654479 | -4866 | evm.model.Herato1301.221 | mucin-5AC-like |
| pan | 403745976 | 403746473 | 2156 | evm.model.Herato1301.423 | probable RNA-binding protein 46 |
| pan | 406392660 | 406393288 | -21349 | evm.model.Herato1301.474 | ADP-ribosylation factor-like protein 8 |
| pan | 406407552 | 406408035 | -6530 | evm.model.Herato1301.474 | ADP-ribosylation factor-like protein 8 |
| pan | 406917596 | 406918131 | -21189 | evm.model.Herato1301.483 | elongation of very long chain fatty acids protein 7-like |
| pan | 409691497 | 409692357 | 99232 | evm.model.Herato1301.535 | pseudouridylate synthase 7 homolog |
| pan | 409691497 | 409692357 | 99232 | evm.model.Herato1301.535 | pseudouridylate synthase 7 homolog |

| | | | | | |
|---|---|---|---|---|---|
| pan | 411290118 | 411290869 | 49174 | evm.model.Herato1301.557 | neurobeachin isoform X1 |
| pan | 414459686 | 414460168 | 5752 | evm.model.Herato1301.596 | adiponectin receptor protein |
| pan | 414704100 | 414704283 | 10619 | evm.model.Herato1301.605 | hypothetical protein KGM_205988 |
| pan | 419975623 | 419975916 | 2198 | evm.model.Herato1301.692 | protein mini spindles isoform X1 |
| pan | 421207998 | 421208431 | 679 | evm.model.Herato1301.719 | uncharacterized protein LOC113396606 isoform X3 |
| pan | 421281863 | 421282086 | -16177 | evm.model.Herato1301.721 | ---NA--- |
| pan | 421281863 | 421282086 | -16177 | evm.model.Herato1301.721 | ---NA--- |
| pan | 421291478 | 421291686 | -6569 | evm.model.Herato1301.721 | ---NA--- |
| pan | 421380814 | 421381243 | -23474 | evm.model.Herato1301.722 | cyclin-dependent kinase inhibitor 1-like |
| pan | 421380814 | 421381243 | -23474 | evm.model.Herato1301.722 | cyclin-dependent kinase inhibitor 1-like |
| pan | 425163226 | 425163623 | -155 | evm.model.Herato1301.780 | protein cramped |
| pan | 427552088 | 427552707 | -39428 | evm.model.Herato1408.13 | E3 ubiquitin-protein ligase AMFR-like |
| pan | 427552088 | 427552707 | -39428 | evm.model.Herato1408.13 | E3 ubiquitin-protein ligase AMFR-like |
| pan | 427832052 | 427832531 | 30888 | evm.model.Herato1408.23 | cholinesterase 2-like |

| | | | | | |
|---|---|---|---|---|---|
| pan | 428197786 | 428198088 | -235217 | evm.model.Herato1408.26 | transcription factor Sox-12-like |
| pan | 428197786 | 428198088 | -235217 | evm.model.Herato1408.26 | transcription factor Sox-12-like |
| pan | 428203199 | 428203838 | -229636 | evm.model.Herato1408.26 | transcription factor Sox-12-like |
| pan | 428203199 | 428203838 | -229636 | evm.model.Herato1408.26 | transcription factor Sox-12-like |
| pan | 429521323 | 429521482 | 6034 | evm.model.Herato1408.31 | serine/threonine-protein kinase MARK2-like |
| pan | 430427849 | 430428337 | -41974 | evm.model.Herato1408.55 | oxysterol-binding protein-related protein 9 |
| pan | 430427849 | 430428337 | -41974 | evm.model.Herato1408.55 | oxysterol-binding protein-related protein 9 |
| pan | 432667287 | 432667616 | -6288 | evm.model.Herato1408.85 | probable beta-hexosaminidase fdl |
| pan | 432667287 | 432667616 | -6288 | evm.model.Herato1408.85 | probable beta-hexosaminidase fdl |
| pan | 437970063 | 437970732 | -24119 | evm.model.Herato1408.228 | glutamyl-tRNA(Gln) amidotransferase subunit B, mitochondrial |
| pan | 437970063 | 437970732 | -24119 | evm.model.Herato1408.228 | glutamyl-tRNA(Gln) amidotransferase subunit B, mitochondrial |
| pan | 438980051 | 438980780 | 52752 | novel_exon_163 | uncharacterized protein LOC113403206 |
| pan | 438980094 | 438980444 | 52606 | novel_exon_163 | uncharacterized protein LOC113403206 |
| pan | 440454848 | 440455266 | 10914 | evm.model.Herato1411.14 | RING finger protein nhl-1-like |
| pan | 442858726 | 442859102 | -8754 | evm.model.Herato1411.56 | potassium/sodium hyperpolarization-activated cyclic nucleotide-gated channel 2 isoform X1 |
| pan | 442858726 | 442859102 | -8754 | evm.model.Herato1411.56 | potassium/sodium hyperpolarization-activated cyclic nucleotide-gated channel 2 isoform X1 |
| pan | 445478469 | 445478653 | 18302 | evm.model.Herato1411.123 | protein FAM91A1 |
| pan | 447412741 | 447413157 | 3589 | evm.model.Herato1411.170 | hemocyte protein-glutamine gammaglutamyltransferase-like |
| pan | 447919995 | 447920711 | 196 | evm.model.Herato1411.178 | phospholipid phosphatase 2-like isoform X1 |
| pan | 450218566 | 450218888 | 15645 | evm.model.Herato1411.196 | pregnancy zone protein-like |
| pan | 450273835 | 450274316 | 10368 | evm.model.Herato1411.199 | pregnancy zone protein-like |
| pan | 450273835 | 450274316 | 10368 | evm.model.Herato1411.199 | pregnancy zone protein-like |
| pan | 450885742 | 450886049 | 46707 | evm.model.Herato1411.214_evm.T | nose resistant to fluoxetine protein 6-like |
| pan | 451735938 | 451736330 | -20544 | evm.model.Herato1411.231 | endonuclease-reverse transcriptase |
| pan | 452528225 | 452528587 | -12243 | evm.model.Herato1411.254 | solute carrier family 22 member 1-like |
| pan | 452528225 | 452528587 | -12243 | evm.model.Herato1411.254 | solute carrier family 22 member 1-like |
| pan | 458299071 | 458299762 | 208876 | evm.model.Herato1505.86 | poly(A)-specific ribonuclease PARN-like isoform X1 |
| pan | 458299071 | 458299762 | 208876 | evm.model.Herato1505.86 | poly(A)-specific ribonuclease PARN-like isoform X1 |
| pan | 458487032 | 458487490 | 21031 | evm.model.Herato1505.86 | poly(A)-specific ribonuclease PARN-like isoform X1 |
| pan | 458891679 | 458891965 | -11 | evm.model.Herato1505.100 | protein penguin |
| pan | 458891679 | 458891965 | -11 | evm.model.Herato1505.100 | protein penguin |
| pan | 459754069 | 459754512 | 19245 | evm.model.Herato1505.131 | RNA-directed DNA polymerase from mobile element jockey |
| pan | 459921640 | 459921997 | -18023 | evm.model.Herato1505.133 | protein FAM234A |
| pan | 465267855 | 465268537 | 11691 | evm.model.Herato1507.46 | Protein pangolin, isoforms A/H/I |
| pan | 469425289 | 469425695 | -11630 | evm.model.Herato1507.135 | alpha-(1,3)-fucosyltransferase 6-like isoform X1 |
| pan | 472021611 | 472022075 | -5632 | evm.model.Herato1507.159 | uncharacterized protein LOC110377276 |

| | | | | | |
|---|---|---|---|---|---|
| pan | 473217468 | 473217878 | 17394 | evm.model.Herato1507.187 | uncharacterized protein LOC113394940 |
| pan | 473217468 | 473217878 | 17394 | evm.model.Herato1507.187 | uncharacterized protein LOC113394940 |
| pan | 477589041 | 477589375 | -64886 | evm.model.Herato1507.262 | uncharacterized protein LOC113402867 |
| pan | 477589041 | 477589375 | -64886 | evm.model.Herato1507.262 | uncharacterized protein LOC113402867 |
| pan | 478054059 | 478054580 | -3995 | evm.model.Herato1507.272 | E3 ubiquitin-protein ligase RING1 |
| pan | 478097199 | 478097423 | 31655 | evm.model.Herato1507.273 | 2-hydroxyacyl-CoA lyase 1 isoform X1 |
| pan | 478870954 | 478871677 | 1600 | evm.model.Herato1507.291 | vacuolar protein sorting-associated protein 18 homolog |
| pan | 487972341 | 487972502 | 11649 | evm.model.Herato1524.19 | ---NA--- |
| pan | 488854846 | 488855309 | -20090 | evm.model.Herato1601.22 | general transcriptional corepressor trfA isoform X1 |
| pan | 488854846 | 488855309 | -20090 | evm.model.Herato1601.22 | general transcriptional corepressor trfA isoform X1 |
| pan | 488861847 | 488862211 | -27042 | evm.model.Herato1601.22 | general transcriptional corepressor trfA isoform X1 |
| pan | 500084469 | 500084942 | -101664 | evm.model.Herato1602.9 | G-protein coupled receptor moody-like |
| pan | 501436919 | 501437205 | 1591 | evm.model.Herato1603.20 | xaa-Pro dipeptidase |
| pan | 507198770 | 507199663 | 2421 | evm.model.Herato1605.14 | diphthine--ammonia ligase |
| pan | 508789757 | 508790200 | -18545 | evm.model.Herato1605.35 | inositol-trisphosphate 3-kinase B isoform X3 |
| pan | 508884965 | 508885189 | 7494 | evm.model.Herato1605.36 | inositol-trisphosphate 3-kinase A isoform X1 |
| pan | 509170978 | 509171664 | 3344 | evm.model.Herato1605.40 | unnamed protein product |
| pan | 509171238 | 509171716 | 3188 | evm.model.Herato1605.40 | unnamed protein product |
| pan | 509184961 | 509185214 | -10422 | evm.model.Herato1605.40 | unnamed protein product |
| pan | 510574594 | 510574851 | 6412 | evm.model.Herato1605.174 | uncharacterized protein LOC110991711 |
| pan | 510665326 | 510665878 | 38760 | evm.model.Herato1605.65 | disheveled-associated activator of morphogenesis 1 isoform X1 |
| pan | 510665326 | 510665878 | 38760 | evm.model.Herato1605.65 | disheveled-associated activator of morphogenesis 1 isoform X1 |
| pan | 511337013 | 511337275 | 5395 | evm.model.Herato1605.84 | proline-rich protein 12-like |
| pan | 511375270 | 511375514 | -32853 | evm.model.Herato1605.84 | proline-rich protein 12-like |
| pan | 511389567 | 511389920 | -47204 | evm.model.Herato1605.84 | proline-rich protein 12-like |
| pan | 515234365 | 515234843 | -13216 | evm.model.Herato1605.131 | uncharacterized protein LOC113402104 isoform X1 |

| | | | | | |
|---|---|---|---|---|---|
| pan | 515560289 | 515561008 | 9927 | evm.model.Herato1605.141 | symplekin |
| pan | 515560350 | 515561120 | 9840 | evm.model.Herato1605.141 | symplekin |
| pan | 516844037 | 516844791 | 36528 | evm.model.Herato1701.18 | spondin-1 |
| pan | 516844037 | 516844791 | 36528 | evm.model.Herato1701.18 | spondin-1 |
| pan | 518247365 | 518247822 | -339539 | evm.model.Herato1701.36 | teneurin-m isoform X1 |
| pan | 518247365 | 518247822 | -339539 | evm.model.Herato1701.36 | teneurin-m isoform X1 |
| pan | 518431725 | 518432094 | -155223 | evm.model.Herato1701.36 | teneurin-m isoform X1 |
| pan | 518431725 | 518432094 | -155223 | evm.model.Herato1701.36 | teneurin-m isoform X1 |
| pan | 518547758 | 518548748 | -38879 | evm.model.Herato1701.36 | teneurin-m isoform X1 |
| pan | 518547758 | 518548748 | -38879 | evm.model.Herato1701.36 | teneurin-m isoform X1 |
| pan | 525418840 | 525419228 | -29201 | evm.model.Herato1701.114 | flocculation protein FLO11 isoform X1 |
| pan | 525594331 | 525594553 | 8479 | evm.model.Herato1701.125 | MIF-like protein mif-2 |
| pan | 525594331 | 525594553 | 8479 | evm.model.Herato1701.125 | MIF-like protein mif-2 |
| pan | 527741856 | 527742591 | 1864 | evm.model.Herato1701.180 | TIP41-like protein |
| pan | 527741856 | 527742591 | 1864 | evm.model.Herato1701.180 | TIP41-like protein |
| pan | 527841872 | 527842169 | -18871 | evm.model.Herato1701.183 | gamma-tubulin complex component 2-like isoform X2 |
| pan | 527854336 | 527854836 | 22789 | evm.model.Herato1701.184 | uncharacterized protein LOC110997031 |
| pan | 527854336 | 527854836 | 22789 | evm.model.Herato1701.184 | uncharacterized protein LOC110997031 |
| pan | 527949894 | 527950258 | -3359 | evm.model.Herato1701.186 | single-stranded DNA-binding protein, mitochondrial |
| pan | 530287860 | 530288066 | 2417 | evm.model.Herato1701.227 | Macrophage MHC class 1 receptor 2 protein |
| pan | 530287860 | 530288066 | 2417 | evm.model.Herato1701.227 | Macrophage MHC class 1 receptor 2 protein |
| pan | 530833825 | 530834056 | -3700 | evm.model.Herato1701.248 | LIM domain only protein 3-like |
| pan | 530833825 | 530834056 | -3700 | evm.model.Herato1701.248 | LIM domain only protein 3-like |
| pan | 532994407 | 532994666 | -18678 | evm.model.Herato1701.307 | cytoplasmic dynein 2 heavy chain 1 |
| pan | 537166850 | 537167407 | 123999 | evm.model.Herato1703.2 | tyrosine-protein phosphatase Lar-like isoform X1 |
| pan | 538095883 | 538096547 | -157 | evm.model.Herato1703.13 | CREB-binding protein-like |

| | | | | | |
|---|---|---|---|---|---|
| pan | 538095883 | 538096547 | -157 | evm.model.Herato1703.13 | CREB-binding protein-like |
| pan | 538536221 | 538536704 | 14549 | evm.model.Herato1703.47 | ---NA--- |
| pan | 538536221 | 538536704 | 14549 | evm.model.Herato1703.47 | ---NA--- |
| pan | 544304925 | 544305237 | 111 | evm.model.Herato1705.19 | leucine-rich repeat serine/threonine-protein kinase 1 |
| pan | 544304925 | 544305237 | 111 | evm.model.Herato1705.19 | leucine-rich repeat serine/threonine-protein kinase 1 |
| pan | 546462952 | 546463359 | 164 | evm.model.Herato1705.97 | DNA-directed RNA polymerases I, II, and III subunit RPABC3 |
| pan | 547697556 | 547697776 | -15301 | evm.model.Herato1708.12 | dynactin subunit 1 |
| pan | 551023576 | 551024217 | -5811 | evm.model.Herato1708.126 | phenoloxidase-activating factor 2-like |
| pan | 558156738 | 558157040 | -5550 | evm.model.Herato1801.128 | delta(14)-sterol reductase |
| pan | 560148271 | 560149088 | 13128 | evm.model.Herato1801.164 | ---NA--- |
| pan | 567622489 | 567623089 | -69501 | evm.model.Herato1805.88 | fatty acid synthase-like |
| pan | 567622489 | 567623089 | -69501 | evm.model.Herato1805.88 | fatty acid synthase-like |
| pan | 568539571 | 568539841 | -36359 | evm.model.Herato1805.102 | circadian clock-controlled protein-like |
| pan | 571539788 | 571540229 | -415 | evm.model.Herato1805.135 | axoneme-associated protein mst101(2) isoform X1 |
| pan | 571539788 | 571540229 | -415 | evm.model.Herato1805.135 | axoneme-associated protein mst101(2) isoform X1 |
| pan | 575223207 | 575223770 | -20368 | evm.model.Herato1805.220 | organic cation transporter protein-like |
| pan | 575681339 | 575681641 | -111627 | evm.model.Herato1805.227 | Down syndrome cell adhesion molecule-like protein 1 homolog |
| pan | 575681424 | 575681640 | -111585 | evm.model.Herato1805.227 | Down syndrome cell adhesion molecule-like protein 1 homolog |
| pan | 577125471 | 577126205 | -20644 | evm.model.Herato1805.249 | glycine receptor subunit alpha-2-like |
| pan | 577125471 | 577126205 | -20644 | evm.model.Herato1805.249 | glycine receptor subunit alpha-2-like |
| pan | 579145988 | 579146389 | -56668 | evm.model.Herato1805.290 | seminal fluid protein HACP027 |
| pan | 580467057 | 580467787 | 12657 | evm.model.Herato1807.5 | Tyrosine-protein phosphatase 99A |
| pan | 580467057 | 580467787 | 12657 | evm.model.Herato1807.5 | Tyrosine-protein phosphatase 99A |
| pan | 586031169 | 586031426 | 7785 | evm.model.Herato1807.83 | ---NA--- |
| pan | 586031169 | 586031426 | 7785 | evm.model.Herato1807.83 | ---NA--- |
| pan | 589246057 | 589246580 | -20752 | evm.model.Herato1901.17 | putative lymphocyte cytosolic protein 2 |
| pan | 589246057 | 589246580 | -20752 | evm.model.Herato1901.17 | putative lymphocyte cytosolic protein 2 |
| pan | 589249593 | 589250096 | -17226 | evm.model.Herato1901.17 | putative lymphocyte cytosolic protein 2 |
| pan | 593862082 | 593862442 | 15325 | evm.model.Herato1901.91 | uncharacterized protein LOC113398113 |
| pan | 593862082 | 593862442 | 15325 | evm.model.Herato1901.91 | uncharacterized protein LOC113398113 |
| pan | 595650757 | 595651253 | -164 | evm.model.Herato1903.5 | serine/threonine-protein kinase pelle |
| pan | 598142735 | 598142935 | -36632 | evm.model.Herato1904.42 | vesicular integral-membrane protein VIP36 |
| pan | 598943338 | 598944004 | -42940 | evm.model.Herato1904.61 | disintegrin and metalloproteinase domain-containing protein 12 |
| pan | 602753516 | 602754094 | -4034 | evm.model.Herato1904.139 | ABC transporter G family member 23 isoform X1 |
| pan | 602753516 | 602754094 | -4034 | evm.model.Herato1904.139 | ABC transporter G family member 23 isoform X1 |
| pan | 603259012 | 603259289 | 2920 | evm.model.Herato1904.147 | TBC1 domain family member 7 |
| pan | 605437457 | 605438070 | 268 | evm.model.Herato1904.204 | UBX domain-containing protein 1 |
| pan | 605437457 | 605438070 | 268 | evm.model.Herato1904.204 | UBX domain-containing protein 1 |
| pan | 606002476 | 606002710 | -7735 | evm.model.Herato1904.213 | excitatory amino acid transporter isoform X2 |
| pan | 607259025 | 607259225 | 75318 | evm.model.Herato1904.238 | hypothetical protein EVAR_56129_1 |
| pan | 618594846 | 618595131 | -11732 | evm.model.Herato1908.103 | probable salivary secreted peptide |
| pan | 618961792 | 618962203 | 1557 | evm.model.Herato1910.19 | RNA polymerase II transcriptional coactivator |
| pan | 618961792 | 618962203 | 1557 | evm.model.Herato1910.19 | RNA polymerase II transcriptional coactivator |
| pan | 620462672 | 620463384 | -21850 | evm.model.Herato1910.69 | multidrug resistance-associated protein 1 isoform X2 |
| pan | 620462672 | 620463384 | -21850 | evm.model.Herato1910.69 | multidrug resistance-associated protein 1 isoform X2 |
| pan | 620885850 | 620886111 | -2536 | evm.model.Herato1910.82 | phospholipase A1-like |
| pan | 622199723 | 622200127 | 50688 | evm.model.Herato1910.116 | Beat-IIIc isoform A |
| pan | 622199723 | 622200127 | 50688 | evm.model.Herato1910.116 | Beat-IIIc isoform A |
| pan | 625362844 | 625363696 | 155285 | evm.model.Herato1910.159 | kinesin-like protein CG14535 isoform X2 |
| pan | 625938802 | 625939115 | 5176 | evm.model.Herato1910.175 | ---NA--- |
| pan | 627420771 | 627421019 | -45418 | evm.model.Herato1910.187 | Leucine-rich repeats and immunoglobulin-like domains protein 2 |

| | | | | | |
|---|---|---|---|---|---|
| pan | 627420771 | 627421019 | -45418 | evm.model.Herato1910.187 | Leucine-rich repeats and immunoglobulin-like domains protein 2 |
| pan | 629945728 | 629946142 | 34517 | evm.model.Herato2001.14 | hyphal wall protein 2-like |
| pan | 630300769 | 630301023 | -205483 | evm.model.Herato2001.17 | zinc finger protein Xfin-like |
| pan | 630300769 | 630301023 | -205483 | evm.model.Herato2001.17 | zinc finger protein Xfin-like |
| pan | 631425366 | 631425568 | 35655 | evm.model.Herato2001.29 | protein KIAA0100 |
| pan | 632079178 | 632079801 | 23614 | evm.model.Herato2001.43 | chitin binding protein |
| pan | 632079178 | 632079801 | 23614 | evm.model.Herato2001.43 | chitin binding protein |
| pan | 635559516 | 635560032 | 103608 | evm.model.Herato2001.133 | troponin C-like |
| pan | 639973245 | 639973529 | 3959 | evm.model.Herato2001.242 | NADPH--cytochrome P450 reductase isoform X1 |
| pan | 639973245 | 639973529 | 3959 | evm.model.Herato2001.242 | NADPH--cytochrome P450 reductase isoform X1 |
| pan | 641532822 | 641533087 | -8697 | evm.model.Herato2001.314 | RNA binding protein fox-1 homolog 3 |
| pan | 647428703 | 647429004 | 27704 | evm.model.Herato2001.443 | protein white |
| pan | 647428703 | 647429004 | 27704 | evm.model.Herato2001.443 | protein white |
| pan | 649285332 | 649285635 | 1663 | evm.model.Herato2001.501 | fidgetin-like protein 1 |
| pan | 652480065 | 652480387 | 9215 | evm.model.Herato2001.584 | kinesin-related protein 6-like isoform X1 |
| pan | 652554277 | 652554896 | 4059 | evm.model.Herato2001.585 | fatty acyl-CoA reductase wat-like |
| pan | 652610607 | 652611263 | 31734 | evm.model.Herato2001.586 | zinc transporter ZIP9 |
| pan | 653604348 | 653604624 | -66828 | evm.model.Herato2001.607 | kanadaptin |
| pan | 655510817 | 655511201 | -167046 | evm.model.Herato2001.648 | synaptotagmin-7 isoform X2 |
| pan | 658089326 | 658089656 | -1856 | evm.model.Herato2001.699 | acetyl-CoA carboxylase isoform X2 |
| pan | 658833017 | 658833327 | 7120 | evm.model.Herato2001.718 | ankyrin-3-like isoform X2 |
| pan | 659072893 | 659073379 | -3235 | evm.model.Herato2001.722 | suppression of tumorigenicity 5 protein-like |
| pan | 660402353 | 660402680 | -120 | evm.model.Herato2101.7 | autophagy-related protein 13 homolog |
| pan | 660519052 | 660519575 | 3890 | evm.model.Herato2101.11 | transmembrane protein 198 |
| pan | 660928477 | 660928908 | -981 | evm.model.Herato2101.20 | L-galactose dehydrogenase-like |
| pan | 661712380 | 661712994 | -59944 | evm.model.Herato2101.34 | hypothetical protein KGM_202987 |
| pan | 662595686 | 662595921 | -81094 | evm.model.Herato2101.49 | protein cycle isoform X3 |
| pan | 664962743 | 664963134 | -30606 | evm.model.Herato2101.110 | probable cytochrome P450 303a1 |
| pan | 664962753 | 664963124 | -30606 | evm.model.Herato2101.110 | probable cytochrome P450 303a1 |
| pan | 664972771 | 664973430 | -40768 | evm.model.Herato2101.110 | probable cytochrome P450 303a1 |
| pan | 664972771 | 664973448 | -40777 | evm.model.Herato2101.110 | probable cytochrome P450 303a1 |
| pan | 665239539 | 665240287 | 23014 | evm.model.Herato2101.112 | dual specificity mitogen-activated protein kinase kinase 1-like |
| pan | 665752529 | 665752940 | 130086 | evm.model.Herato2101.113 | dipeptidase 1-like isoform X2 |
| pan | 668144962 | 668145321 | -9271 | evm.model.Herato2101.160 | Intraflagellar transport protein 122 |
| pan | 669605099 | 669605698 | 20613 | evm.model.Herato2101.201 | solute carrier family 15 member 1-like |
| pan | 670536577 | 670537213 | 25321 | evm.model.Herato2101.220 | prostatic acid phosphatase-like |
| pan | 670536646 | 670537185 | 25301 | evm.model.Herato2101.220 | prostatic acid phosphatase-like |
| pan | 670734642 | 670735150 | -4840 | evm.model.Herato2101.227 | lysophospholipid acyltransferase 5 |
| pan | 670777951 | 670778391 | -180 | evm.model.Herato2101.229 | protein enabled homolog |
| pan | 670972036 | 670972218 | 5860 | evm.model.Herato2101.236 | uncharacterized protein LOC113404094 |
| pan | 671096433 | 671097037 | 2111 | evm.model.Herato2101.242 | uncharacterized protein LOC113404092 |
| pan | 671796507 | 671796679 | -3582 | evm.model.Herato2101.261 | Neuronal pentraxin-2 |
| pan | 672499738 | 672500063 | -9645 | evm.model.Herato2101.272 | DNA-binding protein D-ETS-3 isoform X1 |
| pan | 672831908 | 672832094 | 11168 | evm.model.Herato2101.280 | LIM/homeobox protein Lhx9-like |
| pan | 673076645 | 673076869 | 15569 | evm.model.Herato2101.283 | probable NADH dehydrogenase [ubiquinone] 1 alpha subcomplex subunit 12 |
| pan | 674595737 | 674595956 | -66374 | evm.model.Herato2101.309 | ---NA--- |
| pan | 676150557 | 676151098 | -517 | evm.model.Herato2101.342 | peptidyl-prolyl cis-trans isomerase-like 1 |
| pan | 676150643 | 676151098 | -474 | evm.model.Herato2101.342 | peptidyl-prolyl cis-trans isomerase-like 1 |
| pan | 679607648 | 679608279 | 31693 | evm.model.Herato2101.402 | Nucleic-acid-binding protein from transposon Xelement |
| pan | 684968853 | 684969231 | -17192 | evm.model.Herato2101.489 | mediator of RNA polymerase II transcription subunit 23 |
| pan | 687219693 | 687219983 | -17529 | evm.model.Herato2101.513 | ---NA--- |

notabilis-etylus hybridzone

| Chr | Start | End | TSS | gene ID | gene name |
|---|---|---|---|---|---|
| pan | 1605135 | 1605616 | -90218 | evm.model.Herato0101.19 | nephrin |
| pan | 1605135 | 1605616 | -90218 | evm.model.Herato0101.19 | nephrin |

| | | | | | |
|---|---|---|---|---|---|
| pan | 6276632 | 6276993 | -61843 | evm.model.Herato0101.116 | hypothetical protein EVAR_74554_1 |
| pan | 6276632 | 6276993 | -61843 | evm.model.Herato0101.116 | hypothetical protein EVAR_74554_1 |
| pan | 13657376 | 13657890 | -615 | evm.model.Herato0101.234 | signal transducing adapter molecule 1 |
| pan | 13657376 | 13657890 | -615 | evm.model.Herato0101.234 | signal transducing adapter molecule 1 |
| pan | 14129526 | 14129780 | -53734 | evm.model.Herato0101.241 | hypothetical protein EVAR_80933_1 |
| pan | 17792572 | 17792960 | 40483 | evm.model.Herato0101.296 | protein spire isoform X4 |
| pan | 23940215 | 23940540 | 35192 | evm.model.Herato0101.505 | Zinc finger CCHC domain-containing protein 2 |
| pan | 23940215 | 23940540 | 35192 | evm.model.Herato0101.505 | Zinc finger CCHC domain-containing protein 2 |
| pan | 26332556 | 26332910 | -4477 | evm.model.Herato0101.568 | ---NA--- |
| pan | 26332556 | 26332910 | -4477 | evm.model.Herato0101.568 | ---NA--- |
| pan | 35110929 | 35111103 | -26360 | evm.model.Herato0101.800 | dedicator of cytokinesis protein 7 isoform X2 |
| pan | 35110929 | 35111103 | -26360 | evm.model.Herato0101.800 | dedicator of cytokinesis protein 7 isoform X2 |
| pan | 35995814 | 35996042 | -8956 | evm.model.Herato0101.826 | glutenin, high molecular weight subunit DX5-like |
| pan | 35995814 | 35996042 | -8956 | evm.model.Herato0101.826 | glutenin, high molecular weight subunit DX5-like |
| pan | 36982574 | 36982853 | 16807 | evm.model.Herato0101.847 | carbonyl reductase [NADPH] 3-like |
| pan | 40385859 | 40386148 | -276906 | evm.model.Herato0204.28 | ---NA--- |
| pan | 40385859 | 40386148 | -276906 | evm.model.Herato0204.28 | ---NA--- |
| pan | 47403311 | 47403589 | -27716 | evm.model.Herato0209.47 | uncharacterized protein LOC113395221 |
| pan | 48098230 | 48098500 | 51952 | evm.model.Herato0209.67 | flotillin-2 isoform X1 |
| pan | 48098230 | 48098500 | 51952 | evm.model.Herato0209.67 | flotillin-2 isoform X1 |
| pan | 55888488 | 55889005 | -7048 | evm.model.Herato0214.15 | protein unc-119 homolog |
| pan | 56240827 | 56241073 | 7434 | evm.model.Herato0214.21 | RILP protein |
| pan | 57467577 | 57467997 | 14045 | evm.model.Herato0214.54 | luciferin 4-monooxygenase-like |
| pan | 57467577 | 57467997 | 14045 | evm.model.Herato0214.54 | luciferin 4-monooxygenase-like |
| pan | 58478191 | 58478746 | -13038 | evm.model.Herato0215.147 | hypothetical protein |
| pan | 58478206 | 58478760 | -13023 | evm.model.Herato0215.147 | hypothetical protein |
| pan | 59014803 | 59015305 | 104210 | evm.model.Herato0215.16 | fructose-bisphosphate aldolase isoform X1 |
| pan | 59014803 | 59015305 | 104210 | evm.model.Herato0215.16 | fructose-bisphosphate aldolase isoform X1 |
| pan | 65154561 | 65155053 | 43188 | evm.model.Herato0215.143 | F-box/LRR-repeat protein 7-like |
| pan | 65310311 | 65310658 | 11605 | evm.model.Herato0215.145 | apolipophorin-3-like |
| pan | 65310311 | 65310658 | 11605 | evm.model.Herato0215.145 | apolipophorin-3-like |
| pan | 78052216 | 78053201 | 8166 | evm.model.Herato0310.17 | NADP-dependent malic enzyme |
| pan | 78052216 | 78053201 | 8166 | evm.model.Herato0310.17 | NADP-dependent malic enzyme |
| pan | 78053606 | 78054030 | 7056 | evm.model.Herato0310.17 | NADP-dependent malic enzyme |
| pan | 78053606 | 78054030 | 7056 | evm.model.Herato0310.17 | NADP-dependent malic enzyme |
| pan | 83389895 | 83390116 | 2077 | evm.model.Herato0310.77 | microtubule-associated protein tau isoform X3 |
| pan | 83389895 | 83390116 | 2077 | evm.model.Herato0310.77 | microtubule-associated protein tau isoform X3 |
| pan | 87334574 | 87335509 | 8672 | evm.model.Herato0310.146 | testin |
| pan | 87334574 | 87335509 | 8672 | evm.model.Herato0310.146 | testin |
| pan | 90473573 | 90473833 | 40962 | evm.model.Herato0310.192 | translation initiation factor eIF-2B subunit alpha |
| pan | 90473573 | 90473833 | 40962 | evm.model.Herato0310.192 | translation initiation factor eIF-2B subunit alpha |
| pan | 94688111 | 94688582 | -30732 | evm.model.Herato0310.299 | transmembrane protease serine 12-like |
| pan | 96538512 | 96538733 | 56841 | evm.model.Herato0401.5 | zwei Ig domain protein zig-8-like |
| pan | 96538512 | 96538733 | 56841 | evm.model.Herato0401.5 | zwei Ig domain protein zig-8-like |
| pan | 96564053 | 96564533 | 82512 | evm.model.Herato0401.5 | zwei Ig domain protein zig-8-like |
| pan | 98043098 | 98043785 | -11104 | evm.model.Herato0401.15 | chondroitin sulfate synthase 1 |
| pan | 100338531 | 100338944 | -7041 | evm.model.Herato0401.55 | serine protease snake-like |
| pan | 100338531 | 100338944 | -7041 | evm.model.Herato0401.55 | serine protease snake-like |
| pan | 101553242 | 101553573 | -54527 | evm.model.Herato0403.3 | protein groucho-like isoform X6 |
| pan | 102331096 | 102331428 | -37006 | evm.model.Herato0403.39 | ---NA--- |
| pan | 110368312 | 110368515 | 195837 | evm.model.Herato0411.29 | serine/arginine repetitive matrix protein 2 |
| pan pan | 110368312 | 110368515 | 195837 | evm.model.Herato0411.29 | serine/arginine repetitive matrix protein 2tumor necrosis |
| | 118094013 | 118094257 | 9176 | evm.model.Herato0411.120 | factor alpha-induced protein 8-like protein isoform X1 |
| pan | 118598012 | 118598805 | 69103 | evm.model.Herato0411.133 | TP53-binding protein 1-like |
| pan | 120909891 | 120910623 | -114765 | evm.model.Herato0412.1 | uncharacterized protein LOC113402078 isoform X2 |

| | | | | | |
|---|---|---|---|---|---|
| pan | 120926564 | 120926988 | -98246 | evm.model.Herato0412.1 | uncharacterized protein LOC113402078 isoform X2 |
| pan | 120926564 | 120926988 | -98246 | evm.model.Herato0412.1 | uncharacterized protein LOC113402078 isoform X2 |
| pan | 124902108 | 124903038 | -355 | evm.model.Herato0419.41 | phosphatidylinositol 3-kinase catalytic subunit type 3 |
| pan | 126373838 | 126374483 | 59172 | evm.model.Herato0419.66 | hydrocephalus-inducing protein homolog |
| pan | 126373838 | 126374483 | 59172 | evm.model.Herato0419.66 | hydrocephalus-inducing protein homolog |
| pan | 127698555 | 127699072 | 7195 | evm.model.Herato0501.13 | Bm8 interacting protein 2d-2 |
| pan | 127698555 | 127699072 | 7195 | evm.model.Herato0501.13 | Bm8 interacting protein 2d-2 |
| pan | 131754956 | 131755180 | -13443 | evm.model.Herato0503.30 | Outer dense fiber protein 3-like protein 2 |
| pan | 131754956 | 131755180 | -13443 | evm.model.Herato0503.30 | Outer dense fiber protein 3-like protein 2 |
| pan | 133158685 | 133159612 | -5378 | evm.model.Herato0503.64 | inhibitor of growth protein 5 |
| pan | 133158685 | 133159612 | -5378 | evm.model.Herato0503.64 | inhibitor of growth protein 5 |
| pan | 135554534 | 135555791 | 5356 | evm.model.Herato0503.112 | carboxylesterase 1E |
| pan | 135554543 | 135554870 | 4900 | evm.model.Herato0503.112 | carboxylesterase 1E |
| pan | 139534748 | 139534958 | -23210 | evm.model.Herato0503.143 | protein gooseberry-neuro isoform X1 |
| pan | 139534748 | 139534958 | -23210 | evm.model.Herato0503.143 | protein gooseberry-neuro isoform X1 |
| pan | 140277246 | 140277683 | -208 | evm.model.Herato0503.171 | ---NA--- |
| pan | 140578354 | 140579012 | -21291 | evm.model.Herato0503.180 | Transposable element Tcb1 transposase |
| pan | 140578354 | 140579012 | -21291 | evm.model.Herato0503.180 | Transposable element Tcb1 transposase |
| pan | 143226089 | 143226663 | 123 | evm.model.Herato0503.245 | 60S ribosomal protein L37a |
| pan | 149947160 | 149947380 | 52170 | evm.model.Herato0508.56 | N6-adenosine-methyltransferase non-catalytic subunit |
| pan | 149947160 | 149947380 | 52170 | evm.model.Herato0508.56 | N6-adenosine-methyltransferase non-catalytic subunit |
| pan | 154482099 | 154482305 | 8322 | evm.model.Herato0601.4 | tensin-2-like isoform X3 |
| pan | 154482099 | 154482305 | 8322 | evm.model.Herato0601.4 | tensin-2-like isoform X3 |
| pan | 155895566 | 155895808 | -70178 | evm.model.Herato0601.10 | zinc finger protein Noc |
| pan | 156326465 | 156326645 | -24580 | evm.model.Herato0601.12 | solute carrier family 22 member 13-like |
| pan | 156326465 | 156326645 | -24580 | evm.model.Herato0601.12 | solute carrier family 22 member 13-like |
| pan | 160037581 | 160037785 | 15470 | evm.model.Herato0601.77 | corticotropin-releasing factor-binding protein |
| pan | 161338029 | 161338362 | -74536 | evm.model.Herato0601.104 | protein timeless homolog |
| pan | 161338029 | 161338362 | -74536 | evm.model.Herato0601.104 | protein timeless homolog |
| pan | 166080075 | 166080333 | -349 | evm.model.Herato0606.53 | serine protease gd-like |
| pan | 166080075 | 166080333 | -349 | evm.model.Herato0606.53 | serine protease gd-like |
| pan | 168821698 | 168822130 | -12211 | evm.model.Herato0606.102 | uncharacterized protein LOC113401263 |
| pan | 170139915 | 170140100 | -50440 | evm.model.Herato0606.117 | Synaptotagmin-like protein 4 |
| pan | 170139915 | 170140100 | -50440 | evm.model.Herato0606.117 | Synaptotagmin-like protein 4 |
| pan | 172919591 | 172919764 | 17420 | evm.model.Herato0606.182 | Inactive ubiquitin carboxyl-terminal hydrolase 54 |
| pan | 172919591 | 172919764 | 17420 | evm.model.Herato0606.182 | Inactive ubiquitin carboxyl-terminal hydrolase 54 |
| pan | 176506950 | 176507231 | 19841 | evm.model.Herato0606.254 | uncharacterized protein LOC113398311 |
| pan | 176506950 | 176507231 | 19841 | evm.model.Herato0606.254 | uncharacterized protein LOC113398311 |
| pan | 179805225 | 179805689 | -2402 | evm.model.Herato0606.293 | palmitoyltransferase ZDHHC6 |
| pan | 179805225 | 179805689 | -2402 | evm.model.Herato0606.293 | palmitoyltransferase ZDHHC6 |
| pan | 179934886 | 179935619 | 6899 | evm.model.Herato0606.302 | neurexin-4 isoform X1 |
| pan | 184053908 | 184054163 | -4124 | evm.model.Herato0606.417 | L-dopachrome tautomerase yellow-f-like |
| pan pan pan | 185243322<br>191709203<br>191709203 | 185244139<br>191709400<br>191709400 | -18519<br>-19187<br>-19187 | evm.model.Herato0606.452<br>evm.model.Herato0701.47<br>evm.model.Herato0701.47 | uncharacterized protein LOC113391393pyruvate dehydrogenase (acetyl-transferring) kinase, mitochondrialpyruvate dehydrogenase (acetyl-transferring) kinase, mitochondrial |
| pan | 194092931 | 194093159 | -56988 | evm.model.Herato0701.148 | probable nuclear hormone receptor HR3 isoform X1 |
| pan | 194886840 | 194887224 | -22104 | evm.model.Herato0701.165 | cullin-1 |
| pan | 201316770 | 201317131 | 15461 | evm.model.Herato0701.307 | ---NA--- |
| pan | 201316770 | 201317131 | 15461 | evm.model.Herato0701.307 | ---NA--- |
| pan | 208671571 | 208671899 | 28845 | evm.model.Herato0701.469 | ethanolamine-phosphate cytidylyltransferase |

| | | | | |
|---|---|---|---|---|
| pan | 208671571 208671899 | 28845 | evm.model.Herato0701.469 | ethanolamine-phosphate cytidylyltransferase |
| pan | 212736149 212736409 | -140734 | evm.model.Herato0701.624 | collagen alpha-1(XVIII) chain-like |
| pan | 214475082 214475371 | 11752 | evm.model.Herato0701.643 | protein extra-macrochaetae |
| pan | 214475082 214475371 | 11752 | evm.model.Herato0701.643 | protein extra-macrochaetae |
| pan | 214564094 214564751 | -5967 | evm.model.Herato0701.645 | intersectin-1 isoform X2 |
| pan | 220003590 220003852 | -33835 | evm.model.Herato0701.733 | retinol dehydrogenase 11-like |
| pan | 220003590 220003852 | -33835 | evm.model.Herato0701.733 | retinol dehydrogenase 11-like |
| pan | 220165983 220166408 | -49650 | evm.model.Herato0701.735 | facilitated trehalose transporter Tret1 isoform X5 |
| pan | 223077329 223077543 | 94111 | evm.model.Herato0801.5 | ---NA--- |
| pan | 231332820 231333144 | 1533 | evm.model.Herato0801.94 | nucleolin 2-like |

| | | | | |
|---|---|---|---|---|
| pan | 231332820 231333144 | 1533 | evm.model.Herato0801.94 | nucleolin 2-like |
| pan | 231603369 231603614 | -55423 | evm.model.Herato0801.100 | putative epidermal cell surface receptor isoform X1 |
| pan | 232108618 232109042 | -103518 | evm.model.Herato0801.106 | ---NA--- |
| pan | 232108685 232109048 | -103482 | evm.model.Herato0801.106 | ---NA--- |
| pan | 232581540 232581932 | -64935 | evm.model.Herato0801.107 | hemicentin-1-like isoform X1 |
| pan | 232581540 232581932 | -64935 | evm.model.Herato0801.107 | hemicentin-1-like isoform X1 |
| pan | 232667340 232667968 | 20983 | evm.model.Herato0801.107 | hemicentin-1-like isoform X1 |
| pan | 232667340 232667968 | 20983 | evm.model.Herato0801.107 | hemicentin-1-like isoform X1 |
| pan | 236359663 236359988 | -94873 | evm.model.Herato0801.217 | neural cell adhesion molecule 1-like |
| pan | 236359663 236359988 | -94873 | evm.model.Herato0801.217 | neural cell adhesion molecule 1-like |
| pan | 245499378 245499545 | 23816 | evm.model.Herato0821.60 | piggyBac transposable element-derived protein 3-like |
| pan pan | 245499378 245499545 246654944 246655622 | 23816 -96 | evm.model.Herato0821.60 evm.model.Herato0821.82 | piggyBac transposable element-derived protein 3-likesuccinate dehydrogenase [ubiquinone] cytochrome b small subunit, mitochondrial |
| pan | 253377453 253377634 | 162889 | evm.model.Herato0901.76 | uncharacterized protein LOC113401684 isoform X1 |
| pan | 253377453 253377634 | 162889 | evm.model.Herato0901.76 | uncharacterized protein LOC113401684 isoform X1 |
| pan | 257062073 257062293 | -176441 | evm.model.Herato0901.100 | neurogenic locus protein delta |
| pan | 257062073 257062293 | -176441 | evm.model.Herato0901.100 | neurogenic locus protein delta |
| pan | 257199010 257199726 | -39256 | evm.model.Herato0901.100 | neurogenic locus protein delta |
| pan pan | 257199010 257199726 257694041 257694772 | -39256 4493 | evm.model.Herato0901.100evm.model.Herato0901.104_evm .TU.Herato0901.106 | neurogenic locus protein delta 39S ribosomal protein L19, mitochondrial |
| pan | 266537754 266537987 | 24648 | evm.model.Herato0901.281 | ---NA--- |
| pan | 266537754 266537987 | 24648 | evm.model.Herato0901.281 | ---NA--- |
| pan | 271146968 271147273 | -4865 | evm.model.Herato0901.330 | ---NA--- |

| | | | | |
|---|---|---|---|---|
| pan | 273707743<br>273707954 | -1978 | evm.model.Herato0901.374 | chitinase A-like |
| pan | 274822846<br>274823029 | -15795 | evm.model.Herato0904.26 | ---NA--- |
| pan | 274822846<br>274823029 | -15795 | evm.model.Herato0904.26 | ---NA--- |
| pan | 275196482<br>275196726 | -4908 | evm.model.Herato1001.9 | retinal homeobox protein Rx1-like |
| pan | 275702652<br>275702875 | -15229 | evm.model.Herato1001.23 | ---NA--- |
| pan | 275712168<br>275712672 | -5572 | evm.model.Herato1001.23 | ---NA--- |
| pan | 275712168<br>275712672 | -5572 | evm.model.Herato1001.23 | ---NA--- |
| pan | 276087641<br>276087917 | -8697 | evm.model.Herato1001.34 | putative helix-loop-helix protein hen |
| pan<br>pan<br>pan | 276087641<br>276087917<br>276596775<br>276597297<br>276596775<br>276597297 | -8697<br>433<br>433 | evm.model.Herato1001.34evm.model.Herato1001.47_evm.<br>TU.Herato1001.48evm.model.Herato1001.47_evm.<br>TU.Herato1001.48 | putative helix-loop-helix<br>protein hen inner<br>centromere protein inner<br>centromere protein |
| pan | 281157853<br>281158075 | 8765 | evm.model.Herato1001.118 | laccase-4-like |
| pan | 281157853<br>281158075 | 8765 | evm.model.Herato1001.118 | laccase-4-like |
| pan | 284180137<br>284180409 | -57549 | evm.model.Herato1001.169 | endonuclease-reverse transcriptase HmRTE-e01 |
| pan | 284658235<br>284658504 | -3038 | evm.model.Herato1001.176 | E3 ubiquitin-protein ligase MARCH1-like |
| pan | 284658235<br>284658504 | -3038 | evm.model.Herato1001.176 | E3 ubiquitin-protein ligase MARCH1-like |
| pan | 287085584<br>287085877 | -1121 | evm.model.Herato1003.9 | serine protease inhibitor 2.1-like |
| pan | 287085584<br>287085877 | -1121 | evm.model.Herato1003.9 | serine protease inhibitor 2.1-like |
| pan | 288592377<br>288592734 | 14476 | evm.model.Herato1003.41 | ankyrin repeat domain-containing protein 12 |
| pan | 288592377<br>288592734 | 14476 | evm.model.Herato1003.41 | ankyrin repeat domain-containing protein 12 |
| pan | 295235318<br>295235638 | 2919 | evm.model.Herato1003.150 | oxysterol-binding protein-related protein 1 |
| pan<br>pan | 297147626<br>297147980<br>301366293<br>301366531 | 78407<br>-48471 | evm.model.Herato1003.199 evm.model.Herato1005.77 | uncharacterized protein LOC113393764 isoform X3NADH dehydrogenase [ubiquinone] flavoprotein 2, mitochondrial-like |
| pan | 302206779<br>302207003 | 22552 | evm.model.Herato1005.88 | semaphorin-1A-like |
| pan | 302993939<br>302994268 | -32023 | evm.model.Herato1005.115 | serine/arginine repetitive matrix protein 1-like |
| pan | 302993939<br>302994268 | -32023 | evm.model.Herato1005.115 | serine/arginine repetitive matrix protein 1-like |
| pan | 303103767<br>303104044 | -18521 | evm.model.Herato1005.117 | acetylcholine receptor subunit beta-like 1 |
| pan | 303103767<br>303104044 | -18521 | evm.model.Herato1005.117 | acetylcholine receptor subunit beta-like 1 |
| pan | 303428017<br>303428717 | 33088 | evm.model.Herato1005.222 | ---NA--- |

| | | | | |
|---|---|---|---|---|
| pan | 304422119 304422358 | 20837 | evm.model.Herato1005.168 | septin-7 isoform X1 |
| pan | 309223435 309223683 | 10128 | evm.model.Herato1007.63 | glycine-rich protein DOT1-like |
| pan | 309223435 309223683 | 10128 | evm.model.Herato1007.63 | glycine-rich protein DOT1-like |
| pan | 310179941 310180289 | 67487 | evm.model.Herato1007.87 | venom carboxylesterase-6-like |
| pan | 310179955 310180298 | 67498 | evm.model.Herato1007.87 | venom carboxylesterase-6-like |
| pan | 311175891 311176538 | -36901 | evm.model.Herato1007.97 | ---NA--- |
| pan | 312418674 312419087 | -18926 | evm.model.Herato1007.128 | ---NA--- |
| pan | 312418674 312419087 | -18926 | evm.model.Herato1007.128 | ---NA--- |

| | | | | |
|---|---|---|---|---|
| pan | 313272361 313272581 | -50168 | evm.model.Herato1007.146 | uncharacterized LOC106124082 |
| pan | 313781412 313781624 | 69833 | evm.model.Herato1007.172 | rho GTPase-activating protein conundrum isoform X1 |
| pan pan pan pan | 313781412 313781624 314141458 314141733 314141458 314141733 314142440 314142682 | 69833 22876 22876 23842 | evm.model.Herato1007.172 evm.model.Herato1007.182 evm.model.Herato1007.182 evm.model.Herato1007.182 | rho GTPase-activating protein conundrum isoform X1uncharacterized MFS-type transporter C09D4.1 isoform X2uncharacterized MFS-type transporter C09D4.1 isoform X2uncharacterized MFS-type transporter C09D4.1 isoform X2 |
| pan | 314503736 314504021 | -23520 | evm.model.Herato1007.187 | dynein assembly factor with WDR repeat domains 1 |
| pan | 314503736 314504021 | -23520 | evm.model.Herato1007.187 | dynein assembly factor with WDR repeat domains 1 |
| pan | 314546967 314547366 | 605 | evm.model.Herato1007.188 | lysosomal Pro-X carboxypeptidase |
| pan | 314546967 314547366 | 605 | evm.model.Herato1007.188 | lysosomal Pro-X carboxypeptidase |
| pan | 325564302 325564629 | 3641 | evm.model.Herato1108.86 | phospholipid phosphatase 6 isoform X1 |
| pan | 325564302 325564629 | 3641 | evm.model.Herato1108.86 | phospholipid phosphatase 6 isoform X1 |
| pan | 326448205 326448438 | 9332 | evm.model.Herato1108.114 | ---NA--- |
| pan | 339549358 339550104 | -3478 | evm.model.Herato1108.457 | ZZ-type zinc finger-containing protein 3 |
| pan | 341222830 341223008 | 12736 | evm.model.Herato1108.489 | rab GTPase-activating protein 1-like isoform X1 |
| pan | 341222830 341223008 | 12736 | evm.model.Herato1108.489 | rab GTPase-activating protein 1-like isoform X1 |
| pan | 341348917 341349131 | 23771 | evm.model.Herato1108.491 | serine/arginine repetitive matrix protein 2 |
| pan | 341348917 341349131 | 23771 | evm.model.Herato1108.491 | serine/arginine repetitive matrix protein 2 |
| pan | 343258538 343258739 | -1666 | evm.model.Herato1108.519 | zinc finger protein 62 homolog |
| pan | 344893031 344893228 | -12164 | evm.model.Herato1108.568 | innexin inx7 |
| pan | 344893031 344893228 | -12164 | evm.model.Herato1108.568 | innexin inx7 |
| pan | 346013691 346014002 | -4186 | evm.model.Herato1108.590 | U7 snRNA-associated Sm-like protein LSm11 |
| pan | 346233414 346233600 | 12605 | evm.model.Herato1108.595 | ---NA--- |
| pan | 346233414 346233600 | 12605 | evm.model.Herato1108.595 | ---NA--- |
| pan | 346687145 346687464 | 28480 | evm.model.Herato1108.618 | ---NA--- |
| pan | 346687145 346687464 | 28480 | evm.model.Herato1108.618 | ---NA--- |
| pan pan pan | 352914617 352915112 358550654 358550984 358550654 358550984 | 3246 -46069 -46069 | evm.model.Herato1202.64 evm.model.Herato1202.216 evm.model.Herato1202.216 | protein Cep89 homolog isoform X1sodium- and chloride-dependent transporter XTRP3 isoform X1sodium- and chloride-dependent transporter XTRP3 isoform X1 |
| pan | 369043494 369043800 | -57092 | evm.model.Herato1202.440 | probable cytochrome P450 49a1 |
| pan | 369733928 369734241 | 51091 | evm.model.Herato1202.451 | fibroblast growth factor receptor 3-like isoform X2 |
| pan | 373167408 373167648 | -1642 | evm.model.Herato1202.504 | ESF1 homolog |
| pan pan pan | 373167408 373167648 382411178 382411409 382411178 382411409 | -1642 -523 -523 | evm.model.Herato1202.504 evm.model.Herato1202.756 evm.model.Herato1202.756 | ESF1 homologSWI/SNF-related matrix-associated actin-dependent regulator of chromatin subfamily A containing DEAD/HSWI/SNF-related matrix-associated actin- |

| | | | | | |
|---|---|---|---|---|---|
| | | | | | dependent regulator of chromatin subfamily A containing DEAD/H |
| pan | 383209987 | 383210327 | 55028 | evm.model.Herato1202.776 | mucin-5AC isoform X1 |
| pan pan | 383209987 | 383210327 | 55028 | evm.model.Herato1202.776 | mucin-5AC isoform X1complement component 1 Q |
| pan | 383676938 | 383677257 | 38770 | evm.model.Herato1202.791 | subcomponent-binding protein, mitochondrialcomplement |
| | 383676938 | 383677257 | 38770 | evm.model.Herato1202.791 | component 1 Q subcomponent-binding protein, mitochondrial |
| pan | 385839516 | 385839763 | 24826 | evm.model.Herato1301.6 | protein vestigial |
| pan | 385934868 | 385935360 | 6431 | evm.model.Herato1301.12 | ---NA--- |
| pan | 385934868 | 385935360 | 6431 | evm.model.Herato1301.12 | ---NA--- |
| pan | 385940516 | 385940712 | 931 | evm.model.Herato1301.12 | ---NA--- |
| pan | 385940516 | 385940712 | 931 | evm.model.Herato1301.12 | ---NA--- |
| pan | 387771300 | 387771626 | -6907 | evm.model.Herato1301.33 | protein vein isoform X1 |
| pan | 387771300 | 387771626 | -6907 | evm.model.Herato1301.33 | protein vein isoform X1 |
| pan | 389693316 | 389693952 | -24813 | evm.model.Herato1301.55 | paxillin homolog 1 isoform X1 |
| pan | 389789719 | 389790003 | -20965 | evm.model.Herato1301.57 | elongation of very long chain fatty acids protein 6 |
| pan | 389789719 | 389790003 | -20965 | evm.model.Herato1301.57 | elongation of very long chain fatty acids protein 6 |
| pan | 391289570 | 391289744 | -8017 | evm.model.Herato1301.89 | cuticular protein RR-1 motif 47 precursor |
| pan pan | 391289570 | 391289749 | -8015 | evm.model.Herato1301.89 | cuticular protein RR-1 motif 47 precursorglycine |
| | 397156979 | 397157735 | -5341 | evm.model.Herato1301.242 | dehydrogenase (decarboxylating), mitochondrial |
| pan | 406575959 | 406576181 | 52834 | evm.model.Herato1301.481 | uncharacterized protein LOC117982178 isoform X1 |
| pan | 406575959 | 406576181 | 52834 | evm.model.Herato1301.481 | uncharacterized protein LOC117982178 isoform X1 |
| pan | 411313062 | 411313260 | 71842 | evm.model.Herato1301.557 | neurobeachin isoform X1 |
| pan | 411313062 | 411313260 | 71842 | evm.model.Herato1301.557 | neurobeachin isoform X1 |
| pan | 412573144 | 412573353 | -192819 | evm.model.Herato1301.581 | ubinuclein-1 isoform X3 |
| pan | 413560807 | 413561041 | 10715 | evm.model.Herato1301.589 | ras-related protein Rab-30 |
| pan | 416586255 | 416586611 | -3 | evm.model.Herato1301.628 | AF4/FMR2 family member 1 isoform X1 |
| pan | 416586255 | 416586611 | -3 | evm.model.Herato1301.628 | AF4/FMR2 family member 1 isoform X1 |
| pan | 418157634 | 418158312 | 5454 | evm.model.Herato1301.664 | probable cytochrome P450 301a1, mitochondrial |
| pan | 418157634 | 418158312 | 5454 | evm.model.Herato1301.664 | probable cytochrome P450 301a1, mitochondrial |
| pan | 418949158 | 418949383 | -183997 | evm.model.Herato1301.673 | heat shock protein 21.3 |
| pan | 420033226 | 420034274 | 1081 | evm.model.Herato1301.693 | protein goliath isoform X2 |
| pan | 421265015 | 421265206 | -33041 | evm.model.Herato1301.721 | ---NA--- |
| pan | 427986320 | 427986764 | -72647 | evm.model.Herato1408.25 | esterase FE4-like |
| pan | 427986320 | 427986764 | -72647 | evm.model.Herato1408.25 | esterase FE4-like |
| pan | 428630361 | 428630712 | 128059 | evm.model.Herato1408.27 | ---NA--- |
| pan | 429927387 | 429927859 | -21879 | evm.model.Herato1408.48 | reverse transcriptase |
| pan | 432132149 | 432132737 | 28860 | evm.model.Herato1408.71 | putative phosphatidate phosphatase |
| pan | 435600084 | 435600387 | -80444 | evm.model.Herato1408.165 | Outer dense fiber protein 3-like protein 2 |
| pan | 435600084 | 435600387 | -80444 | evm.model.Herato1408.165 | Outer dense fiber protein 3-like protein 2 |
| pan pan | 442744791 | 442745123 | 57850 | evm.model.Herato1411.54 | transcription factor TFIIIB component B"receptor- |
| pan | 444785017 | 444785197 | 26941 | evm.model.Herato1411.107 | type tyrosine-protein phosphatase kappa isoform |
| | 444785017 | 444785197 | 26941 | evm.model.Herato1411.107 | X1receptor-type tyrosine-protein phosphatase kappa isoform X1 |
| pan | 445226230 | 445226410 | -754 | evm.model.Herato1411.114 | ras-like GTP-binding protein Rho1 isoform X2 |
| pan | 445226230 | 445226410 | -754 | evm.model.Herato1411.114 | ras-like GTP-binding protein Rho1 isoform X2 |
| pan | 448287016 | 448287271 | -35295 | evm.model.Herato1411.184 | endocuticle structural glycoprotein SgAbd-3-like |
| pan | 448287016 | 448287271 | -35295 | evm.model.Herato1411.184 | endocuticle structural glycoprotein SgAbd-3-like |
| pan | 452632455 | 452632671 | -18148 | evm.model.Herato1411.255 | general odorant-binding protein 1-like |
| pan | 452632455 | 452632671 | -18148 | evm.model.Herato1411.255 | general odorant-binding protein 1-like |
| pan | 453146748 | 453147200 | 72512 | evm.model.Herato1502.1 | 60S ribosomal protein L38 |

| | | | | |
|---|---|---|---|---|
| pan | 453146748 453147200 | 72512 | evm.model.Herato1502.1 | 60S ribosomal protein L38 |
| pan | 454345096 454345377 | -9086 | evm.model.Herato1505.20 | peptidyl-tRNA hydrolase ICT1, mitochondrial |
| pan pan pan | 454345096 454345377 460350719 460351010 460350724 460350955 | -9086 -19207 -19232 | evm.model.Herato1505.20evm.model.Herato1505.143_evm peptidyl-tRNA hydrolase ICT1, mitochondrial .TU.Herato1505.146evm.model.Herato1505.143_evm nudC domain-containing protein 3 .TU.Herato1505.146 | nudC domain-containing protein 3 |
| pan | 467737096 467737578 | -87459 | evm.model.Herato1507.78 | nudix hydrolase 8-like |
| pan | 467737096 467737578 | -87459 | evm.model.Herato1507.78 | nudix hydrolase 8-like |
| pan | 468000016 468000246 | -6697 | evm.model.Herato1507.89 | uncharacterized protein LOC113400528 |
| pan pan pan pan | 468000016 468000246 471443633 471443879 471443633 471443879 471648976 471649233 | -6697 -12516 -12516 -28094 | evm.model.Herato1507.89 evm.model.Herato1507.151 evm.model.Herato1507.151 evm.model.Herato1507.156 | uncharacterized protein LOC113400528evolutionarily conserved signaling intermediate in Toll pathway, mitochondrialevolutionarily conserved signaling intermediate in Toll pathway, mitochondrialbrain-specific angiogenesis inhibitor 1-associated protein 2 isoform X3 |
| pan | 472207958 472208231 | -55977 | evm.model.Herato1507.163 | acidic repeat-containing protein-like |
| pan | 474264556 474264774 | -67192 | evm.model.Herato1507.215 | Regulator of G-protein signaling 7 |
| pan | 476732908 476733209 | 5117 | evm.model.Herato1507.249 | caskin-2 isoform X1 |
| pan | 476732908 476733209 | 5117 | evm.model.Herato1507.249 | caskin-2 isoform X1 |
| pan | 478102737 478103098 | 37261 | evm.model.Herato1507.273 | 2-hydroxyacyl-CoA lyase 1 isoform X1 |
| pan | 487914187 487915309 | -26769 | evm.model.Herato1524.18 | sodium-independent sulfate anion transporter-like |
| pan | 488333710 488333971 | -859 | evm.model.Herato1601.8 | serine/arginine repetitive matrix protein 1 |
| pan | 488333710 488333971 | -859 | evm.model.Herato1601.8 | serine/arginine repetitive matrix protein 1 |
| pan | 489240124 489240453 | 11703 | evm.model.Herato1601.32 | piggyBac transposable element-derived protein 4-like |
| pan pan | 489240124 489240453 490499823 490500099 | 11703 -120817 | evm.model.Herato1601.32 evm.model.Herato1601.49 | piggyBac transposable element-derived protein 4-likeacyl-CoA:lysophosphatidylglycerol acyltransferase 1like |
| pan | 492925936 492926320 | -19375 | evm.model.Herato1601.58 | uncharacterized LOC106121622 precursor |
| pan | 492925936 492926320 | -19375 | evm.model.Herato1601.58 | uncharacterized LOC106121622 precursor |
| pan | 497995591 497995758 | 39966 | evm.model.Herato1601.108 | spectrin beta chain isoform X1 |
| pan | 497995591 497995758 | 39966 | evm.model.Herato1601.108 | spectrin beta chain isoform X1 |
| pan | 502418487 502418838 | -25421 | evm.model.Herato1603.127 | ---NA--- |
| pan | 502874422 502874616 | -5212 | evm.model.Herato1603.47 | CWF19-like protein 2 homolog |
| pan | 503206138 503206341 | -32230 | evm.model.Herato1603.58 | mucin-5AC isoform X1 |
| pan | 503206138 503206341 | -32230 | evm.model.Herato1603.58 | mucin-5AC isoform X1 |
| pan | 504937412 504937709 | 5489 | evm.model.Herato1603.84 | uncharacterized protein LOC113399111 |
| pan | 504937412 504937709 | 5489 | evm.model.Herato1603.84 | uncharacterized protein LOC113399111 |
| pan | 505779592 505779821 | -93039 | evm.model.Herato1603.89 | apoptosis-stimulating of p53 protein 1 isoform X4 |
| pan | 505779592 505779821 | -93039 | evm.model.Herato1603.89 | apoptosis-stimulating of p53 protein 1 isoform X4 |
| pan | 514678315 514678662 | 15979 | evm.model.Herato1605.121 | ---NA--- |
| pan | 514678320 514678599 | 15950 | evm.model.Herato1605.121 | ---NA--- |
| pan | 516114995 516115358 | 27483 | evm.model.Herato1605.160 | helix-loop-helix protein delilah-like |

| | | | | |
|---|---|---|---|---|
| pan | 516114995 516115358 | 27483 | evm.model.Herato1605.160 | helix-loop-helix protein delilah-like |
| pan | 521500041 521500424 | 92188 | evm.model.Herato1701.63 | nephrin-like isoform X1 |
| pan | 532953168 532953375 | 15094 | evm.model.Herato1701.306 | cytoplasmic dynein 2 heavy chain 1-like |
| pan | 532953168 532953375 | 15094 | evm.model.Herato1701.306 | cytoplasmic dynein 2 heavy chain 1-like |
| pan | 533257935 533258195 | 24357 | evm.model.Herato1701.315 | SUN domain-containing protein 3-like |
| pan | 533257935 533258195 | 24357 | evm.model.Herato1701.315 | SUN domain-containing protein 3-like |
| pan | 533584122 533584420 | -930 | evm.model.Herato1701.334 | G1/S-specific cyclin-D3 |
| pan | 533584122 533584420 | -930 | evm.model.Herato1701.334 | G1/S-specific cyclin-D3 |
| pan | 544085362 544085635 | -29870 | evm.model.Herato1705.13 | fat-like cadherin-related tumor suppressor homolog |
| pan | 544085362 544085635 | -29870 | evm.model.Herato1705.13 | fat-like cadherin-related tumor suppressor homolog |

| | | | | |
|---|---|---|---|---|
| pan | 545973766 545973960 | -17038 | evm.model.Herato1705.76 | translocon-associated protein subunit delta |
| pan | 545973766 545973960 | -17038 | evm.model.Herato1705.76 | translocon-associated protein subunit delta |
| pan | 551774678 551774959 | -5605 | evm.model.Herato1712.7 | unnamed protein product |
| pan pan | 551774678 551774959<br>552842888 552843379 | -5605<br>-12378 | evm.model.Herato1712.7<br>evm.model.Herato1801.20 | unnamed protein productRNA-directed DNA polymerase from mobile element jockey |
| pan | 553014251 553014614 | 39352 | evm.model.Herato1801.17 | basic juvenile hormone-suppressible protein 1-like |
| pan pan | 556975312 556975615<br>557316011 557316444 | -66615<br>39819 | evm.model.Herato1801.97<br>evm.model.Herato1801.108 | Protein YIF1B-Bleucine zipper putative tumor suppressor 2 homolog isoform X1 |
| pan pan<br>pan | 557495185 557495624<br>557534754 557535032<br>557534754 557535032 | -11461<br>19619<br>19619 | evm.model.Herato1801.111<br>novel_exon_214 novel_exon_214 | nucleolin-likeRetrovirus-related Pol polyprotein from type-2 retrotransposable element R2DM; EndonucleaseRetrovirus-related Pol polyprotein from type-2 retrotransposable element R2DM; Endonuclease |
| pan | 559422520 559422705 | -10258 | evm.model.Herato1801.147 | hybrid signal transduction histidine kinase D-like |
| pan | 559422520 559422705 | -10258 | evm.model.Herato1801.147 | hybrid signal transduction histidine kinase D-like |
| pan | 563474237 563475462 | -19761 | evm.model.Herato1805.41 | hexokinase type 2 isoform X1 |
| pan | 563915416 563915916 | 48755 | evm.model.Herato1805.55 | protein couch potato isoform X1 |
| pan | 566064268 566064449 | -64750 | evm.model.Herato1805.81 | toll-like receptor Tollo |
| pan | 566064268 566064449 | -64750 | evm.model.Herato1805.81 | toll-like receptor Tollo |
| pan | 567453569 567453829 | 99589 | evm.model.Herato1805.88 | fatty acid synthase-like |
| pan | 567453569 567453829 | 99589 | evm.model.Herato1805.88 | fatty acid synthase-like |
| pan | 567992230 567992535 | -1592 | evm.model.Herato1805.95 | exportin-4-like |
| pan | 572718227 572718506 | -28576 | evm.model.Herato1805.160 | muscarinic acetylcholine receptor DM1 |
| pan | 573007484 573007696 | -28616 | evm.model.Herato1805.166 | uncharacterized protein LOC116769663 isoform X2 |
| pan | 573007484 573007704 | -28620 | evm.model.Herato1805.166 | uncharacterized protein LOC116769663 isoform X2 |
| pan | 573077682 573078157 | -1946 | evm.model.Herato1805.169 | tubulin-folding cofactor B |
| pan | 573077682 573078157 | -1946 | evm.model.Herato1805.169 | tubulin-folding cofactor B |
| pan | 575215299 575215478 | -21479 | evm.model.Herato1805.219 | organic cation transporter protein-like |
| pan | 576294146 576294579 | -37298 | evm.model.Herato1805.234 | ---NA--- |
| pan | 576294146 576294579 | -37298 | evm.model.Herato1805.234 | ---NA--- |
| pan | 576939877 576940254 | -13102 | evm.model.Herato1805.247 | inhibin beta B chain |
| pan | 582743269 582743661 | 9255 | evm.model.Herato1807.37 | catenin delta-2 isoform X1 |
| pan | 582743269 582743661 | 9255 | evm.model.Herato1807.37 | catenin delta-2 isoform X1 |
| pan | 582769636 582769834 | 35525 | evm.model.Herato1807.37 | catenin delta-2 isoform X1 |
| pan | 582945825 582946065 | 12939 | evm.model.Herato1807.38 | hypothetical protein evm_001344 |
| pan | 583499973 583500202 | -7586 | evm.model.Herato1807.41 | latrophilin Cirl isoform X3 |
| pan | 585464983 585465215 | -7368 | evm.model.Herato1807.68 | zinc finger protein rotund-like |
| pan | 586396145 586396443 | 4417 | evm.model.Herato1807.87 | transcriptional regulator ATRX-like |
| pan | 586396145 586396443 | 4417 | evm.model.Herato1807.87 | transcriptional regulator ATRX-like |
| pan | 587385242 587385518 | -2843 | evm.model.Herato1807.114 | Transcription factor Adf-1 |
| pan | 587385242 587385518 | -2843 | evm.model.Herato1807.114 | Transcription factor Adf-1 |
| pan | 589803073 589803383 | 34105 | evm.model.Herato1901.117 | ---NA--- |
| pan | 592706260 592706664 | -33229 | evm.model.Herato1901.66 | mycosubtilin synthase subunit C |
| pan | 592706260 592706664 | -33229 | evm.model.Herato1901.66 | mycosubtilin synthase subunit C |
| pan | 594767787 594768428 | 243 | evm.model.Herato1901.115 | uncharacterized protein LOC112056053 |
| pan | 602872903 602873510 | 4305 | evm.model.Herato1904.140 | nucleolar protein 12 |
| pan | 602872903 602873510 | 4305 | evm.model.Herato1904.140 | nucleolar protein 12 |
| pan | 602889111 602889562 | -11825 | evm.model.Herato1904.140 | nucleolar protein 12 |
| pan | 602889111 602889562 | -11825 | evm.model.Herato1904.140 | nucleolar protein 12 |
| pan | 604029667 604030052 | 5209 | evm.model.Herato1904.163 | ---NA--- |
| pan | 604029667 604030052 | 5209 | evm.model.Herato1904.163 | ---NA--- |
| pan | 608477896 608478167 | 1112 | evm.model.Herato1904.263 | Outer dense fiber protein 3-like protein 2 |

| | | | | |
|---|---|---|---|---|
| pan | 609365811<br>609366051 | -1836 | evm.model.Herato1904.269 | nose resistant to fluoxetine protein 6-like |
| pan | 609365811<br>609366051 | -1836 | evm.model.Herato1904.269 | nose resistant to fluoxetine protein 6-like |
| pan<br>pan<br>pan<br>pan | 610686415<br>610686692<br>610760654<br>610760855<br>610773751<br>610774194<br>610773751<br>610774194 | -4155<br>-17654<br>-30872<br>-30872 | evm.model.Herato1905.5 evm.model.Herato1905.8<br>evm.model.Herato1905.8 evm.model.Herato1905.8 | 4ATP-binding cassette sub-family G member 4-like F2 cell-surface antigen heavy chain-like isoform X2ATP-binding cassette sub-family G member 4-like isoform X2ATP-binding cassette sub-family G member 4-like isoform X2 |
| pan | 612406397<br>612406638 | 77188 | evm.model.Herato1906.21 | headcase protein |
| pan<br>pan<br>pan | 612406397<br>612406638<br>614361975<br>614362189<br>614361975<br>614362189 | 77188<br>-12546<br>-12546 | evm.model.Herato1906.21 evm.model.Herato1908.10<br>evm.model.Herato1908.10 | headcase proteinphosphoinositide 3-kinase adapter protein 1 isoform<br>X2phosphoinositide 3-kinase adapter protein 1 isoform<br>X2 |
| pan | 618093638<br>618093802 | -6713 | evm.model.Herato1908.94 | 39S ribosomal protein L48, mitochondrial |
| pan | 618093638<br>618093802 | -6713 | evm.model.Herato1908.94 | 39S ribosomal protein L48, mitochondrial |
| pan | 618959143<br>618959337 | -1200 | evm.model.Herato1910.19 | RNA polymerase II transcriptional coactivator |
| pan | 620145575<br>620145989 | 26423 | evm.model.Herato1910.61 | protein turtle isoform X1 |
| pan | 621316080<br>621316517 | 17658 | evm.model.Herato1910.102 | dnaJ homolog subfamily C member 10-like |
| pan | 621316080<br>621316517 | 17658 | evm.model.Herato1910.102 | dnaJ homolog subfamily C member 10-like |
| pan | 621731732<br>621732089 | -60280 | evm.model.Herato1910.108 | uncharacterized protein LOC117984558 |
| pan | 624295389<br>624295635 | 9449 | evm.model.Herato1910.152 | etoposide-induced protein 2.4 homolog |
| pan | 624410084<br>624410619 | -81104 | evm.model.Herato1910.153 | ---NA--- |
| pan | 628814865<br>628815964 | 4962 | evm.model.Herato1910.225 | protein borderless |
| pan | 630310190<br>630310603 | -214983 | evm.model.Herato2001.17 | zinc finger protein Xfin-like |
| pan<br>pan<br>pan<br>pan<br>pan | 630310190<br>630310603<br>632911060<br>632911331<br>632911060<br>632911331<br>641775817<br>641776153<br>641775817<br>641776153 | -214983<br>885<br>885<br>-353<br>-353 | evm.model.Herato2001.17evm.model.Herato2001.72_evm.<br>TU.Herato2001.73evm.model.Herato2001.72_evm.<br>TU.Herato2001.73 evm.model.Herato2001.326<br>evm.model.Herato2001.326 | zinc finger protein Xfin-like<br>THO complex subunit 6<br>THO complex subunit 6probable cytosolic iron-sulfur protein assembly protein<br>Ciao1 isoform X1probable cytosolic iron-sulfur protein assembly protein<br>Ciao1 isoform X1 |
| pan | 643452007<br>643452427 | 173 | evm.model.Herato2001.383 | RNA-binding protein Musashi-like Rbp6 |
| pan | 643452007<br>643452427 | 173 | evm.model.Herato2001.383 | RNA-binding protein Musashi-like Rbp6 |
| pan | 644386709<br>644387075 | -46274 | evm.model.Herato2001.394 | uncharacterized protein LOC113400031 |

| | | | | |
|---|---|---|---|---|
| pan | 644386709 644387075 | -46274 | evm.model.Herato2001.394 | uncharacterized protein LOC113400031 |
| pan | 644496038 644496329 | 63017 | evm.model.Herato2001.394 | uncharacterized protein LOC113400031 |
| pan | 652409589 652409827 | 25421 | evm.model.Herato2001.582 | putative B-cell lymphoma 3-encoded protein |
| pan pan pan pan pan | 652409589 652409827 664820583 664820820 664820583 664820820 671750430 671750741 671750430 671750741 | 25421 44929 44929 21878 21878 | evm.model.Herato2001.582 evm.model.Herato2101.108 evm.model.Herato2101.108 evm.model.Herato2101.260 evm.model.Herato2101.260 | putative B-cell lymphoma 3-encoded proteinalpha-1,3-mannosyl-glycoprotein 4-beta-N- acetylglucosaminyltransferase A-likealpha-1,3-mannosyl-glycoprotein 4-beta-Nacetylglucosaminyltransferase A-likecoiled-coil domain-containing protein AGAP005037 isoform X1coiled-coil domain-containing protein AGAP005037 isoform X1 |
| pan | 682848573 682848799 | -4907 | evm.model.Herato2101.451 | otoancorin like protein |

| | Fst | | | | | sweep | | | |
|---|---|---|---|---|---|---|---|---|---|
| | notabilis vs | demophoon | demophoon | demophoon | demophoon | notabilis | etylus hydara | demophoon | |
| total | 464 | 720 | 1286 | 1315 | 2725 | 492 | 494 | 746 | 746 |
| 0-0.1 | 410(88,4%) | 626(87%) | 930(72,4%) | 858(65,3%) | 324(11,9%) | 491(99,8%) | 492(99,6%) | 742(99,5%) | 738(99%) |
| 0.1-0.2 | 28(6,1%) | 41(5,7%) | 181(14,1%)** | 213(16,2%)* | 737(27,1%) | 1(0,3%) | 0(0%) | 0(0%) | 6(0,9%)** |
| 0.2-0.3 | 8(1,8%) | 26(3,7%)*** | 78(6,1%)*** | 96(7,4%)*** | 756(27,8%) | 0(0%) | 2(0,5%) | 3(0,5%) | 0(0%) |
| 0.3-0.4 | 6(1,3%) | 11(1,6%) | 52(4,1%)*** | 70(5,4%)*** | 494(18,2%) | 0(0%) | 0(0%) | 1(0,2%) | 2(0,3%) |
| 0.4-0.5 | 6(1,3%)* | 7(1%) | 16(1,3%) | 32(2,5%)** | 234(8,6%) | 0(0%) | 0(0%) | 0(0%) | 0(0%) |
| 0.5-0.6 | 3(0,7%) | 3(0,5%) | 19(1,5%)*** | 22(1,7%)*** | 93(3,5%) | 0(0%) | 0(0%) | 0(0%) | 0(0%) |
| 0.6-0.7 | 2(0,5%) | 5(0,7%)** | 3(0,3%) | 13(1%)** | 57(2,1%) | 0(0%) | 0(0%) | 0(0%) | 0(0%) |
| 0.7-0.8 | 1(0,3%) | 1(0,2%) | 3(0,3%) | 5(0,4%) | 24(0,9%) | 0(0%) | 0(0%) | 0(0%) | 0(0%) |
| 0.8-0.9 | 0(0%) | 0(0%) | 3(0,3%) | 2(0,2%) | 4(0,2%) | 0(0%) | 0(0%) | 0(0%) | 0(0%) |
| 0.9-1 | 0(0%) | 0(0%) | 1(0,1%) | 4(0,4%)** | 2(0,1%) | 0(0%) | 0(0%) | 0(0%) | 0(0%) |

TableS6: Custom binomial test results, the significant one are in bold

| Geo. morph | GO category | GO term | GO term Name | gene name | Gene name |
|---|---|---|---|---|---|
| demophoo | Trascription | GO:0003676 | nucleic acid binding | evm.model.Herato0413.1 | Retrovirus-related Pol polyprotein from transposon opus-like |
| | | GO:000372 | RNA binding | evm.model.Herato0821.65 | polyribonucleotide nucleotidyltransferase 1 |
| notabilis vs | | GO:0003676 | nucleic acid binding | evm.model.Herato0413.1 | Retrovirus-related Pol polyprotein from transposon opus-like Protein |
| | | GO:0003676 | nucleic acid binding | evm.model.Herato1411.45 | angiogenic factor with G patch and FHA domains 1 isoform X2 |
| | Trascription | GO:0003682 | chromatin binding | evm.model.Herato0101.589 | chromodomain-helicase-DNA-binding protein 7 |
| | | GO:0003677 | DNA binding | evm.model.Herato1801.64 | optix |
| | | GO:0003723 | RNA binding | evm.model.Herato0821.65 | polyribonucleotide nucleotidyltransferase 1 |

TableS7: gene in Fst that are related to Transcription factor or chromatin remodeling

| | | | | | | | | | | | |
|---|---|---|---|---|---|---|---|---|---|---|---|
| H. e. lemophoo | pan pan | 10555853 | 10556334 | LTR/Gypsy,RC | 306 | 63.49 | -6031 | ID=evm.TU.Herato0101.165 spherulin-2A-like | -NA- | response to stress; function in insects | No |
| | pan | 249036381 | 249036811 | NA | NA | NA | -3903 | ID=novel_exon_96 --NA-- | -NA- | Not available. Unknown similarity but with | |
| | | 58476471 | 58477381 | RC/Helitron | 708 | 77.72 | -14580 | ID=evm.TU.Herato0215.147 hypothetical protein | -NA- | no experimentally | Unknown |
| H. e. hydara | pan pan | 168523255 | 168523563 | DNA/hAT | 171 | 55.34 | -2682 | ID=evm.TU.Herato0606.96 protein spindle-F | -NA- | Involved in mitotic spindle formation during | No |
| | pan pan | 209944819 | 209945370 | RC/Helitron | 115 | 20.83 | 7024 | ID=evm.TU.Herato0701.517 tether containing UBX domain for GLUT4 | -NA- | Likely involved in vesicle trafficking, partic | No |
| | pan pan | 427832052 | 427832531 | LTR/Gypsy | 476 | 99.17 | 30888 | ID=evm.TU.Herato1408.23 cholinesterase 2-like | GO:0052689carboxylic ester hydrolase activity | Breaks down acetylcholine; involved in ne | No |
| | pan pan | 427833038 | 427834545 | LTR/Gypsy | 1508 | 100.00 | 29388 | ID=evm.TU.Herato1408.23 cholinesterase 2-like | GO:0052689carboxylic ester hydrolase activity | Same as above. | No |
| | | 429316871 | 429317232 | NA | NA | NA | 151848 | ID=evm.TU.Herato1408.28 protein D2-like | -NA- | Potentially involved in cellular structural or | No |
| | | 465267855 | 465268537 | RC/Helitron | 683 | 100.00 | 11691 | ID=evm.TU.Herato1507.46 uncharacterized protein | -NA- | Function unknown; predicted based on se | Unknown |
| | | 473551006 | 473551957 | LTR/Gypsy,LTR | 263 | 27.63 | -39669 | ID=evm.TU.Herato1507.195 Tigger transposable element-derived protein 6 | GO:0110165cellular anatomical entity | Derived from transposable elements; may | No |
| | | 515560289 | 515561120 | LINE/RTE-RTE | 613 | 73.68 | | | | Associated with mRNA processing and re | Yes |
| | | | | | | | 9871 | ID=evm.TU.Herato1605.141 symplekin | GO:0006397mRNA processing | | |
| H. e. favorinus | pan pan | 184738636 | 184738915 | NA | NA | NA | -17529 | ID=evm.TU.Herato0606.437 serine/threonine-protein kinase TEL1-like | 4 catalytic activity | telomere maintenance. | Yes |
| | pan pan | 211464003 | 211464602 | LINE/RTE-BovB | 434 | 72.33 | 10597 | ID=evm.TU.Herato0701.601 activating transcription factor of chaperone | -NA- | regulating protein folding and | Yes |
| | pan pan | 312636532 | 312636895 | NA | NA | NA | -3704 | ID=evm.TU.Herato1007.137 BLOC-1-related complex subunit 8 homolog | 5 lysosomal membrane | chaperone pigmentation; potentially | No |
| | pan pan | 316465727 | 316466799 | Simple_repeat | 32 | 29.82 | -23062 | ID=evm.TU.Herato1101.1 retinal guanylyl cyclase 1 | 0;GO:0009 | affects insect phototransduction; may | No |
| | pan pan | 38281574 | 38282344 | LINE/L2 | 86 | 11.15 | activity;phosphorus-oxygen lyase | | | play a role in regulation. | Yes |
| | pan | 386083877 | 386085206 | LINE/I-Jockey | 719 | 54.06 | -21506 | ID=evm.TU.Herato0203.2 SWI5-dependent HO expression protein 3-like | -NA- | function and stability. | No |
| | | 493899738 | 493900570 | NA | NA | NA | 9991 | ID=evm.TU.Herato1301.14 neurofilament heavy polypeptide isoform X1 | -NA- | role in muscle attachment and | No |
| | | 51490214 | 51490901 | DNA/TcMar-Ma | 590 | 85.76 | -22414 | ID=evm.TU.Herato1601.69 connectin-like | -NA- | cleavage; may influence insect | No |
| | | 557245224 | 557245883 | LINE/RTE-BovB | 408 | 61.82 | 9517 | ID=evm.TU.Herato0211.114 nardilysin-like isoform X1 | 2;GO:0046 ion | processing, important for gene | Yes |
| | | 613801026 | 613801637 | LINE/L2 | 612 | 100.00 | binding;proteolysis | | | biosynthesis, crucial for insect | No |
| | | 671189111 | 671189998 | LINE/R1 | 888 | 100.00 | 1248 | ID=evm.TU.Herato1801.105 probable oligoribonuclease | 5;GO:0003 | late embryogenesis; critical in | No |
| | | | | | | | activity;nucleic acid | | | insect | |
| | | | | | | | 34091 | ID=evm.TU.Herato1907.1 juvenile hormone acid O-methyltransferase | 0 transferase activity | | |
| | | | | | | | 18762 | ID=evm.TU.Herato2101.243 late embryogenesis abundant protein, group 3-like | -NA- | | |
| H. e. etylus | pan pan | 161337939 | 161338369 | RC/Helitron,SIN | 247 | 57.31 | -74577 | ID=evm.TU.Herato0601.104 protein timeless homolog | -NA- | developmental timing in insects. | Yes |
| | pan | 304102616 | 304103249 | LTR | 158 | 24.92 | -30917 | ID=evm.TU.Herato1005.156 proclotting enzyme | 2;GO:0006 | healing. | No |
| | | 57328840 | 57329056 | DNA/TcMar-Ma | 178 | 82.03 | activity;proteolysis | | | involved in detoxification and metabolism | No |
| | | | | | | | -5090 | ID=evm.TU.Herato0214.50 cytochrome P450 4C1-like | 7;GO:0005 binding;oxidoreductase activity, | | |
| H. e. notabilis | pan pan | 135554519 | 135555807 | DNA,DNA/TcM | 449 | 34.83 | 5357 | ID=evm.TU.Herato0503.112 carboxylesterase 1E | 7 hydrolase activity | bonds; involved in insect | No |
| | pan pan | 143166040 | 143166517 | Simple_repeat | 50 | 98.75 | -16745 | ID=evm.TU.Herato0503.242 integrin beta-nu | 0;GO:0007 membrane;cell adhesion | metabolism, tissue development | No |
| | pan | 178810204 | 178811044 | RC/Helitron,SIN | 655 | 77.88 | 13848 | ID=evm.TU.Herato0606.269 enzyme-like uncha | 0;GO:0004 activity;metallopeptidase | and maintenance in vertebrates, but | No |
| | | 422872671 | 422873572 | RC/Helitron,SIN | 285 | 31.60 | -10248 | ID=evm.TU.Herato1301.744 LOC113398135 unc | -NA- | may play a role in ion sequence. | Unknown |
| | | 47518084 | 47518972 | RC/Helitron,SIN | 476 | 53.54 | 17959 | ID=evm.TU.Herato0209.49 LOC113395223 isofor activity | 0;GO:0016 membrane;oxidoreductase | validation. | Unknown |
| e. chestertoni | pan pan | 17274167 | 17274698 | LINE/I-Jockey | 56 | 10.52 | -43401 | ID=evm.TU.Herato0101.288 insulin-like growth factor-binding protein complex acid la | 0;GO:0071 | potentially impacting insect | No |
| | pan pan | 228367224 | 228367882 | DNA | 171 | 25.95 | membrane;cell periphery;locomotion -6583 | ID=evm.TU.Herato0801.48 probable | | size and important for insect | No |
| | pan pan | 257402467 | 257403176 | RC/Helitron,SIN | 165 | 23.24 | | maleylacetoacetate isomerase 2 isoform X1 | -NA- | metabolism. structure or | No |
| | pan pan | 309390083 | 309390757 | DNA/PiggyBac, | 354 | 52.44 | -97395 | ID=evm.TU.Herato0901.102 nuclear anchorage protein 1 | 5;GO:0016 nuclear | positioning. sequence. | Unknown |
| | pan pan | 343811050 | 343812296 | LINE/RTE-BovB | 665 | 53.33 | envelope;membrane | | | potential role in insect neurobiology or | No |
| | | 458253784 | 458254266 | LTR,RC/Helitro | 456 | 94.41 | -11617 | ID=evm.TU.Herato1007.73 uncharacterized protein LOC113401061 | -NA- | meiosis in insects, including | Yes |
| | | 551694723 | 551695408 | LINE/I-Jockey | 113 | 16.47 | -77245 | ID=evm.TU.Herato1108.527 prion-like-(Q/N-rich) domain-bearing protein 25 | -NA- | Heliconius. | Unknown |
| | | 555630700 | 555631219 | LINE/I-Jockey | 518 | 99.62 | 223739 | ID=evm.TU.Herato1505.85 cortex | 7;GO:0010 complex | Not available. | No |
| | | 578264290 | 578264565 | LTR/Gypsy | 302 | 10.94 | binding;ubiquitin-protein | | | for energy production in insects. | Yes |
| | | 592939160 | 592939536 | LINE/L2,Simple | 127 | 33.69 | 6048 | ID=evm.TU.Herato1712.2 ---NA--- | -NA- | regulation; zinc finger proteins are key in turnover; may regulate protein stability in | Yes |
| | | | | | | | -10694 | ID=evm.TU.Herato1801.85 short/branched chain specific acyl-CoA dehydrogenase, m | 5;GO:0050 | | |
| | | | | | | | activity;flavin adenine dinucleotide | | | | |
| | | | | | | | -1943 | ID=evm.TU.Herato1805.277 zinc finger protein 567-like | -NA- | | |
| | | | | | | | 3336 | ID=evm.TU.Herato1901.70 ubiquitin-like domain-containing CTD phosphatase 1 | 8;GO:0008 | | |
| | | | | | | | activity;metabolic process | | | | |

Table S8:TEs content of NON-homologous unique ATAC-peaks
